# Expanding the Repertoire of Photoswitchable Unnatural Amino Acids for Enzyme Engineering

**DOI:** 10.1101/2025.04.17.649278

**Authors:** Caroline Hiefinger, Michela Marcon, Verena Pape, Guillem Casadevall, Ranit Lahmy, Julian Nazet, Michael Bartl, Astrid Bruckmann, Sílvia Osuna, Burkhard König, Andrea Hupfeld

**Affiliations:** Institute of Biophysics and Physical Biochemistry and Regensburg Center for Biochemistry, University of Regensburg, Universitätsstraße 31, D-93053 Regensburg (Germany); Institute of Organic Chemistry, University of Regensburg, Universitätsstraße 31, D-93053 Regensburg (Germany); Institut de Química Computacional i Catàlisi and Departament de Química, Universitat de Girona, c/ Maria Aurèlia Capmany 69, 17003 Girona (Spain); Institute of Biochemistry, Genetics and Microbiology, University of Regensburg, Universitatsstrasse 31, D-93053 Regensburg (Germany); ICREA, Pg. Lluís Companys 23, 08010 Barcelona (Spain)

**Keywords:** enzyme catalysis, diazo compounds, photocontrol, spiro compounds, unnatural amino acids

## Abstract

Photoswitchable unnatural amino acids (psUAAs) play a crucial role in the engineering of light-sensitivity in enzymes, which holds significant promise for diverse applications such as biotherapy and biocatalysis. Besides near-quantitative photoconversion, the success and expediency of a psUAA for a certain application is defined by its interaction potential with the enzyme, its thermal stability and its effective wavelength of irradiation. To establish high versatility in the current repertoire, we have designed and synthesized six psUAAs based on azobenzene, arylazopyrazole, arylazothiazole, hemithioindigo and spiropyran photoswitches. The resulting psUAAs exhibit an enhanced interaction potential within an enzyme owing to their capacity for hydrogen bonding, ionic interactions, and metal ion coordination. Moreover, we observed diverse photochemical behaviors among the psUAAs, with four of them reversibly switching between the isomers with purely visible light. Notably, we identified orthogonal aminoacyl-tRNA synthetases that facilitate the incorporation of five of the six psUAAs co-translationally and computationally analyzed the synthetase-psUAA interactions. Finally, we evaluated the photochemical behavior of the five psUAAs within an enzymatic model and tested the photocontrol of catalysis confirming their diversity. Ultimately, our findings significantly expanded the repertoire of psUAAs and demonstrated their feasibility for enzyme engineering studies.

## Introduction

Reversible spatiotemporal control by light is gaining increasing significance in enzyme engineering applications.^[1]^ In this regard, protein engineering with photoswitchable unnatural amino acids (psUAAs) facilitated by amber suppression using orthogonal aminoacyl-tRNA synthetase (aaRS) / tRNA pairs, has emerged as a powerful method.^[2]^ Its key advantage is the genetic encoding of light-sensitivity whilst minimally altering the enzyme via a single amino acid exchange. Generally, psUAAs comprise a synthetic photoswitch^[3–5]^ that can adopt two configurations, e.g., *E* and *Z*, which are approximating 100% of the more stable configuration, mostly *E*, in thermal equilibrium (TEQ). Photocontrol is achieved by irradiation with a specific wavelength *λ* that shifts the equilibrium towards the less stable configuration, thereby establishing a so-called photostationary state (PSS^λ^) with a defined isomer composition. Moreover, the higher the fraction of the less stable isomer at the PSS^λ^, and consequently the more quantitative the photoconversion, the greater the potential for efficient photocontrol. Ultimately, reversibility is facilitated by either the photoinduced reestablishment of a PSS^λ^ similar to TEQ or relaxation of the less stable isomer to the thermal equilibrium TEQ^post^.

Hitherto, photocontrol with psUAAs has mainly been the result of allosteric conformational changes in the enzyme induced by the isomer switch.^[6–8]^ Thus, besides the efficiency of the photoswitch, the successful engineering of photocontrol in enzymes requires a well-defined communication between the incorporated psUAA and the orthosteric site leading to functional differences initiated by the photoinduced configurational changes.^[9]^ This communication is determined by both the position of incorporation and the interaction potential of the psUAA with the adjacent enzyme environment. Considering the vast properties of enzyme targets and incorporation sites, the availability of a broad selection of unpolar and polar interactions, that may even potentially allow the psUAA to take part in catalysis, would significantly increase the success rate of photocontrol engineering with psUAAs.

Furthermore, the diversity of applications for photocontrolled enzymes including biotherapeutic strategies and industrial biocatalysis necessitates a repertoire of psUAAs that is adapted to the specific demands of each approach. This entails psUAAs, which i) either maintain their configuration or thermally relax to their initial state after irradiation stops, and ii) respond to various wavelengths including visible light, which confers reduced toxicity and higher penetration depths.

Notably, the current repertoire of psUAAs (**Figure S1**)^[10]^ only covers limited diversity in photoswitch efficiencies, interaction potentials, thermal stabilities and effective wavelengths of irradiation. Most of these psUAAs comprise an azobenzene photoswitch, starting with unsubstituted phenylalanine-4′-azobenzene (**AzoF**; **Figure 1**).^[11,12]^ Various substitutions allowed for crosslinking **AzoF** with cysteins in a protein via click chemistry,^[13,14]^ and for switching using exclusively visible light.^[8,15]^ However, these changes came in part at the cost of less quantitative photoconversions. A wider range of diversity can be explored by using different scaffolds from the vast variety of existing photoswitches.^[3,4]^ Most recently, two diazocine-based psUAAs were introduced, which similarly reacted solely to visible light but lost switching efficiency.^[16]^ Two further studies reported the synthesis and incorporation of an arylazopyrazole-based psUAA (AapF) with improved quantitative photoconversions but a similar wavelength profile compared to **AzoF**.^[17,18]^ While some psUAAs from this current repertoire exhibit hydrogen bond acceptors in addition to the predominantly hydrophobic photoswitch scaffold, hydrogen bond donors or charged groups are hitherto missing. Moreover, all of them display high thermal stabilities necessitating a second irradiation step to achieve reversibility of photocontrol.

**Figure 1.**
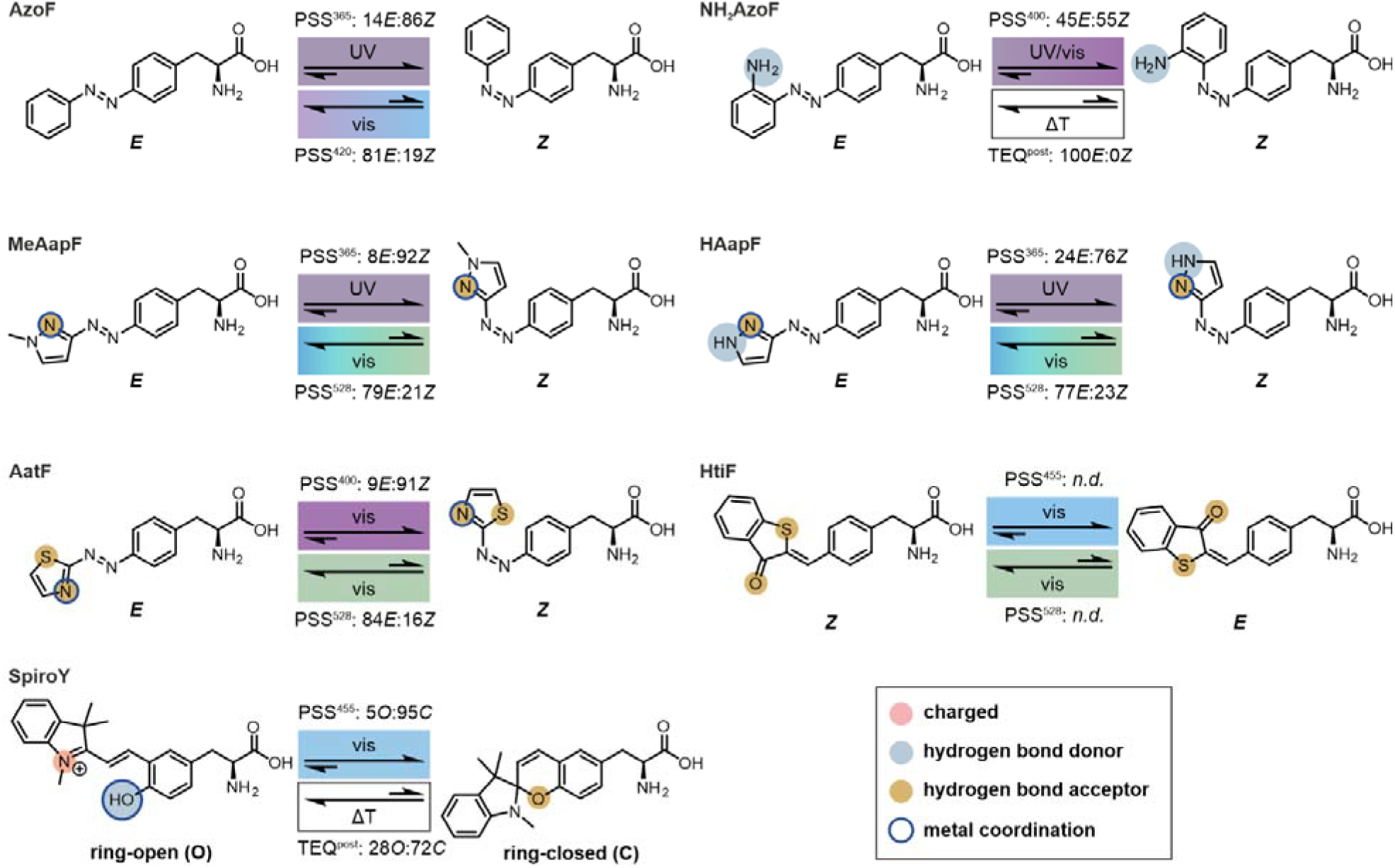
psUAA design and summary of photochemical properties. Photoisomerization upon irradiation with a specific wavelength *λ* in the UV (dirty violet) or visible (violet to green) range, or thermal relaxation (ΔT, white box) induce the formation of either a PSS^λ^ or a TEQ^post^, respectively. Each state is characterized by a certain isomer distribution as determined by HPLC or UV/Vis. Colored circles indicate functional groups for potential polar interactions.

In this study, we set out to improve the versatility of the psUAA repertoire for the recombinant production of photocontrolled enzymes using azobenzene, arylazopyrazole, arylazothiazole, hemithioindigo and spiropyran photoswitch scaffolds. We describe the synthesis of six psUAAs, which i) showed improved photoswitch efficiencies, ii) mostly contain hydrogen bond acceptors, donors and/or charged groups, or iii) exhibit variation in thermal stabilities as well as effective wavelengths of irradiation. Remarkably, we were able to identify suitable aaRS/tRNA pairs facilitating the incorporation of five of the six psUAAs into a model enzyme. Photochemical and kinetic characterization of the resulting enzyme variants demonstrated different behavior and photocontrol potential of each psUAA.

## Results and Discussion

### Design and synthesis

To obtain the intended psUAA diversity regarding photoswitch efficiencies, interaction potentials, thermal stabilities and effective wavelengths of irradiation in our design (**Figure 1**), we decided to exploit the current variety of photoswitch scaffolds. As a first step, we considered an **AzoF** derivative, **NH**_**2**_**AzoF**, that is different from previous designs (**Figure S1**) due to the presence of a primary amine that acts as a hydrogen bond donor and nucleophile. *Ortho*-substitution of azobenzenes with such an electron-donating group has been shown to considerably increase thermal relaxation of the *Z* isomer and red-shift the effective wavelength.^[19]^ Moreover, similar to the previously reported AapF,^[17,18]^ other arylazopyrazole photoswitches displayed equally high photoconversion yields.^[20,21]^ Hence, we designed **MeAapF** as a less sterically demanding version of AapF. While exchanging a functional group in the heteroarene ring may significantly alter the photochemical properties of arylazopyrazoles,^[20]^ the introduction of a secondary, free amine enhances the interaction and potential by serving as hydrogen bond donor or nucleophile. For this reason, we included **HAapF** in our design. In general, arylazopyrazoles as well as the structurally related arylazothiazoles are known to coordinate metal ions via their heteroaromatic moiety,^[22]^ which is appealing for metalloenzymes. Furthermore, the latter have recently emerged as attractive scaffolds with fast thermal relaxation of the *Z* isomer and irradiation wavelengths in the visible range.^[23]^ Hence, we tried to capture these properties in the design of **AatF**. Furthermore, we included photoswitches, which are based on other isomerization mechanisms than diazo compounds. To this end, we selected **HtiF** that has previously been incorporated into an ion channel via chemical acylation, but has so far not been photochemically characterized or incorporated via amber suppression.^[24]^ The olefinic bond of this hemithioindigo-based psUAA confers the ability to switch between *E* and *Z* isomers with visible light bidirectionally. **HtiF** is not only capable of interacting with other amino acids in the protein environment through π-stacking interactions but may also participate in hydrogen bonding.^[25]^ Finally, we included a psUAA that is able to undergo an electronical rearrangement resulting in the interconversion between an open (*O*) and a closed (*C*) state. To simultaneously increase the interaction potential, we chose **SpiroY**, which isomerizes between a flexible, charged merocyanine (*O*) and a rigid, neutral spiropyran (*C*). While the open isomer can serve as Brønsted base or complexation site for metal ions, the closed form is inert.^[26]^

Next, we synthesized the six psUAAs and **AzoF**, which we used for comparison in the subsequent characterization. To this end, four main synthetic routes have been established (**Figure 2**): Mills Coupling 1 of protected 4-amino-l-phenylalanine to nitroso derivatives (**AzoF, NH2AzoF**), Mills Coupling 2 of 3-aminopyrazole to protected 4-nitroso-l-phenylalanine (**HAapF**),^[27]^ Negishi Coupling of the iodo- or bromo-derivatives of the photoswitches to protected iodo-l-alanine (**MeAapF, HtiF, SpiroY**),^[28]^ and nucleophilic substitution of 2-hydrazineylthiazole to the oxaspiro[4,5]decadien derivative (**AatF**). A detailed description of each synthesis can be found in the Supporting Information (**Schemes S1–S6**). After flash chromatography purification, the psUAAs were isolated as colored solids with overall yields of 3–74%.

**Figure 2.**
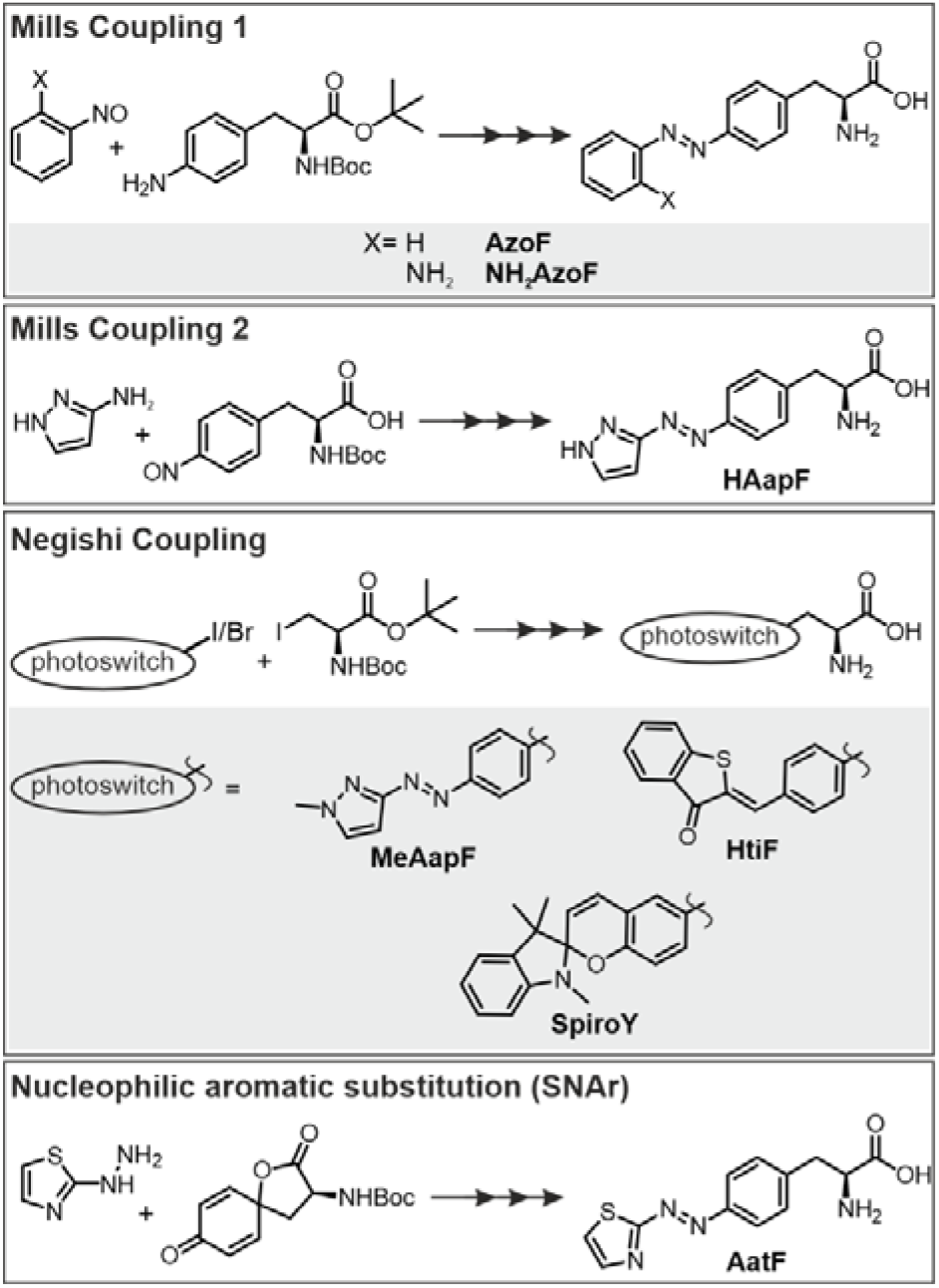
Schematic overview of the four main synthetic routes employed for the synthesis of psUAAs based on azobenzene, arylazopyrazole, arylazothiazole, hemithioindigo, and spiropyran photoswitches.

### Photochemical Characterization

After successful synthesis, we performed an extensive photochemical characterization of each psUAA in aqueous solution (50 mM Tris/HCl pH 7.5), to ensure compatibility with their biological application, as well as in DMSO. For this purpose, we acquired absorbance spectra of the TEQs, compared various PSS^λ^ spectra, determined the isomer distributions at the most relevant PSS^λ^, measured the half-lives of thermal relaxation, and evaluated the photostabilities over various switching cycles. We started our analysis by recording absorbance spectra of the TEQs (**Figure 3, Figure S2**) and identifying typical maxima for each photoswitch (**Table S1**). The TEQ spectra of **AzoF, NH**_**2**_**AzoF, MeAapF**, and **HAapF** were characteristic of the respective *E* isomers^[6,12,20,21]^ revealing pronounced π→π* transitions in the range of 290 nm to 335 nm and smaller n→π* transitions at 400–490 nm. Remarkably, the n→π* transition of **NH**_**2**_**AzoF** was broadened, which opens the possibility of red-shifted effective wavelengths for *E→Z* isomerization. While spectra in DMSO were similar to those in buffer for **AzoF, MeAapF** and **HAapF**, we observed a slight shift of the n**→**π* transition towards higher wavelengths for **NH**_**2**_**AzoF**. As reported for arylazothiazoles,^[23]^ **AatF** exhibited one major band in both buffer and DMSO for the π→π* transition at ∼370 nm indicative of the *E* isomer. Similar to **NH**_**2**_**AzoF**, the pronounced red-shift of this band promises effective isomerization wavelengths in the visible range. Notably, solubility of **HtiF** in buffer was quite low, which is in compliance with recent observations of high aggregation tendencies of hemithioindigos in the absence of organic (co)solvents.^[29]^ The absorbance spectra of the thermally stable *Z* isomer of **HtiF** in both buffer and DMSO demonstrated two absorbance maxima at ∼330 nm and ∼440 nm, which is in agreement with reported hemithioindigo spectra.^[30]^ Interestingly, we observed a special scenario for **SpiroY**. Previous studies have shown that spiropyrans are stabilized in their positively charged *O* isomer by high salt concentrations in organic solvents.^[31]^ Accordingly, the TEQ absorbance spectrum of **SpiroY**, recorded directly after dilution into buffer from the salt-enriched DMSO stock solution, exhibited absorbance characteristics of the conjugated *O* isomer. However, the absorbance signal decreased towards a new TEQ^post1^ with *t*_*½*_^*post1*^ = 1.9 min (**Figure S3A**) indicating the deprotonation of *O* due to the lower salt concentration and thus isomerization to the colorless *C* isomer. We observed similar, however, significantly slower (*t*_*½*_^*post1*^ = 17.3 h) spectral changes upon dilution in DMSO (**Figure S3B**).

**Figure 3.**
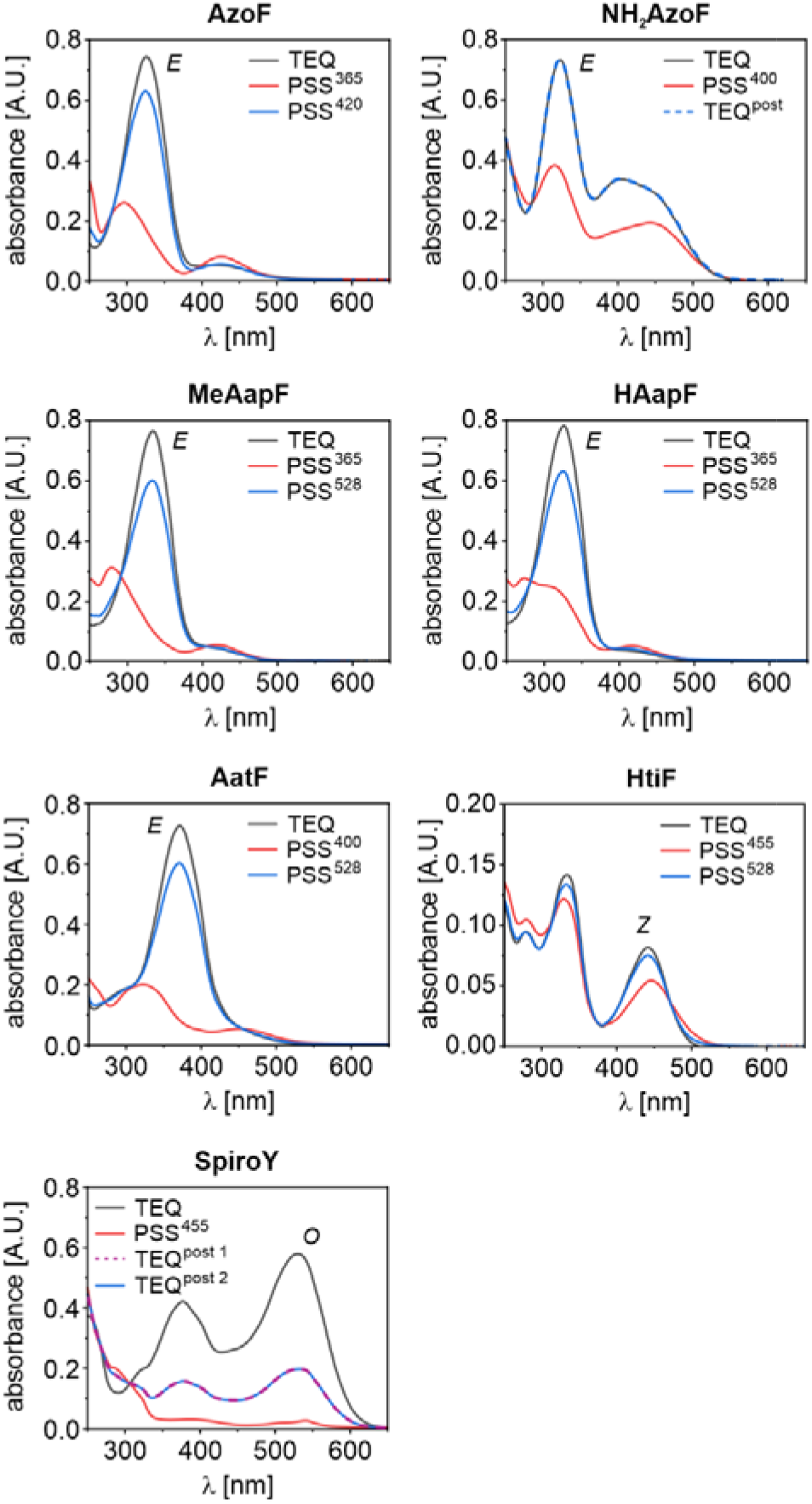
Absorbance spectra of the psUAAs in buffer. Real-time tracking of the absorbance signal assured the complete establishment of the respective PSS^λ^ or TEQ^post^ prior to spectrum acquisition (**Table S2** and **Figure S6**). The most abundant isomer in the TEQ is indicated next to the grey spectrum.

To initiate the photoinduced isomerization of each psUAA, we tested different wavelengths in the range of 365 nm and 567 nm (**Figures S4–S5**) but excluded wavelengths <365 nm owing to their ineptitude for biological applications. We thereby monitored the absorbance at the defined maximum during irradiation until each PSS^λ^ was fully established (**Tables S2–S3, Figures S6–S7**) and then recorded spectra. In accordance with the spectral properties evident from the TEQ spectra, the most effective irradiation wavelengths towards a PSS^λ^ enriched in the thermally less stable isomer were in the UV range for **AzoF, MeAapF** and **HAapF**, and in the near-visible to green range for **NH**_**2**_**AzoF, AatF, HtiF** and **SpiroY**. Instead, photoinduced reestablishment of a PSS^λ^ similar to TEQ was accomplished exclusively with visible light for all psUAAs except **NH**_**2**_**AzoF** and **SpiroY**. However, both psUAAs exhibited a thermally unstable *Z* quantitative switching in the *E*→*Z* isomerization and mediocre to good photoconversion in the *Z*→*E* isomerization, outperforming **AzoF** and **HAapF**. HPLC analysis of **NH**_**2**_**AzoF** and **HtiF** was unsuccessful owing to the extremely short *t*_*½*_(ΔT) and solubility issues,^[29]^ respectively. **SpiroY** demonstrated a nearly quantitative photoinduced switch to 5*O*:95*C* (PSS^455^) and a and *C* isomer, which allowed them to relax spontaneously to TEQ^post^ and TEQ^post2^, which are equivalent to TEQ and TEQ^post1^, respectively. For both isomerization processes, the most effective wavelengths in buffer (**Figure 3**) and DMSO (**Figure S2**) were either the same or similar for each of the six psUAAs.

We then evaluated the thermal relaxation of the less stable isomers for each psUAA by determining the thermal half-lives *t*_*½*_(ΔT) in buffer at 25°C (**Table 1**). While **AzoF, MeAapF, HAapF** and **AatF** offer bistability with *t*_*½*_(ΔT) values between hours to several days, **NH**_**2**_**AzoF, HtiF** and **SpiroY** showed rapid full reversibility within <20 minutes (**Figure S8–S9**).

**Table 1.** Half-lives *t*_*½*_(ΔT) of thermal relaxation for each psUAA in buffer at 25°C.

| psUAA | $t_{1/2}(\Delta T)^{[a]}$ | unit |
| --- | --- | --- |
| **AzoF** | $16.4 \pm 0.40^{[b]}$ | d |
| **NH<sub>2</sub>AzoF** | $9.91 \pm 0.02^{[c]}$ | s |
| **MeAapF** | $45.6 \pm 1.70^{[b]}$ | d |
| **HAapF** | $10.3 \pm 0.20^{[b]}$ | d |
| **AatF** | $1.80 \pm 0.004^{[c]}$ | h |
| **HtiF** | $2.04 \pm 0.003^{[c]}$ | min |
| **SpiroY** | $1.90 \pm 0.0001^{[c]}$ | min |

Next, we assessed the quantity of photoconversion by separating the isomers via HPLC following the absorbance at specific isosbestic points (**Table S4, Figures S10–S11**). As a result, we obtained isomer distributions for TEQ, PSS^λ^ and TEQ^post^ in buffer (**Figure 1, Table S5**) and in DMSO (**Table S6**). In buffer, all psUAAs except **AzoF** (97*E*:3*Z*) and **SpiroY** (93*O*:7*C*) revealed 100% of the thermodynamically more stable isomer in TEQ. The latter can be explained by the above-described establishment of TEQ^post1^ (28*O*:72*C*) after dilution. We first evaluated the photoinduced switch of the bistable psUAAs, obtaining isomer distributions of 14*E*:86*Z* for **AzoF** (PSS^365^), 8*E*:92*Z* for **MeAapF** (PSS^365^), 24*E*:76*Z* for **HAapF** (PSS^365^), and 9*E*:91*Z* for **AatF** (PSS^400^). Moreover, the photoinduced backswitch established 81*E*:19*Z* for **AzoF** (PSS^420^), 79*E*:21*Z* for **MeAapF** (PSS^528^), 77*E*:23*Z* for **HAapF** (PSS^528^), and 84*E*:16*Z* for **AatF** (PSS^528^). Conclusively, **MeAapF**, and **AatF** achieved nearly thermal relaxation to 28*O*:72*C* (TEQ^post2^) indicating a complete reestablishment of TEQ^post1^. In DMSO, isomer distributions of all psUAAs coincided with those in buffer with slight deviations for **SpiroY**, which has been reported to be sensitive to solvent changes.^[31,32]^ While the measurements for **NH**_**2**_**AzoF** were again inconclusive, **HtiF** showed isomer distributions of 77*E*:23*Z* (PSS^420^) and 2*E*:98*Z* (PSS^505^).

Although HPLC provides an exact method for the determination of isomer distributions, it is not applicable for the evaluation of psUAAs incorporated into a protein. Hence, we further examined the isomer ratios using UV/Vis spectra, an approach that is commonly used for diazo photoswitches.^[20]^ Overall, the estimated values were in good agreement with the HPLC results (**Tables S5–S6**). In this way, we were also able to yield an isomer distribution upon irradiation in buffer for **NH**_**2**_**AzoF** of 45*E*:55*Z* (PSS^400^) and fully recover 100*E*:0*Z* in TEQ^post^. It should be noted that the fast thermal relaxation of **NH**_**2**_**AzoF** counters its photoinduced isomerization. As a result, the isomer distribution depends on the intensity of irradiation as is observed for photoreceptors used in optogenetics,^[33]^ opening up the possibility to further tune the PSS^λ^ composition. While there are currently no UV/Vis-based methods to estimate the isomer ratio for **HtiF**, we obtained similar values for **SpiroY** compared to the HPLC evaluation by exploiting the lack of an absorbance signal in *C* using the rule of three.

Finally, we assessed the photostability of each psUAA over ten switching cycles (**Figures S12–S13**). Except **HtiF**, which indicated slight photobleaching in buffer, all psUAAs maintained high photostability without fatigue in both buffer and DMSO.

In conclusion, we developed a repertoire of psUAAs with high diversity (**Figure 1**). Moreover, the psUAAs demonstrated the anticipated photochemical properties typical for each photoswitch,^[20,23,30,34]^ which constitutes beneficial knowledge for the rational chemical design of psUAAs.

### Screening of aminoacyl-tRNA synthetases (aaRSs) for selective incorporation of each psUAA

We further explored the incorporation of the psUAAs during recombinant protein production via amber suppression. To this end, we have screened a selection of aaRSs for their ability to efficiently aminoacylate the orthogonal tRNA with each synthesized psUAA. All chosen aaRSs were based on the tyrosyl-tRNA synthetase from *Methanocaldococcus jannaschii* (*Mj*Tyr-RS). Although *Mj*Tyr-RSs are only orthogonal in prokaryotic cells, they are more adapted to binding aromatic amino acids and have previously achieved exceptionally high incorporation efficiencies.^[35,36]^ Accordingly, for the incorporation of our aromatic psUAAs, we used established *Mj*Tyr-RS variants that contain amino acid exchanges at defined positions (Y32, L65, E107, F108, Q109, Q155, D158, I159, L162, A167, V188, R257, and D286).^[12,18,35,37]^ Moreover, we performed a computational design based on the original AzoF-RS^[12]^ (aaRS-a), from which we selected further variants. The resulting library contained 22 *Mj*Tyr-RSs (a-v; **Table S7–S8**).

Screening of the synthetases was accomplished using two plasmids: i) pETBAD encoding the reporter sfGFP with an amber stop codon (TAG) at position Y151 (pETBAD_sfGFP_Y151TAG), and ii) pGLNS harboring an aaRS variant and the respective tRNA. We initially optimized the screening procedure using **AzoF** (**Figure S14**), which resulted in low background signals when pGLNS without aaRS was employed as negative control (−; **Figure 4A**). sfGFP expression in the presence of **AzoF** was examined by measuring fluorescence intensities for each culture containing aaRSs a–**v** in biological triplicates. Interestingly, aaRS-a yielded relatively low intensities, probably due to the use of only one gene copy compared to the commonly used two copies in the pEVOL plasmid system.^[38]^ Since pEVOL_aaRS-a usually facilitates reasonable incorporation efficiencies in large scale,^[6,7]^ we set the respective fluorescence value as lower threshold (solid line) for the identification of suitable aaRSs for the other psUAAs. Moreover, we defined a second threshold (dashed line) at 30% of the wild type sfGFP signal of the positive control (+) to determine aaRSs with high incorporation efficiencies. Importantly, we excluded misincorporation of natural amino acids for each aaRS, except aaRS-s, by performing the screening experiment in the absence of psUAAs (**Figure S15**).

**Figure 4.**
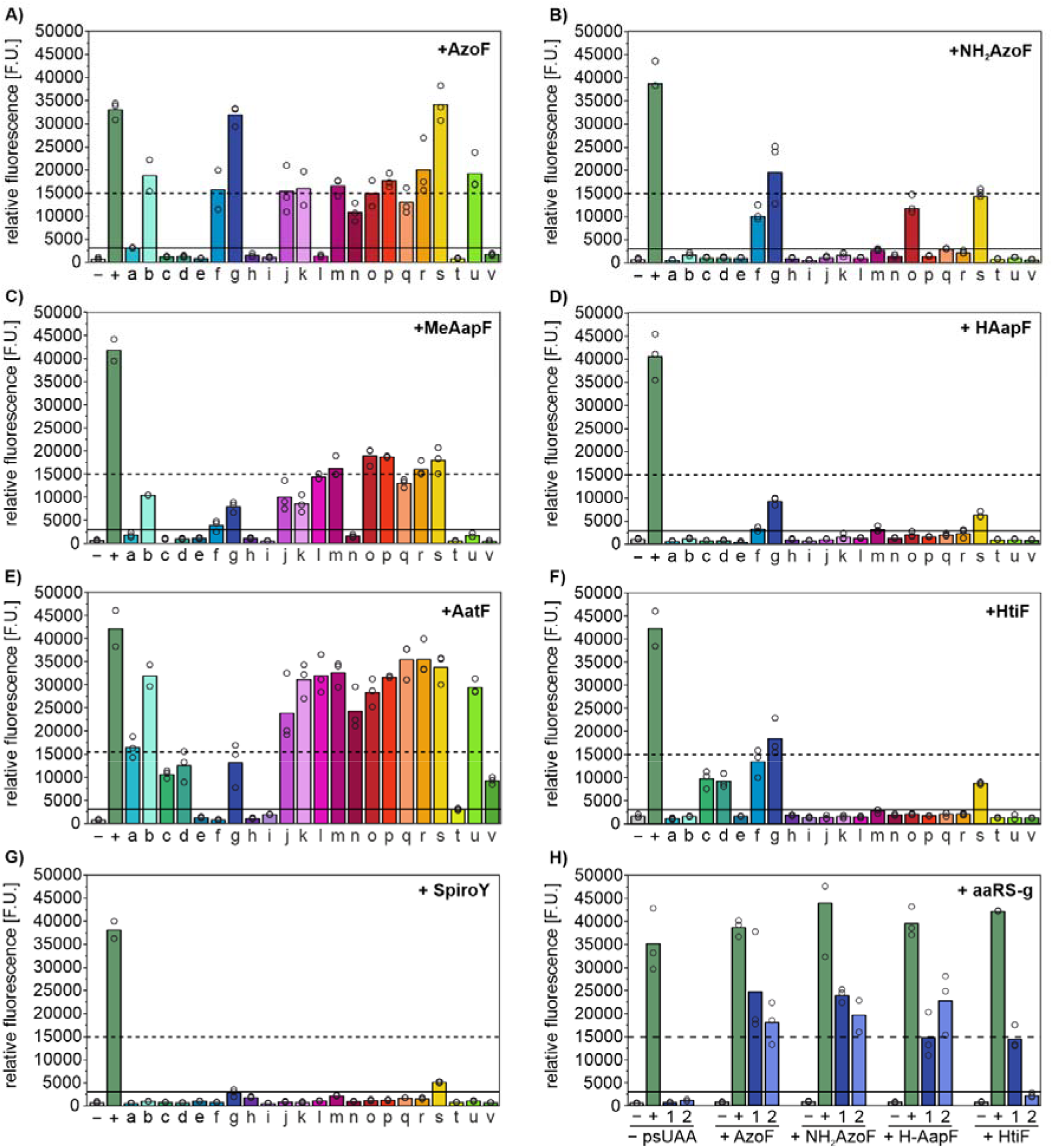
Screening of *Mj*Tyr-RS variants (a-v) using pGLNS for the orthogonal incorporation of psUAAs in sfGFP_Y151TAG using pETBAD_sfGFP_Y151TAG (A–G) and influence of aaRS copy number (H). Empty circles: fluorescence intensity of three biological replicates; columns: mean fluorescence intensity; “−”: pGLNS without aaRS; “+”: wild type sfGFP; 1: pGLNS_aaRS-g; 2: pEVOL_aaRS-g.

To our astonishment, the codon-optimized variant aaRS-b harboring the additional R257G mutation significantly outperformed aaRS-a regarding **AzoF** incorporation, and expression was even further enhanced with aaRS-g, another previous AzoF-RS design (**Figure 4A**).^[35]^ Remarkably, we observed fluorescence intensities above the second threshold for **NH**_**2**_**AzoF, MeAapF, AatF** and **HtiF** with the highest signals obtained using aaRS-g, aaRS-p, aaRS-q and aaRS-g, respectively (**Figure 4B–G**). While aaRS-g yielded fluorescence intensities in the presence of **HAapF** between both thresholds, no aaRS could be identified for the incorporation of **SpiroY. SpiroY**-*C* is structurally rigid and bulky, which may significantly hinder binding to an aaRS. Moreover, the charged state of **SpiroY**-*O*, which is potentially also present in large amounts within the expression host (cf. TEQ^post1^), may interfere with the hydrophobic binding pocket of the *Mj*Tyr-RSs. Hence, we anticipate that successful incorporation of **SpiroY** will require more extensive engineering efforts of aaRSs.

To further boost the incorporation efficiency of **HAapF**, we compared sfGFP expression yields obtained using either one copy of aaRS-g in pGLNS or two copies in pEVOL (**Figure 4H**).^[38]^ Again, misincorporation of natural amino acids in the absence of psUAA could be excluded for both expression systems. Interestingly, the new biological replicates demonstrated fluorescence intensities approximating the second threshold in the presence of pGLNS_aaRS-g, and even surpassed it with pEVOL_aaRS-g. We also tested the pEVOL system for **AzoF, NH**_**2**_**AzoF** and **HtiF**, which showed the highest incorporation yields with aaRS-g. However, the fluorescence and conclusively, the expression level remained similar or decreased with pEVOL_aaRS-g compared to pGLNS_aaRS-g. The overall observed differences and fluctuations in expression levels might be associated with off-target interactions of the psUAAs and aaRSs with host cell components or with perturbations in energy metabolism.^[39]^ Previous studies have particularly shown that overexpression of both an aaRS and a gene of interest can deplete translational factors and metabolites, especially if the aaRS/tRNA pair is highly efficient.^[40]^

Finally, we were interested in the binding mode of **AzoF, NH**_**2**_**AzoF, MeAapF, HAapF, AatF**, and **HtiF** with their respective aaRSs. To that end, we constructed a cluster model of the following combination of *Mj*Tyr-RS variants and psUAAs: **AzoF, NH**_**2**_**AzoF, HAapF, HtiF** in aaRS-g, **MeAapF** in aaRS-p, and **AatF** in aaRS-q (**Table S9**). The studied *Mj*Tyr-RS variants contain mutations mostly in the psUAA binding region. In all three aaRS-g, -p, and -q, mutation Y32G is introduced, which replaces the sterically bulky Y32 for a substantially smaller and more flexible glycine (**Figure 5**). This mutation is clearly needed to generate additional space to allow for binding of all psUAAs. In aaRS-g, the space left by Y32G substitution can be potentially occupied by the new arginine residue included at position 162 (L162R, **Figure S16**). However, for **AzoF, NH**_**2**_**AzoF, HtiF**, and **HAapF** binding, R162 needs to be displaced from the pocket (**Figure 5A**). It should be noted that such conformation of R162 has also been reported in previous X-ray structures (PDB: 6WRT) and is also predicted by AlphaFold2 (**Figure S16**). Both aaRS-p and -q present a histidine residue at position 162 instead (L162H), which establishes a hydrogen bond with the new glutamate residue included at position 65 (L65E). This hydrogen bond is also maintained when **AatF** and **MeAapF** are bound in the pocket (**Figure 5B–C**). In aaRS-g position 65 contains instead a valine (L65V) to allow enough space for **AzoF, NH**_**2**_**AzoF, HtiF**, and **HAapF** binding (**Figure S17**). Another important mutation for providing additional space is D158 to either glycine (in aaRS-g and -p) or alanine (in aaRS-q). Only in aaRS-g a tyrosine residue is introduced at position 159 (i.e., I159Y), which establishes a hydrogen bond with H177 and hydrophobic interactions with the aromatic rings of the different UAAs (**Figure 5A**). In aaRS-p and -q, two additional mutations are included: the bulky F108 is replaced by a smaller alanine, and Q109 is replaced by either a methionine (Q109M) or glutamate (Q109E), respectively. Our calculations indicated that E109 establishes a hydrogen bond with H70, altering the sidechain conformation of H70 and thus, providing additional space for efficient **AatF** incorporation.

**Figure 5.**
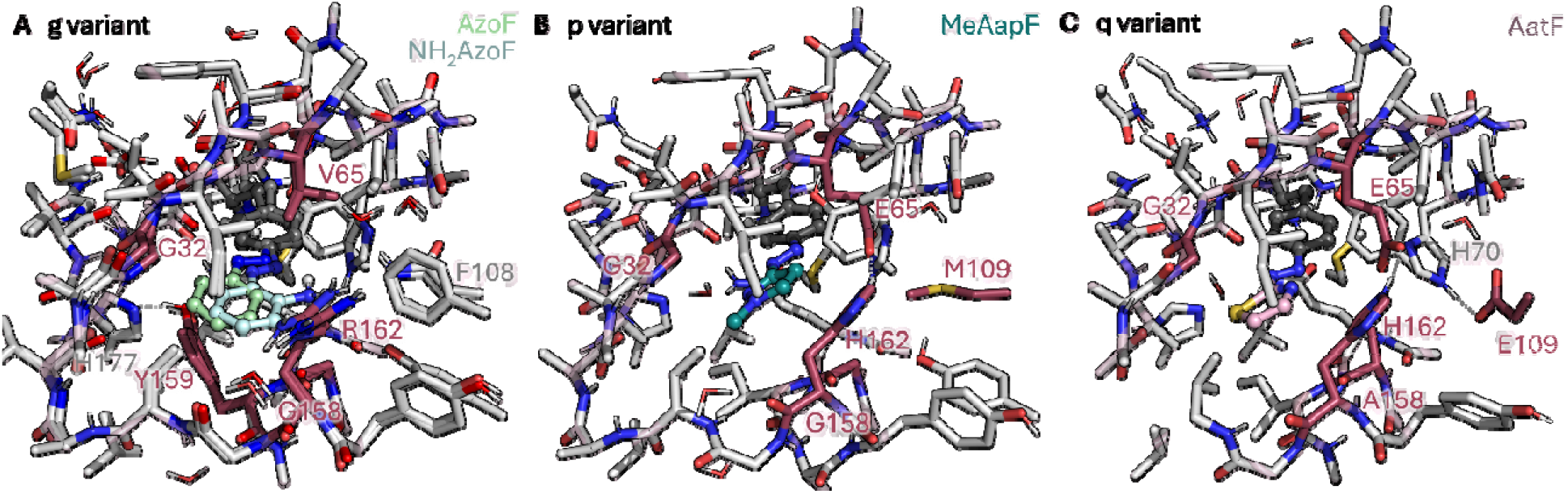
Optimized cluster models of *Mj*Tyr-RS variants aaRS-g, -p, and -q. (**A**) Overlay of the optimized structures of aaRS-g containing exemplarily either **AzoF** (shown in dark grey and green spheres) or **NH**_**2**_**AzoF** (in grey and cyan). (**B**) Optimized structure of the aaRS-p containing **MeAapF** bound in the pocket. (**C**) Optimized structure of aaRS-q containing **AatF** bound in the pocket. All mutations are colored in raspberry.

Ultimately, our findings make it possible to effectively use **NH**_**2**_**AzoF, MeAapF, HAapF, AatF**, and **HtiF** for the recombinant production of potentially light-sensitive enzymes in prokaryotes. For studies in eukaryotic cells, that are beyond the scope of this work, we recommend to test orthogonal aaRSs based on pyrrolysyl-tRNA synthetases, which have been evolved for incorporation of AzoF derivatives,^[8,14,15]^ or tyrosyl-tRNA synthetases from *Geobacillus stearothermophilus*.^[41]^

### Incorporation into an enzyme and characterization

The identification of aaRSs for the co-translational incorporation allowed us to further characterize **NH**_**2**_**AzoF, MeAapF, HAapF, AatF** and **HtiF** in comparison to **AzoF** within an enzymatic environment. For this purpose, we selected HisF from *Thermotoga maritima*, which catalyzes the reaction of *N’*-[(5’-phosphoribulosyl)formimino]-5-amino-imidazole-4-carboxamide ribonucleotide (PrFAR) and NH_3_ to imidazole glycerol phosphate and 5-aminoimidazol-4-carboxamidribotide (**Figure S18**), because of several advantageous aspects. Its heat-stability allows for a highly efficient purification by performing a heat-step and more importantly, the catalyzed reaction can be monitored directly by following the decline in absorbance of PrFAR. Moreover, hitherto unpublished data of HisF variants containing **AzoF** at various positions was available from our previous project.^[7]^ This data provided a comparison of expression yields and activities before and after irradiation for each HisF-AzoF variant and demonstrated that wild type HisF (HisF-wt) is largely unaffected by irradiation (**Figure S19**). For the incorporation of our novel psUAAs, we aimed for a position, in which **AzoF** incorporation facilitated >50% of wildtype activity, and potentially a slight activation of catalysis upon irradiation. Only two positions, K13 and L35 met these criteria, of which we chose K13 owing to apparently higher expression yields.

Using the corresponding aaRS identified in the screening, we produced HisF variants containing either **AzoF, NH**_**2**_**AzoF, MeAapF, HAapF, AatF**, or **HtiF** at position K13, delineated as “K13psUAA”. The enzymes were derived in moderate to high yields and in moderate to excellent purities (**Table 2**; **Figure S20**). It should be noted that differences in yield may not only result from distinct efficiencies of the respective aaRSs, but also from the compatibility between the psUAA and the site of incorporation.^[42]^ Tryptic digest coupled to mass spectrometry confirmed the identity of each K13psUAA variant (**Figure S21**).

**Table 2.**
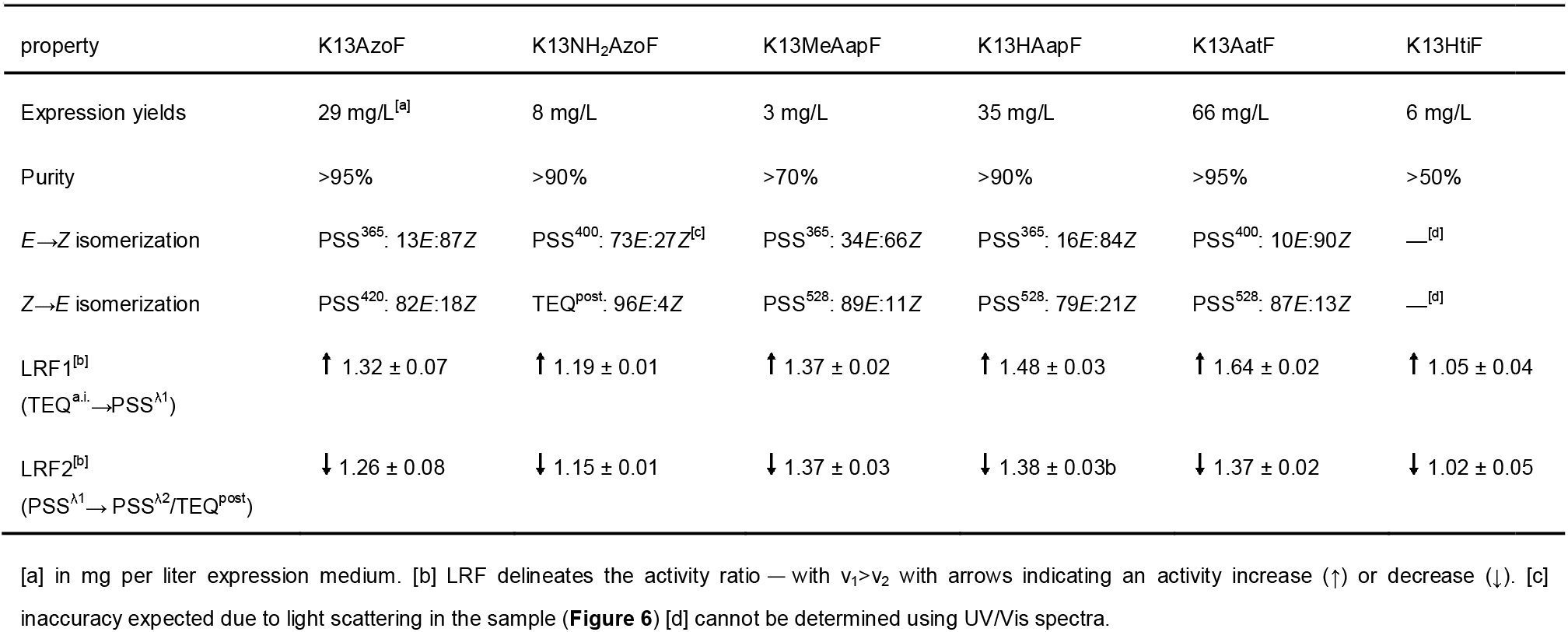
Properties of HisF variants containing psUAAs at position K13.

Since the chemical environment within a protein strongly diverges from that in buffer or solvent, we performed a photochemical characterization of the psUAAs within each HisF variant. Similar to the isolated psUAAs, we acquired absorbance spectra of the thermally equilibrated, as-isolated enzymes (TEQ^a.i.^), compared PSS^λ^ or TEQ^post^ absorbance spectra, estimated the respective isomer distributions at PSS^λ^, and evaluated the photostabilities over various switching cycles.

Absorbance spectra of each K13psUAA variant in TEQ^a.i.^ comprised protein-related maxima at ∼280 nm in addition to the characteristic maxima of each photoswitch (**Figure 6**). The absorbance maxima of the π→π* and n→π* transitions (**Table S10**) thereby coincided well with those of the isolated psUAAs. General differences in signal strength of the photoswitches, particularly compared to the protein signal, might be attributed to quenching effects by the adjacent protein environment. Photoinduced isomerization in both directions was then monitored in real-time using the most effective wavelengths of the isolated psUAAs in buffer and DMSO (**Figure S22**; **Table S11**). K13AzoF, K13MeAapF, K13HAapF and K13AatF showed well resolved isomerization profiles. However, K13NH_2_AzoF and K13HtiF, which already exhibited weak signals in TEQ^a.i.^, demonstrated only mediocre or even minor changes upon irradiation. Considering the higher absorbance values close to the protein signal (**Figure 6**), we assume that both enzymes contained soluble aggregates that could not by removed by centrifugation and that caused light scattering. We further obtained the thermal half-lives of K13NH_2_AzoF (*t*_*½*_ =14 s) and K13AatF (*t*_*½*_ =80 min) (**Figure S22**) as well as the spectra of each PSS^λ^ or TEQ^post^, which aligned well with those of the isolated psUAAs, particularly regarding the spectral behavior of the π→π* and n→π* transitions (**Figure 6**).

**Figure 6.**
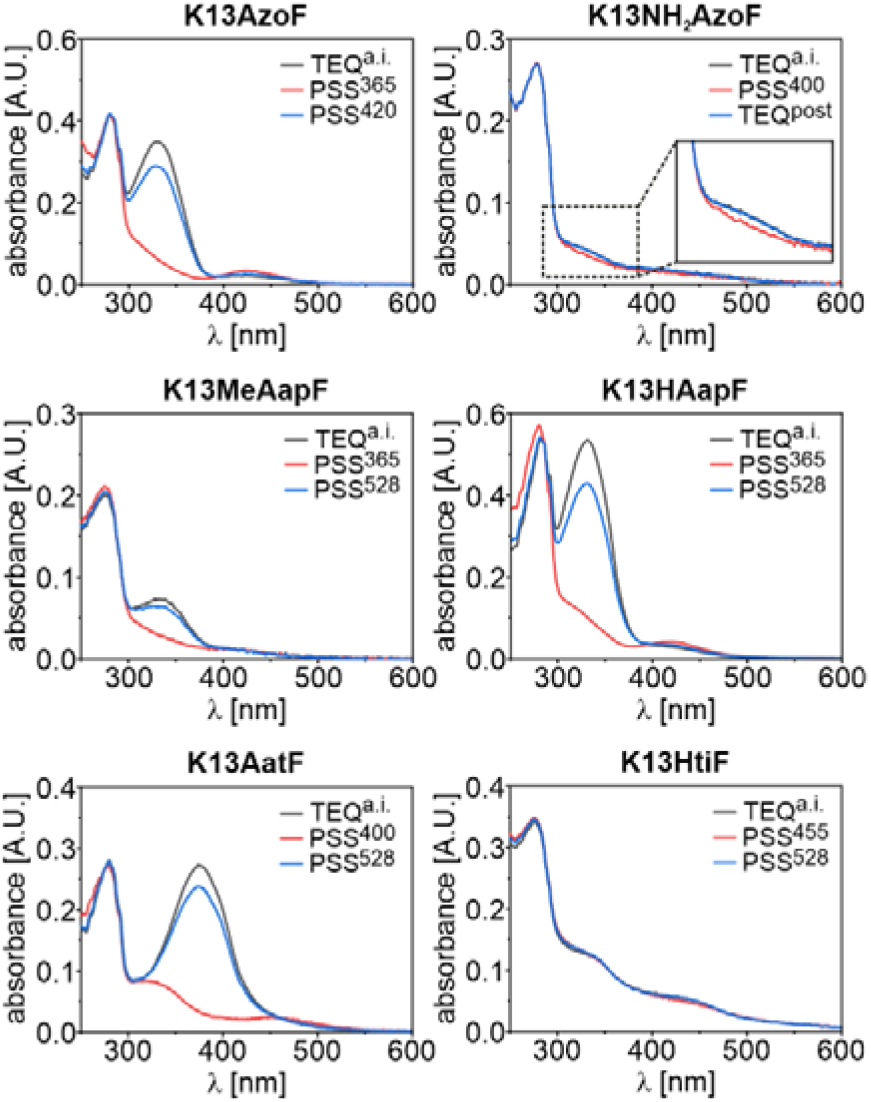
Absorbance spectra of HisF-psUAA variants in buffer. Real-time tracking of the absorbance signal assured the complete establishment of the respective PSS^λ^ or TEQ^post^ prior to spectrum acquisition (e.g. **Figure S22**).

In the next step, we estimated the isomer distributions at the PSS^λ^ and TEQ^post^ for the HisF variants containing diazo-based psUAAs from UV/Vis spectra. For this, TEQ^a.i.^ was assumed to contain 100% *E*.^[20]^ For the *E*→*Z* isomerization, we observed nearly quantitative (K13AatF), good (K13AzoF, K13HAapF) and mediocre (K13NH_2_AzoF, K13MeAapF) photoconversion yields (**Table 2**). Likewise, *Z*→*E* isomerization obtained either quantitative (K13NH_2_AzoF), good (K13AzoF, K13MeAapF, K13AatF) or mediocre (K13HAapF) *E* isomer yields. Notably, the highest isomer distributions for K13MeAapF and K13HAapF, for which the most effective *Z*→*E* isomerization wavelengths in buffer and DMSO differed, were obtained with 528 nm (**Figure S23**). Likewise, we tested all isomerization wavelengths that were effective in buffer and DMSO for K13HtiF. However, since potential differences in photoconversion remained elusive owing to pronounced light scattering, we chose to use 455 nm and 528 nm, which were most effective in buffer, for further activity studies.

Overall, the determined isomer distributions aligned well with those obtained from the isolated psUAAs for **AzoF** and **AatF**, whereas **MeAapF** and **HAapF** demonstrated partly less or more quantitative photoconversions. **NH**_**2**_**AzoF** appeared to switch only inefficiently in the enzyme environment, however, the isomer distributions might be significantly underestimated due to light scattering. The photochemical characterization of **HtiF** was even more difficult and less informative. To reassess the photochemical behavior of **HtiF** within a different location, we produced the variant L35HtiF. While position K13 points into the solvent, L35 is part of a hydrophobic pocket, which might protect the largely hydrophobic **HtiF** from the solvent (**Figure S24**). However, L35HtiF also exhibited light scattering, an even less pronounced absorbance signal compared to K13HtiF, and no signal change upon irradiation (**Figure S25**). Further studies will show which positions, and more specifically which chemical environment, will be more suited for the incorporation of **HtiF**.

Finally, cycle performance measurements revealed that each HisF-psUAA variant, except K13HtiF, was able to withstand ten cycles of repeated photoswitching without any obvious signs of photobleaching or light scattering (**Figure S26**).

With this, we confirmed that the diversity of properties regarding the photoswitch efficiencies, the thermal stabilities and the effective wavelengths of irradiation was largely maintained after the psUAAs were incorporated into an enzyme.

### Photocontrol of HisF activity

In a first approach to assess the capability for enzymatic photocontrol, we measured HisF activities in the TEQ^a.i.^, PSS^λ^, and/or TEQ^post^ for each HisF-psUAA variant. The light-regulation-factor (LRF), which constitutes the ratio of activities *v* before and after irradiation (with *v*_*1*_>*v*_*2*_), thereby defines the photocontrol efficiency. After reconfirming that HisF-wt is not affected by irradiation (LRF ≤1.05; **Figure S27**), we initially determined the LRF of fK13AzoF at PrFAR concentrations either in saturation (LRF <1.2) or close to the *K*_*m*_ value (2.5 µM)^[7]^ of HisF-wt (LRF >1.2) and chose the latter for all subsequent measurements (**Figure S28**). Based on this result, we expected modest LRF values in the range of 1.2–2.0. Hence, we used an interaction model in a global fit analysis of two data sets, e.g., TEQ^a.i.^ and PSS^365^, to obtain LRF values ± standard errors (S.E.) (**Table S12**). This allowed us to evaluate whether differences in LRF values between the HisF-psUAA variants are reliable. Upon *E*→*Z (Z*→*E* for HtiF) isomerization we observed either a relatively high (K13AatF), good (K13AzoF, K13MeAapF, K13HAapF), minor (K13NH_2_AzoF) or insignificant (K13HtiF) increase in activity (**Table 2**). Remarkably, activities returned to initial values either with nearly full (K13MeAapF), good (K13AzoF, K13NH_2_AzoF, K13HAapF) or mediocre (K13AatF) reversibility. Notably, despite the light scattering effects, K13NH_2_AzoF and K13HtiF, retained well measurable enzymatic activities.

## Conclusion

Protein engineering with psUAAs has become a highly promising method to design photocontrolled enzymes, particularly for the two pioneering applications biotherapy and biocatalysis. However, the current repertoire of psUAAs has been limited in terms of quantitative switching, whether through irradiation or thermal relaxation, interaction potentials with the enzyme environment, and effective wavelengths of irradiation. In this work, we have enhanced the versatility of the repertoire by designing and synthesizing six psUAAs with diverse photochemical behaviors, five of which can be incorporated to facilitate advanced photocontrol of enzymatic reactions.

We have used **AzoF**, the best-known psUAA to date, as the point of reference. This bistable compound achieves good photoconversions in both directions with UV and visible light, but only offers hydrophobic interactions with the enzyme, with which it is possible to induce catalytically relevant conformational changes. In comparison, the visible light and thermally induced switch between a charged and neutral state in **SpiroY** is extremely interesting for the photocontrol of enzyme chemistry, not least due to its ability to coordinate metal ions in one of its isomeric forms, which is highly significant for metalloenzymes. However, its unique behavior also results in non-quantitative switching that greatly depends on the presence of charge stabilizers in close proximity. Moreover, it is the only psUAA for which further engineering studies, likely involving directed evolution and/or computational design, are required to identify an aaRS for the co-translational incorporation. Similar to **SpiroY, HtiF** appears to be a blackbox regarding photoconversion yields, although full recovery of the TEQ is possible by thermal relaxation. Nevertheless, **HtiF** offers a hydrogen bond acceptor and can be switched purely with visible light. Furthermore, while the two bistable arylazopyrazole-based psUAAs, **MeAapF** and **HAapF**, showed only mediocre to good photoconversion yields upon irradiation with UV and visible light, they can act as hydrogen bond acceptors and coordinate metals. Notably, **HAapF** and **NH**_**2**_**AzoF** can potentially participate as nucleophiles in catalysis, e.g., in Friedel-Crafts alkylation reactions and are, thus, quite valuable for biocatalysis. Additionally, **NH**_**2**_**AzoF** is a hydrogen bond donor and can be switched purely with violet light. Although we merely observed mediocre photoconversions for this psUAA, its switching efficiency may allow for further fine-tuning by modulating the intensity of irradiation to effectively compete with its fast thermal relaxation. This also ensures full reversibility to TEQ, allowing for exact spatial control with automatic deactivation of the enzyme once it exits the target site. All of these attributes make **NH**_**2**_**AzoF** one of the most interesting psUAAs of this study. Finally, another highly appealing psUAA for enzyme engineering is **AatF**. It demonstrates excellent photochemical performance by revealing good to nearly quantitative photoconversion in both directions with visible light and short-time bistability with the option to recover TEQ by thermal relaxation within hours. Most remarkably, it offers a versatile interaction potential with enzymes provided by its capability to act as hydrogen bond acceptor and to coordinate metals.

In conclusion, we expect the here described psUAAs to significantly increase the success rate of photocontrol engineering in enzymes. The extended repertoire promises to ease the identification of positions, in which enzyme conformation and hence catalysis can be controlled by the isomer switch. Moreover, it paves the way for photocontrolled catalysis via photoswitchable catalytic or metal-coordinating residues in the redesign of existing enzymes or in the de novo design of novel enzymes.

## Supporting information

aaRS-design files

## Supporting Information

The authors have cited additional references within the Supporting Information.^[43]^

## Acknowledgements

The authors thank Reinhard Sterner for his valuable support throughout the project as well as Alexander Deiters for mentoring C.H.. Moreover, we highly appreciate the excellent technical assistance of Sabine Laberer (protein biochemistry), Dennis Grimm and Andreas Pielmeier (synthesis of psUAAs). Furthermore, we like to thank Patricia Luckner and Eduard Hochmuth for their help with mass spectrometry. This work was funded by the Deutsche Forschungsgemeinschaft (STE 891/12-2).

## Entry for the Table of Contents

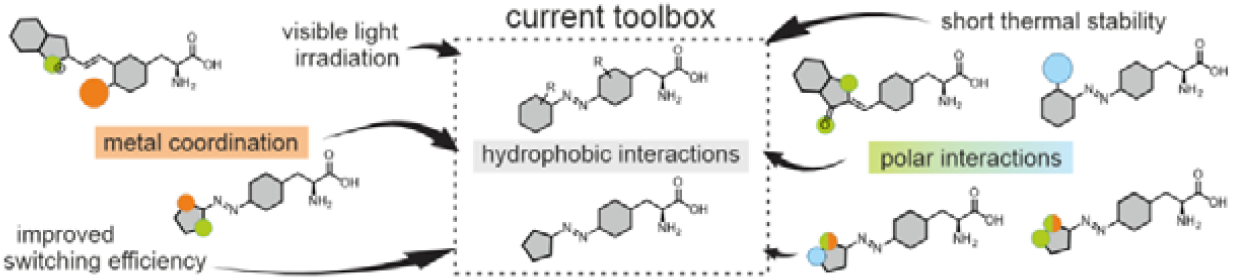

Enzyme engineering with photoswitchable unnatural amino acids (psUAAs) has high potential for applications such as biocatalysis. Here we extend the current repertoire of psUAAs to include more versatile properties paving the way to more advanced photocontrol engineering. We report on the synthesis and photochemical behavior, suitable aminoacyl-tRNA synthetases for co-translational incorporation, and initial enzymatic studies of these psUAAs.

