## Supporting Information for "Expanding the Repertoire of Photoswitchable Unnatural Amino Acids for Enzyme Engineering"

#### Introducing High Versatility for Enzyme Engineering in the Repertoire of Photoswitchable Unnatural Amino Acids

### 1. Supporting material

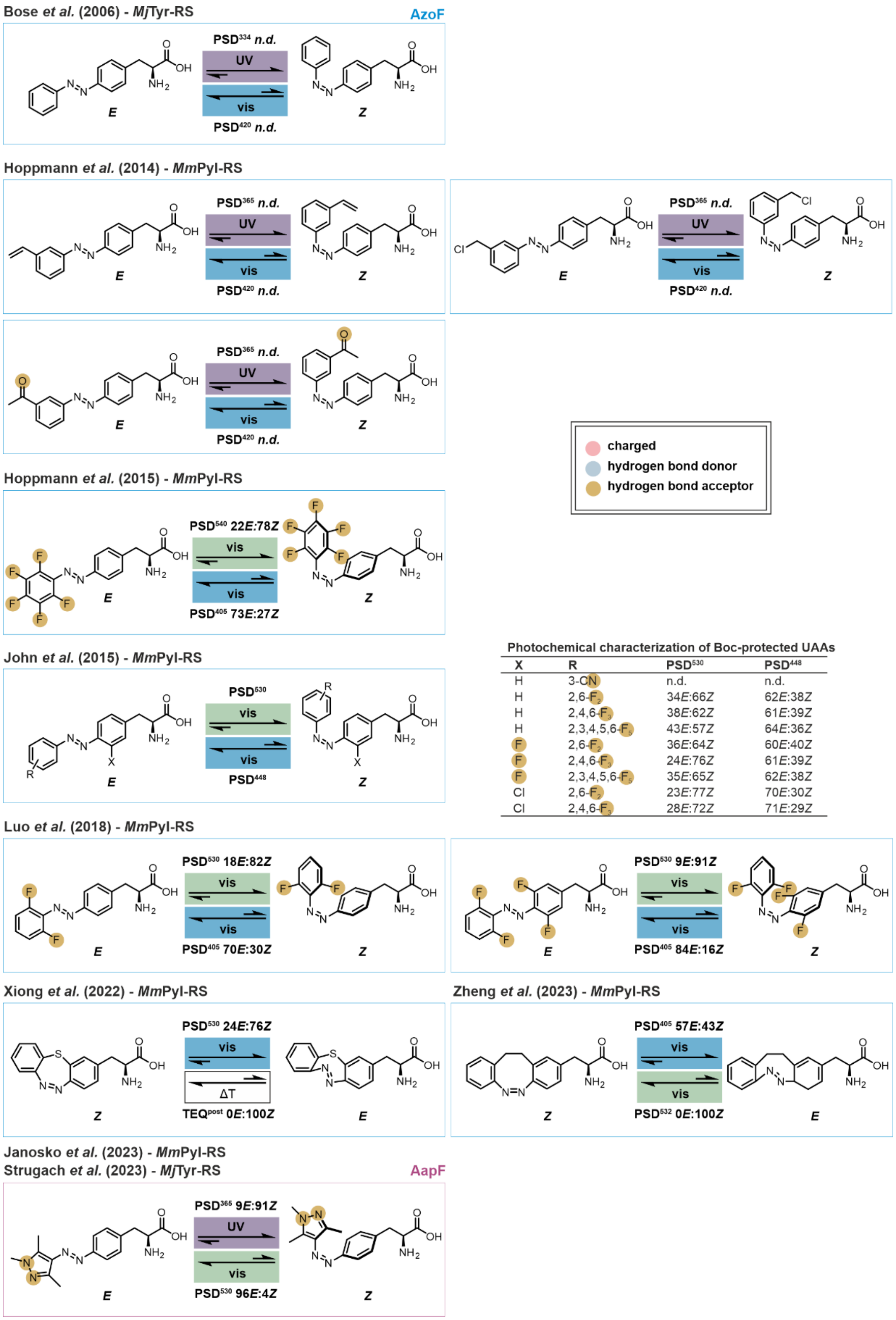

Photochemical characterization of Boc-protected UAAs

| X | R | PSD530 | PSD448 |
|---|---|---|---|
| H | 3-CN | n.d. | n.d. |
| H | 2,6-F2 | 34E:66Z | 62E:38Z |
| H | 2,4,6-F3 | 38E:62Z | 61E:39Z |
| H | 2,3,4,5,6-F5 | 43E:57Z | 64E:36Z |
| F | 2,6-F2 | 36E:64Z | 60E:40Z |
| F | 2,4,6-F3 | 24E:76Z | 61E:39Z |
| F | 2,3,4,5,6-F5 | 35E:65Z | 62E:38Z |
| Cl | 2,6-F2 | 23E:77Z | 70E:30Z |
| Cl | 2,4,6-F3 | 28E:72Z | 71E:29Z |

**Figure S1.** Hitherto available psUAAs and corresponding aaRSs as reported by Bose *et al.*,[1] Hoppmann *et al.*,[2] John *et al.*,[3] Luo *et al.*,[4] Xiong *et al.*,[5] Zheng *et al.*,[6] Janosko *et al.*,[7] and Strugach *et al.*.[8]

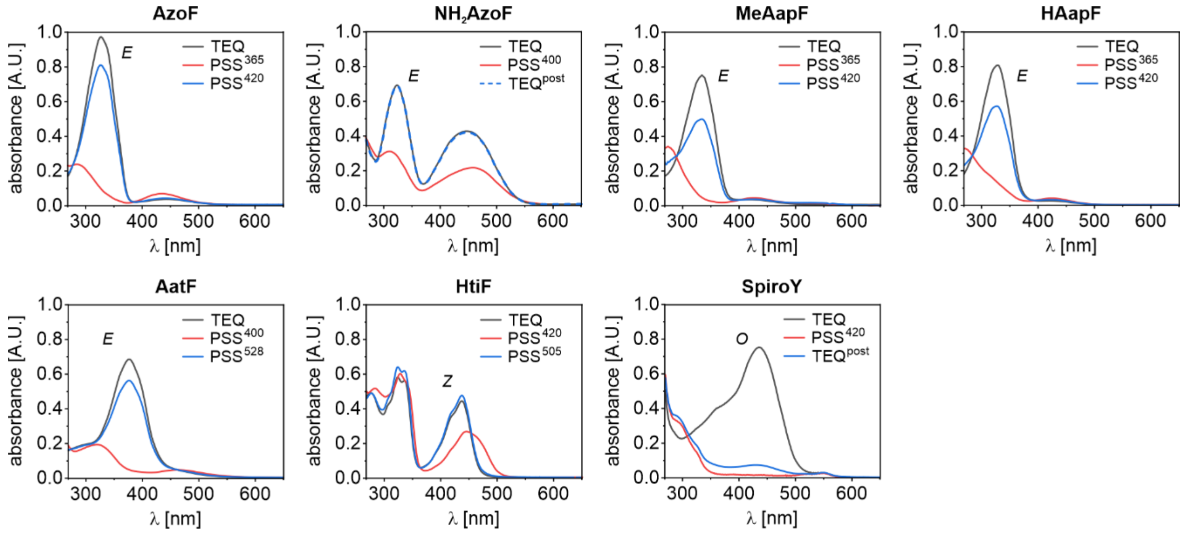

**Figure S2.** Absorbance spectra of the psUAAs in DMSO. Real-time tracking of the absorbance signal assured the complete establishment of the respective PSS$^{\lambda}$ or TEQ$^{post}$ prior to spectrum acquisition (**Table S3** and **Figure S7**). The most abundant isomer in the TEQ is indicated next to the grey spectrum.

**Table S1.** Absorbance maxima $\lambda_{max}$ of each psUAA in 50 mM Tris/HCl pH 7.5 and DMSO.

| | | buffer | | DMSO | |
|---|---|---|---|---|---|
| psUAA | isomer | $\lambda_{max}$[a] [nm] | $\lambda_{max}$[b] [nm] | $\lambda_{max}$[a] [nm] | $\lambda_{max}$[b] [nm] |
| **AzoF** | *E* | 324 | 423 | 329 | 446 |
| | *Z* | 299 | 427 | 287 | 437 |
| **NH2AzoF** | *E* | 323 | 404 | 325 | 449 |
| | *Z* | 314 | 443 | 311 | 461 |
| **MeAapF** | *E* | 333 | 419 | 335 | 426 |
| | *Z* | 278 | 417 | 275 | 427 |
| **HAapF** | *E* | 326 | 412 | 329 | 424 |
| | *Z* | 274 | 419 | 271 | 425 |
| **AatF** | *E* | 371 | —[c] | 378 | —[c] |
| | *Z* | 323 | 455 | 325 | 467 |
| **HtiF** | *E* | 328 | 446 | 329 | 438 |
| | *Z* | 333 | 441 | 330 | 446 |
| **SpiroY** | *O* | 377 | 530 | 438 | —[c] |
| | *C* | —[c] | —[c] | —[c] | —[c] |

[a] π→π* transition in case of diazo compounds. [b] n→π* transition in case of diazo compounds. [c] no spectroscopic signal.

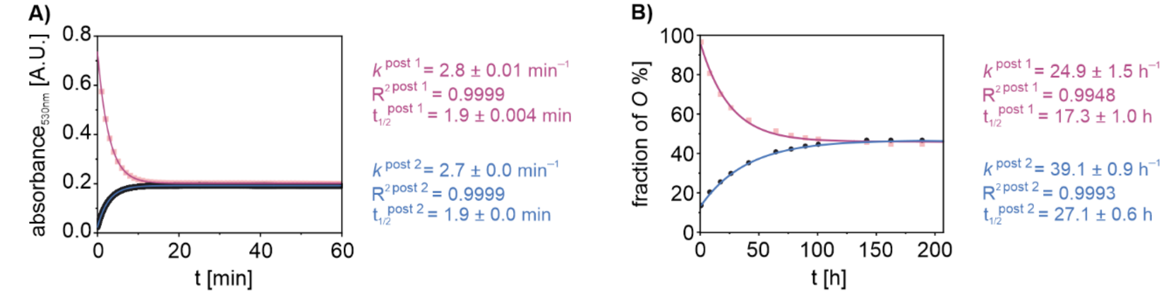

**Figure S3.** Establishment of TEQ$^{post1}$ and TEQ$^{post2}$ of **SpiroY** in 50 mM Tris/HCl pH 7.5 (A) or DMSO (B). TEQ$^{post1}$ is established after dilution of the **SpiroY** stock solution ($k^{post\ 1}$) and TEQ$^{post2}$ after thermal relaxation post irradiation with 455 nm ($k^{post\ 2}$). For the fast relaxation in buffer (A), the absorbance of *O* was followed over time. For the slow relaxation in DMSO (B), the isomer ratios at certain timepoints were determined via HPLC and the fraction of *O* was plotted against the time. Mono-exponential fitting yielded the rate constants *k*. Corresponding thermal half-lives $t_{1/2}$ were deduced from the respective *k* values.

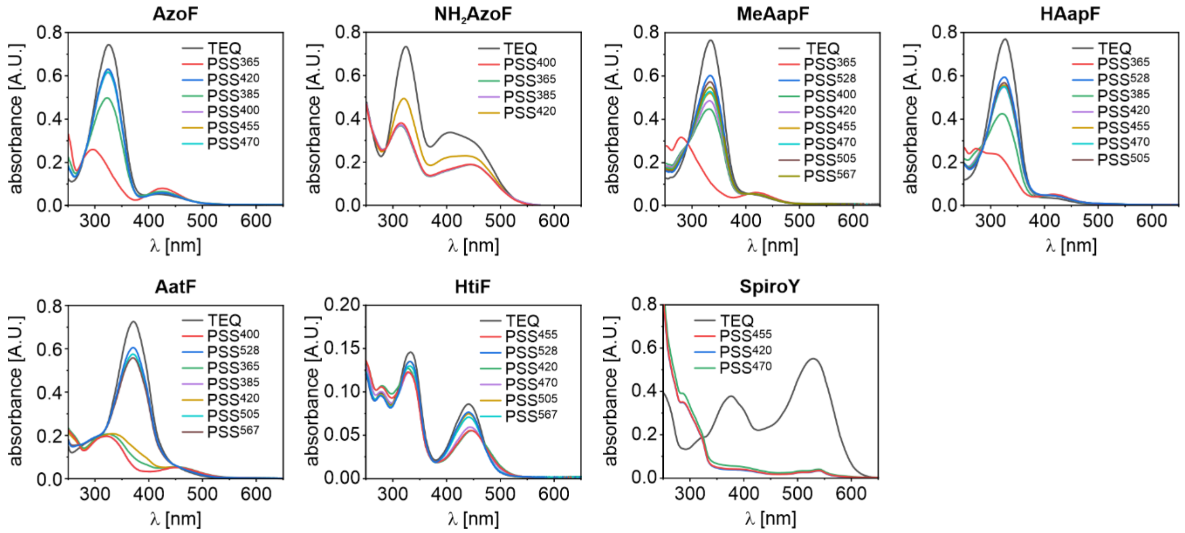

**Figure S4.** Absorbance spectra of each psUAA in 50 mM Tris/HCl pH 7.5 were acquired in the TEQ and in the PSS$^{\lambda}$ after irradiation with various wavelengths (*λ*). Real-time tracking of the absorbance signal assured the complete establishment of each PSS$^{\lambda}$ (**Table S2** and **Figure S6**). The most effective wavelengths, which induce the highest PSS for each isomer, were selected for further experiments. Note that for **$NH_2$AzoF** the *E*:*Z* distribution greatly depends on the intensity of irradiation owing to the fast thermal relaxation (cf. **Figure S6** and **Figure S8**). In this case, the most effective wavelength was 400 nm, because it induced the highest PSS with the lowest intensity of irradiation (**Table S14**).

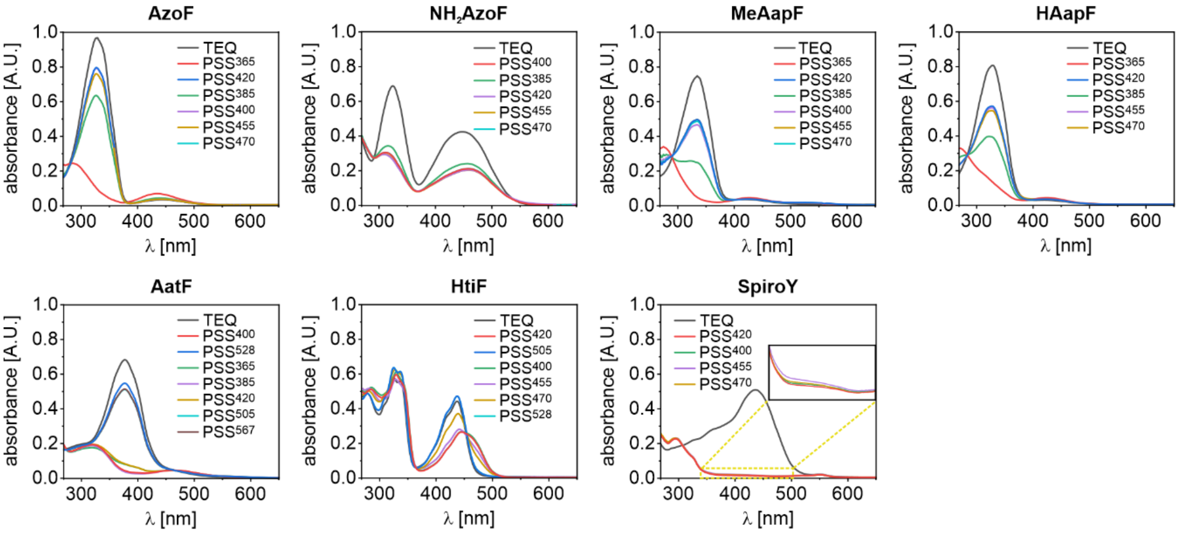

**Figure S5.** Absorbance spectra of each psUAA in DMSO were acquired in the TEQ and in the PSS$^{\lambda}$ after irradiation with various wavelengths ($\lambda$). Real-time tracking of the absorbance signal assured the complete establishment of each PSS$^{\lambda}$ (**Table S3** and **Figure S7**). The most effective wavelengths, which induce the highest PSS for each isomer, were selected for further experiments. Note that for **NH$_2$AzoF** the *E*:*Z* distribution greatly depends on the intensity of irradiation owing to the fast thermal relaxation (cf. **Figure S7**). In this case, the most effective wavelength was 400 nm, because it induced the highest PSS with the lowest intensity of irradiation (**Table S14**).

**Table S2.** Determination of isomerization half-lives ($t_{1/2}$) for each irradiation wavelength ($\lambda$) in 50 mM Tris/HCl pH 7.5.

| psUAA | $\lambda$[a] | $t_{1/2}$[b] [s] | psUAA | $\lambda$[a] | $t_{1/2}$[b] [s] |
|---|---|---|---|---|---|
| **AzoF** | 365 nm | 1.3 ± 0.006 | **AatF** | 365 nm | 0.4 ± 0.007 |
| | 385 nm | 3.0 ± 0.02 | | 385 nm | 0.6 ± 0.008 |
| | 400 nm | 1.8 ± 0.002 | | 400 nm | 0.6 ± 0.004 |
| | 420 nm | 0.9 ± 0.001 | | 420 nm | 0.6 ± 0.005 |
| | 455 nm | 1.0 ± 0.001 | | 505 nm | 5.5 ± 0.004 |
| | 470 nm | 1.8 ±0.001 | | 528 nm | 13.8 ± 0.01 |
| **$NH_2$AzoF** | 365 nm | 0.7 ± 0.001 | | 567 nm | 18.3 ± 0.01 |
| | 385 nm | 1.0 ± 0.001 | **HtiF** | 420 nm | 0.7 ± 0.003 |
| | 400 nm | 0.6 ± 0.002 | | 455 nm | 0.5 ± 0.002 |
| | 470 nm | 0.8 ± 0.0004 | | 470 nm | 0.8 ± 0.02 |
| **MeAapF** | 365 nm | 0.7 ± 0.003 | | 505 nm | 3.3 ± 0.02 |
| | 400 nm | 1.3 ± 0.001 | | 528 nm | 13.2 ± 0.02 |
| | 420 nm | 1.1 ± 0.001 | | 567 nm | 118.5 ± 0.02 |
| | 455 nm | 0.8 ± 0.003 | **SpiroY** | 420 nm | –[c] |
| | 470 nm | 1.8 ± 0.001 | | 455 nm | –[c] |
| | 505 nm | 32.5 ± 0.05 | | 470 nm | –[c] |
| | 528 nm | 129.1 ± 0.06 | | | |
| | 567 nm | 108.0 ± 0.03 | | | |
| **HAapF** | 365 nm | 1.2 ± 0.002 | | | |
| | 385 nm | 3.0 ± 0.004 | | | |
| | 420 nm | 1.03 ± 0.005 | | | |
| | 455 nm | 1.4 ± 0.005 | | | |
| | 470 nm | 2.5 ± 0.005 | | | |
| | 505 nm | 35.1 ± 0.03 | | | |
| | 528 nm | 164.7 ±0.04 | | | |

[a] The most effective wavelengths according to **Figure S4** are highlighted in red (first switch) and blue (back switch). [b] $t_{1/2}$ values were derived from the respective isomerization rates *k* (**Figure S6**). [c] Isomerization rates of **SpiroY** could not be determined because the establishment of TEQ[post 1] prevented an unambiguous fit.

**Table S3.** Determination of isomerization half-lives ($t_{½}$) for each irradiation wavelength ($\lambda$) in DMSO.

<table>
<tr><th>psUAA</th><th>$\lambda$[a]</th><th>$t_{½}$[b] [s]</th><th>psUAA</th><th>$\lambda$[a]</th><th>$t_{½}$[b] [s]</th></tr>
<tr><td>**AzoF**</td><td>365 nm</td><td>2.0 ± 0.003</td><td>**AatF**</td><td>365 nm</td><td>0.4 ± 0.006</td></tr>
<tr><td></td><td>385 nm</td><td>7.4 ± 0.0003</td><td></td><td>385 nm</td><td>0.5 ± 0.007</td></tr>
<tr><td></td><td>400 nm</td><td>3.1 ± 0.0004</td><td></td><td>400 nm</td><td>0.5 ± 0.005</td></tr>
<tr><td></td><td>420 nm</td><td>1.2 ± 0.0004</td><td></td><td>420 nm</td><td>0.4 ± 0.006</td></tr>
<tr><td></td><td>455 nm</td><td>0.9 ± 0.0003</td><td></td><td>505 nm</td><td>3.5 ± 0.002</td></tr>
<tr><td>**$NH_2$AzoF**</td><td>385 nm</td><td>1.4 ± 0.002</td><td></td><td>528 nm</td><td>8.6 ± 0.004</td></tr>
<tr><td></td><td>400 nm</td><td>0.9 ± 0.001</td><td></td><td>567 nm</td><td>11.6 ± 0.007</td></tr>
<tr><td></td><td>420 nm</td><td>0.3 ± 0.001</td><td>**HtiF**</td><td>400 nm</td><td>0.02 ± 0.001</td></tr>
<tr><td></td><td>455 nm</td><td>0.03 ± 0.001</td><td></td><td>420 nm[d,e]</td><td>0.35 ± 0.003 | 29.5 ± 1.47</td></tr>
<tr><td></td><td>470 nm</td><td>0.4 ± 0.005</td><td></td><td>455 nm</td><td>0.2 ± 0.0001</td></tr>
<tr><td>**MeAapF**</td><td>365 nm</td><td>0.5 ± 0.002</td><td></td><td>470 nm</td><td>0.6 ± 0.001</td></tr>
<tr><td></td><td>385 nm</td><td>2.0 ± 0.002</td><td></td><td>505 nm[d]</td><td>7.2 ± 0.007</td></tr>
<tr><td></td><td>400 nm</td><td>2.9 ± 0.001</td><td></td><td>528 nm</td><td>42.1 ± 0.02</td></tr>
<tr><td></td><td>420 nm</td><td>1.0 ± 0.002</td><td>**SpiroY**</td><td>420 nm[d]</td><td>–[c]</td></tr>
<tr><td></td><td>455 nm</td><td>0.7 ± 0.001</td><td></td><td>400 nm</td><td>–[c]</td></tr>
<tr><td></td><td>470 nm</td><td>2.4 ± 0.001</td><td></td><td>455 nm</td><td>–[c]</td></tr>
<tr><td>**HAapF**</td><td>365 nm</td><td>1.4 ± 0.003</td><td></td><td>470 nm</td><td>–[c]</td></tr>
<tr><td></td><td>385 nm</td><td>4.4 ± 0.006</td><td></td><td></td><td></td></tr>
<tr><td></td><td>420 nm</td><td>1.2 ± 0.004</td><td></td><td></td><td></td></tr>
<tr><td></td><td>455 nm</td><td>0.8 ± 0.007</td><td></td><td></td><td></td></tr>
<tr><td></td><td>470 nm</td><td>1.7 ± 0.003</td><td></td><td></td><td></td></tr>
</table>

[a] The most effective wavelengths according to **Figure S5** are highlighted in red (first switch) and blue (back switch). [b] $t_{½}$ values were derived from the respective isomerization rates *k* (**Figure S7**). [c] Isomerization rates of **SpiroY** could not be determined because the establishment of TEQ[post] [1] prevented an unambiguous fit. [d] Discrepancies were observed in the effective wavelengths in DMSO compared with the measurements in buffer. [e] Two $t_{½}$ values were obtained relating to the double-exponential fit for HtiF (**Figure S7**).

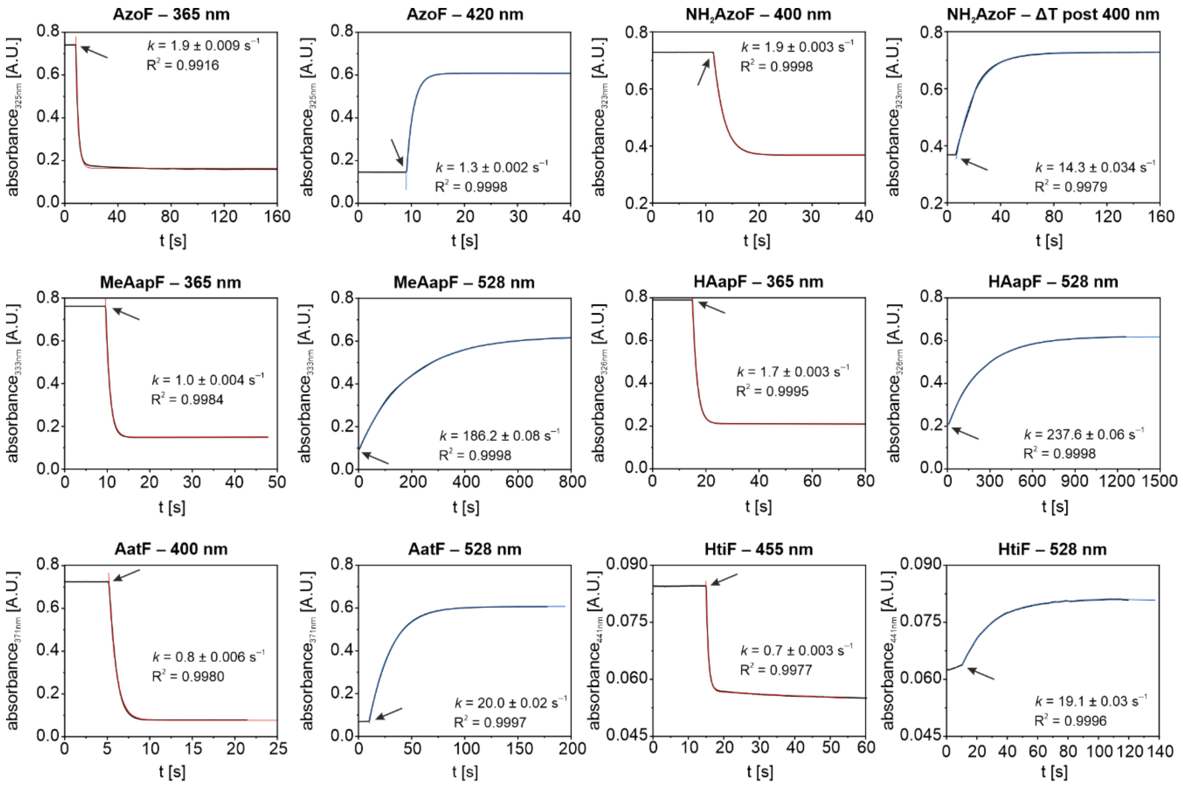

**Figure S6.** Exemplary isomerization rates of each psUAA in 50 mM Tris/HCl pH 7.5 upon irradiation with the most effective wavelengths (**Figure S4**) or through thermal relaxation (*ΔT*). The absorbance was followed at the π→π* or n→π* transition. Irradiation (arrow) was started and either maintained until a plateau was reached (for evaluation of isomerization upon irradiation) or stopped (for evaluation of thermal relaxation). The rates of isomerization *k* were determined by fitting the data with a mono-exponential function. Isomerization rates of **SpiroY** could not be determined because the establishment of $TEQ^{post}$ [1] prevented an unambiguous fit. Half-lives of isomerization $t_{1/2}$ were deduced from the respective *k* values and are listed in **Table S2**. The color code is according to **Figure 3** and **Table S2**.

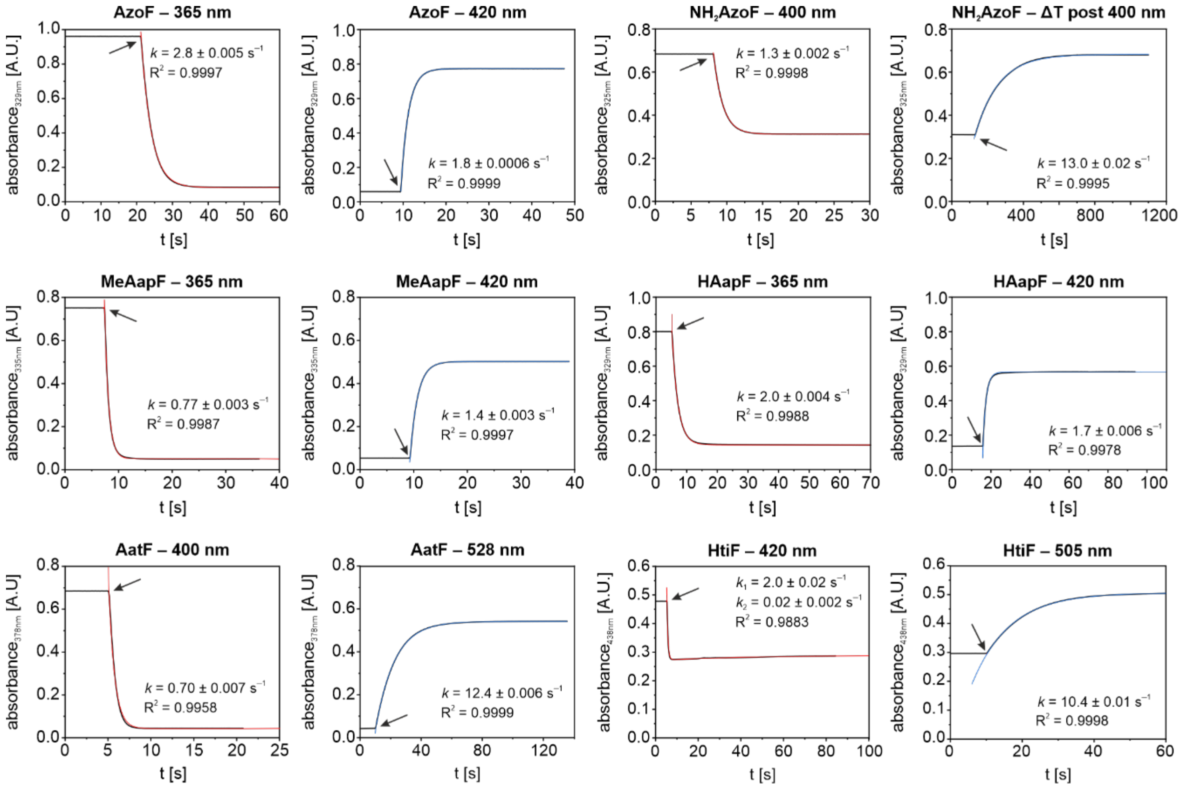

**Figure S7.** Exemplary isomerization rates of each psUAA in DMSO upon irradiation with the most effective wavelengths (**Figure S5**) or through thermal relaxation (*ΔT*). The absorbance was followed at the π→π* or n→π* transition. Irradiation (arrow) was started and either maintained until a plateau was reached (for evaluation of isomerization upon irradiation) or stopped (for evaluation of thermal relaxation). The rates of isomerization *k* were determined by fitting the data with a mono-exponential function. Isomerization rates of **SpiroY** could not be determined because the establishment of TEQ$^{post}$ [1] prevented an unambiguous fit. In case of **HtiF**, a double-exponential behavior was observed, presumably originating from the underlying isomerization mechanism of hemithioindigos comprising various transient intermediates.[9] Half-lives of isomerization $t_{½}$ were deduced from the respective *k* values and are listed in **Table S3**. The color code is according to **Figure S2** and **Table S3**.

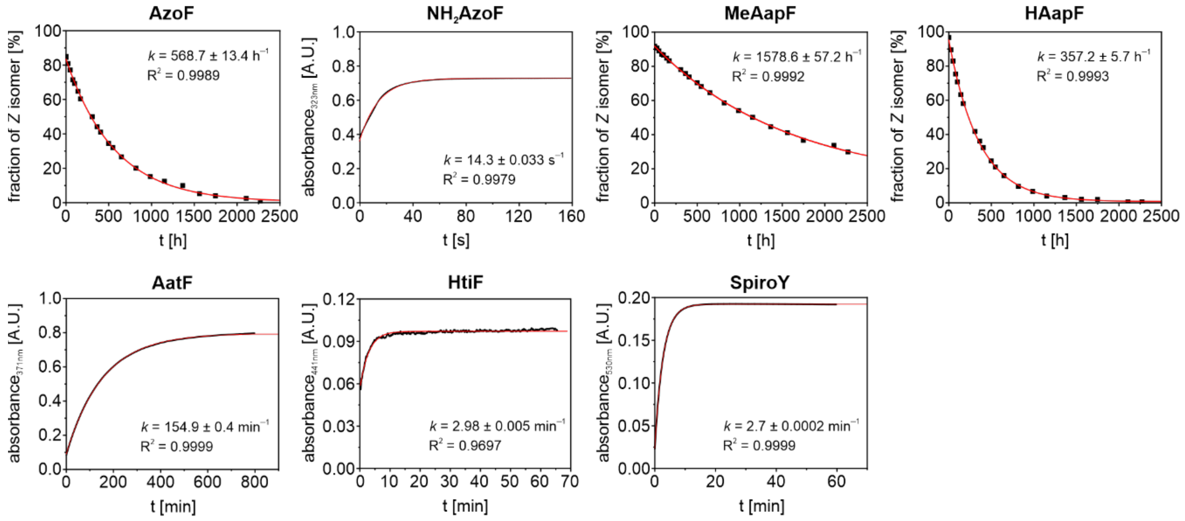

**Figure S8.** Thermal half-lives of each psUAA in 50 mM Tris/HCl pH 7.5 at 25 °C. Starting points of the measurements were the PSS$^{\lambda}$ enriched in the thermally less stable isomer: PSS$^{365}$ for **AzoF**, PSS$^{385}$ for **$NH_2$AzoF**, PSS$^{365}$ for **MeAapF**, PSS$^{340}$ for **HAapF**, PSS$^{400}$ for **AatF**, PSS$^{455}$ for **HtiF**, and PSS$^{455}$ for **SpiroY**. Note that we chose to irradiate **HAapF** with 340 nm, in contrast to all other experimental setups, since this wavelength achieved a remarkable PSS distribution of 9*E*:91*Z* (**Figure S9**). For psUAAs with a fast thermal relaxation (**$NH_2$AzoF**, **AatF**, **HtiF**, **SpiroY**), the absorbance at the π→π* or n→π* transition was followed over time. For psUAAs with longer rates of thermal isomerization (**AzoF**, **MeAapF**, **HAapF**), the isomer ratios at certain timepoints were determined via HPLC and the fraction of *Z* isomer was plotted against the time. Mono-exponential fitting yielded the rate constants of isomerization *k*. Corresponding thermal half-lives $t_{1/2}(\Delta T)$ were deduced from the respective *k* values.

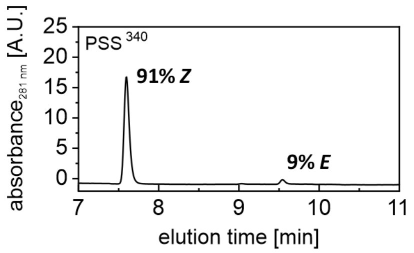

**Figure S9.** HPLC chromatogram of 20 µM **HAapF** in 50 mM Tris/HCl pH 7.5 to determine the isomer distribution in PSS$^{340}$. The absorbance was monitored at 281 nm and the isomer ratio was calculated from the peak areas. Irradiation at even shorter wavelengths (340 nm) induced a PSS with higher amount of *Z* isomer than observed for irradiation with 365 nm, however, it is not suited for most biological applications.

**Table S4.** Isosbestic points of psUAAs in 50 mM Tris/HCl pH 7.5 and DMSO.

| psUAA | Tris/HCl | DMSO |
|---|---|---|
| **AzoF** | **278 nm** | **279 nm** |
| | 391 nm | 386 nm |
| **$NH_2$AzoF** | **285 nm** | **291 nm** |
| | 525 nm | 539 nm |
| **MeAapF** | **290 nm** | **291 nm** |
| | 404 nm | 402 nm |
| **HAapF** | **281 nm** | **284 nm** |
| | 383 nm | 393 nm |
| **AatF** | **272 nm** | **275 nm** |
| | 456 nm | 464 nm |
| **HtiF** | 312 nm | **363 nm** |
| | 373 nm | 454 nm |
| | **471 nm** | |
| **SpiroY** | **306 nm** | **308 nm** |

Isosbestic points were determined from the respective UV/Vis spectra in 50 mM Tris/HCl pH 7.5 or DMSO (**Figure S4** and **Figure S5**). Isosbestic points that were used for HPLC measurements are marked in bold.

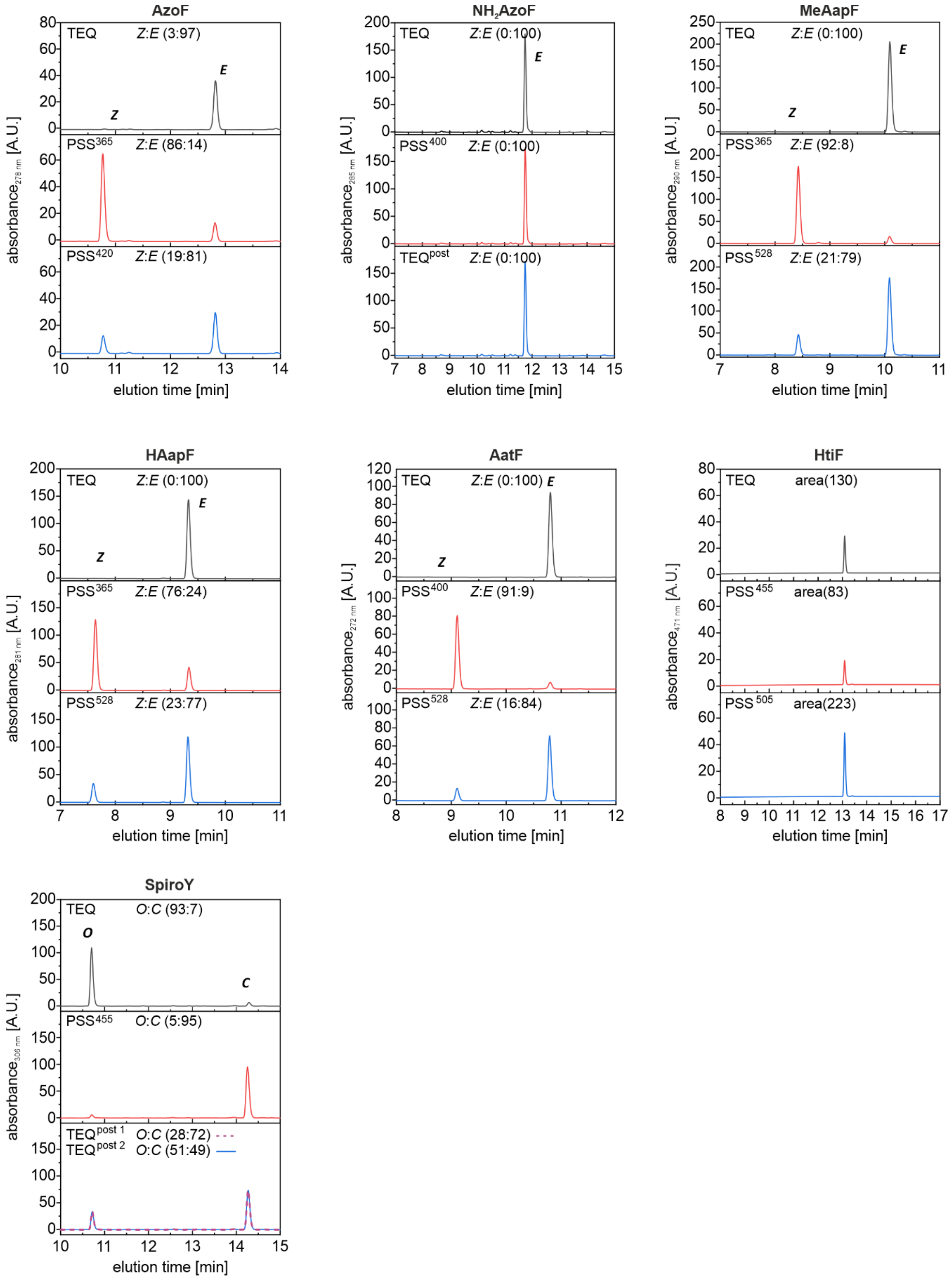

**Figure S10.** HPLC chromatograms of each psUAA in 50 mM Tris/HCl pH 7.5. The absorbance was monitored at certain isosbestic points (**Table S4**). Measurements of 400 µM **AzoF**, **NH$_2$AzoF**, **MeAapF**, **HAapF**, **AatF**, as well as 400 µM **HtiF**, and 400 µM **SpiroY** were performed in the thermal equilibrium (TEQ), after irradiation with the selected wavelengths (PSS$^{\lambda}$, **Table S2**), or after thermal relaxation (TEQ$^{post}$). The samples were irradiated right before the HPLC measurement and the column thermostat was set to 15 °C to prolong thermal relaxation of **NH$_2$AzoF**, **HtiF**, and **SpiroY**. Isomer ratios of each psUAA were determined by integration of the respective isomer peaks. For **NH$_2$AzoF**, thermal relaxation was still too fast and thus, 100% of the *E* isomer were regenerated before the measurement was finished. Only one peak was visible for **HtiF**, as well. However, we ruled out fast thermal relaxation, because the peak areas in TEQ, $PSS^{455}$, and $PSS^{505}$ mismatched.

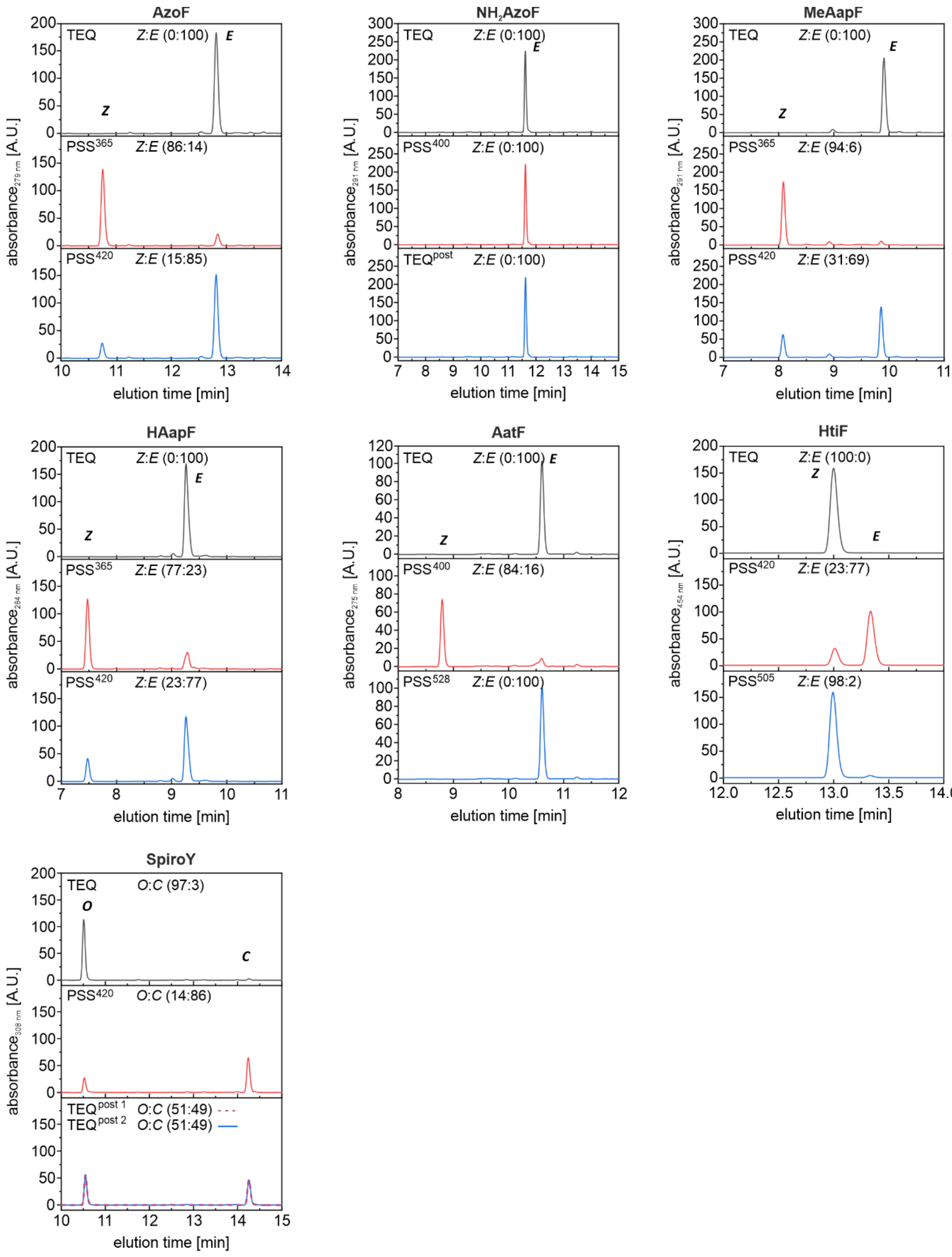

**Figure S11.** HPLC chromatograms of each psUAA in DMSO. The absorbance was monitored at certain isosbestic points (**Table S4**). Measurements of 400 µM AzoF, NH$_2$AzoF, MeAapF, HAapF, AatF, as well as 400 µM HtiF, and 400 µM SpiroY were performed in the thermal equilibrium (TEQ), after irradiation with the selected wavelengths ($PSS^{\lambda}$,

**Table S3**) or after thermal relaxation (TEQ$^{post}$). The samples were irradiated right before HPLC measurement. Isomer ratios of each psUAA were determined by integration of the respective isomer peaks. **For $NH_2$AzoF**, thermal relaxation was really fast and thus, 100% of the *E* isomer were regenerated before the measurement was finished.

**Table S5.** Determination of isomer ratios at the established PSS$^{\lambda}$ after irradiation or thermal isomerization in 50 mM Tris/HCl pH 7.5 determined by HPLC and UV/Vis measurements.

| UAA | | HPLC | | UV/Vis | |
|---|---|---|---|---|---|
| | | *E* | *Z* | *E* | *Z* |
| **AzoF** | TEQ | 96.6 | 3.40 | 100 | 0.00 |
| | PSS$^{365}$ | 13.9 | 86.1 | 15.0 | 85.0 |
| | PSS$^{420}$ | 80.7 | 19.3 | 79.0 | 21.0 |
| **$NH_2$AzoF** | TEQ | 100 | 0.00 | 100 | 0.00 |
| | PSS$^{400}$ | –[a] | –[a] | 45.5 | 54.5 |
| | TEQ$^{post}$ | 100 | 0.00 | 100 | 0.00 |
| **MeAapF** | TEQ | 100 | 0.00 | 100 | 0.00 |
| | PSS$^{365}$ | 8.3 | 91.7 | 9.50 | 90.5 |
| | PSS$^{420}$ | 63.0 | 37.0 | 70.8 | 29.2 |
| | PSS$^{528}$ | 79.3 | 20.7 | 77.3 | 22.7 |
| **HAapF** | TEQ | 100 | 0.00 | 100 | 0.00 |
| | PSS$^{365}$ | 24.2 | 75.8 | 24.0 | 76.0 |
| | PSS$^{505}$ | 73.0 | 27.0 | 61.3 | 38.7 |
| | PSS$^{528}$ | 77.3 | 22.7 | 77.5 | 22.5 |
| **AatF** | TEQ | 100 | 0.00 | 100 | 0.00 |
| | PSS$^{400}$ | 9.10 | 90.9 | 8.90 | 91.1 |
| | PSS$^{528}$ | 84.2 | 15.8 | 82.5 | 17.5 |
| **HtiF** | TEQ | –[b] | –[b] | –[c] | –[c] |
| | PSS$^{455}$ | –[b] | –[b] | –[c] | –[c] |
| | PSS$^{528}$ | –[b] | –[b] | –[c] | –[c] |
| | | *O* | *C* | *O* | *C* |
| **SpiroY** | TEQ | 93.4 | 6.60 | 100 | 0.00 |
| | PSS$^{455}$ | 5.40 | 94.6 | 3.50 | 96.5 |
| | TEQ$^{post\ 1}$ | 28.4 | 71.6 | 33.8 | 66.2 |
| | TEQ$^{post\ 2}$ | 28.4 | 71.6 | 33.8 | 66.2 |

The isomer ratios of each psUAA were determined in the thermal equilibrium (TEQ), as well as after irradiation with the selected wavelengths (PSS$^{\lambda}$, **Table S2**) or after thermal relaxation (TEQ$^{post}$). [a] *E*:*Z* ratio could not be determined since thermal isomerization was faster than the HPLC measurement. [b] isomer ratio could not be determined since only one isomer was visible in HPLC measurements. [c] *E*:*Z* ratio could not be determined since there is no method available.

**Table S6.** Determination of Isomer ratios at the established PSS$^{\lambda}$ after irradiation or thermal isomerization in DMSO determined by HPLC and UV/Vis measurements.

| UAA | | HPLC | | UV/Vis | |
|---|---|---|---|---|---|
| | | *E* | *Z* | *E* | *Z* |
| **AzoF** | TEQ | 100 | 0 | 100 | 0.00 |
| | PSS$^{365}$ | 13.7 | 86.3 | 5.40 | 94.6 |
| | PSS$^{420}$ | 85.2 | 14.8 | 82.4 | 17.6 |
| **NH$_2$AzoF** | TEQ | 100 | 0.00 | 100 | 0.00 |
| | PSS$^{400}$ | –[a] | –[a] | 37.2 | 62.8 |
| | TEQ$^{post}$ | 100 | 0.00 | 99.3 | 0.70 |
| **MeAapF** | TEQ | 100 | 0.00 | 100 | 0.00 |
| | PSS$^{365}$ | 6.00 | 94.0 | 4.30 | 95.7 |
| | PSS$^{420}$ | 69.0 | 31.0 | 63.9 | 36.1 |
| **HAapF** | TEQ | 100 | 0.00 | 100 | 0.00 |
| | PSS$^{365}$ | 23.1 | 76.9 | 13.3 | 86.7 |
| | PSS$^{420}$ | 76.9 | 23.1 | 69.6 | 30.4 |
| **AatF** | TEQ | 100 | 0.00 | 100 | 0.00 |
| | PSS$^{400}$ | 16.4 | 83.6 | 5.8 | 94.2 |
| | PSS$^{528}$ | 100 | 0.00 | 81.8 | 18.2 |
| **HtiF** | TEQ | 0.00 | 100 | –[b] | –[b] |
| | PSS$^{420}$ | 76.8 | 23.2 | –[b] | –[b] |
| | PSS$^{505}$ | 2.40 | 97.6 | –[b] | –[b] |
| | | *O* | *C* | *O* | *C* |
| **SpiroY** | TEQ | 97.2 | 2.80 | 100 | 0.00 |
| | PSS$^{420}$ | 14.0 | 86.0 | 1.90 | 98.1 |
| | TEQ$^{post\ 1}$ | 50.6 | 49.4 | –[c] | –[c] |
| | TEQ$^{post\ 2}$ | 50.6 | 49.4 | –[c] | –[c] |

The isomer ratios of each psUAA were determined in the thermal equilibrium (TEQ), after irradiation with the selected wavelengths (PSS$^{\lambda}$, **Table S2**) or after thermal relaxation (TEQ$^{post}$). [a] *E*:*Z* ratio could not be determined since thermal isomerization was faster than the HPLC measurement. [b] *E*:*Z* ratio could not be determined since there is no method available. [c] *E*:*Z* ratio was not determined owing to the slow establishment of TEQ$^{post1}$ and TEQ$^{post2}$ (**Figure S3**).

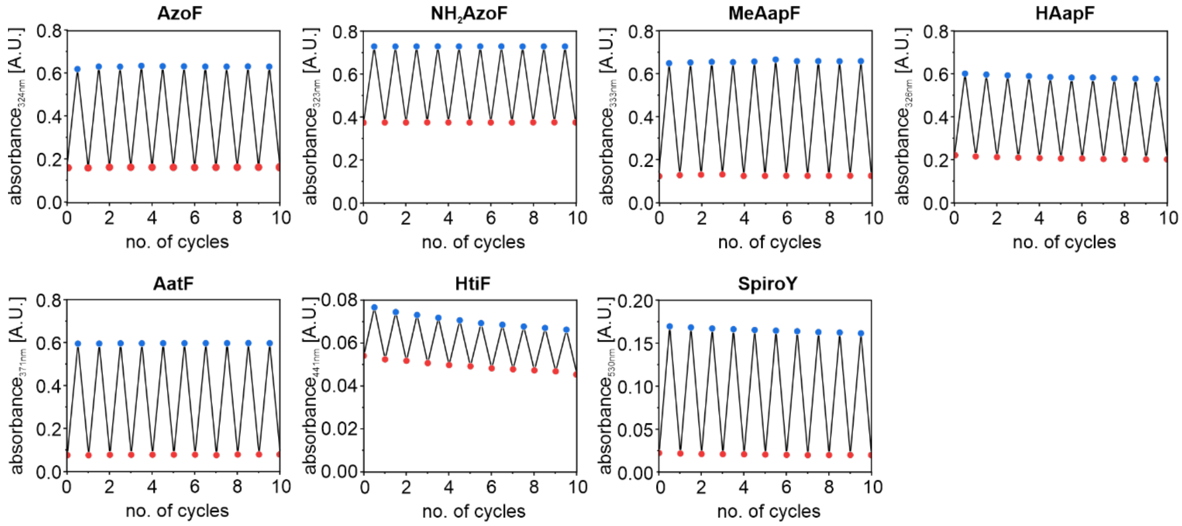

**Figure S12.** Cycle performance of the photoisomerization of each psUAA in 50 mM Tris/HCl pH 7.5. Alternating isomerization between $PSS^{365}$ and $PSS^{420}$ for **AzoF**, $PSS^{385}$ and $TEQ^{post}$ for **NH$_2$AzoF**, $PSS^{365}$ and $PSS^{528}$ for **MeAapF**, $PSS^{365}$ and $PSS^{528}$ for **HAapF**, $PSS^{400}$ and $PSS^{528}$ for **AatF**, $PSS^{455}$ and $PSS^{528}$ for **HtiF**, and $PSS^{455}$ and $TEQ^{post\ 2}$ for **SpiroY**. The absorbance at a characteristic maximum was plotted against the number of cycles (switching to *E* and subsequently to *Z* or inversely represents one cycle).

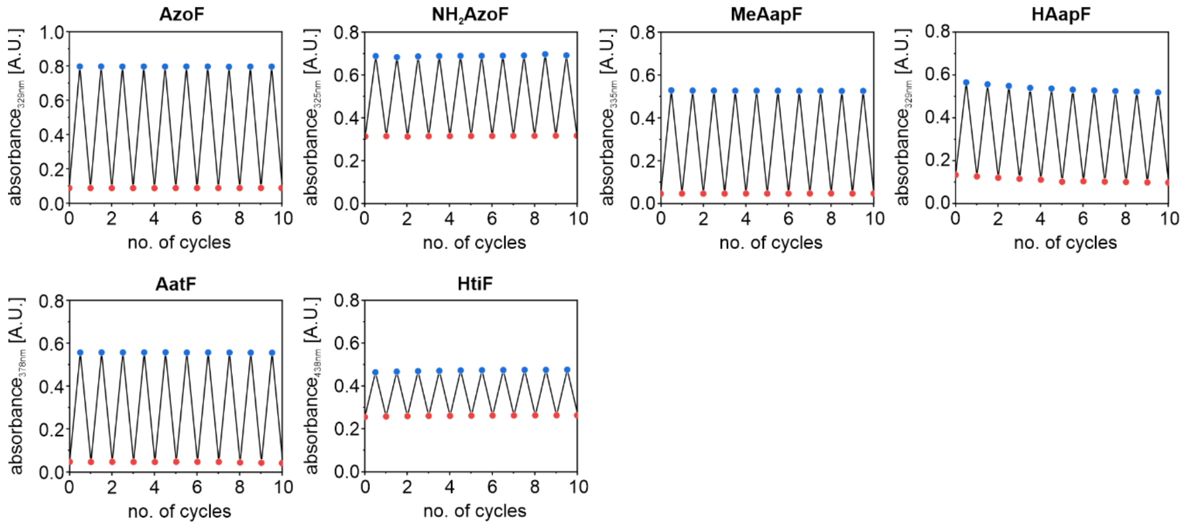

**Figure S13.** Cycle performance of the photoisomerization of each psUAA in DMSO. Alternating isomerization between $PSS^{365}$ and $PSS^{420}$ for **AzoF**, $PSS^{420}$ and $TEQ^{post}$ for **NH$_2$AzoF**, $PSS^{365}$ and $PSS^{420}$ for **MeAapF**, $PSS^{365}$ and $PSS^{420}$ for **HAapF**, $PSS^{400}$ and $PSS^{528}$ for **AatF**, and $PSS^{420}$ and $PSS^{505}$ for **HtiF**. The absorbance at a characteristic maximum was plotted against the number of cycles (switching to *E* and subsequently to *Z* or inversely represents one cycle). For **SpiroY**, evaluation of cycle performance was not feasible because there was no wavelength found that could induce ring-opening to switch back towards thermal equilibrium and the thermal half-life was rather long in DMSO.

**Table S7.** Identities and sequences of aaRSs tested for the incorporation of psUAAs.

| aaRS | aaRS | sequence |
|---|---|---|
| a | AzoF-RS[1] | MDEFEMIKRNTSEIISEEELREVLKKDEKSAGIGFEPSGKIHLGHYLQIKKMIDLQNAGFDIIIELA<br>DLHAYLNQKGELDEIRKIGDYNKKVFEAMGLKAKYVYGSEAELDKDYTLNVYRLALKTTLKRA<br>RRSMELIAREDENPKVAEVIYPIMQVNGIHYHGVDVAVGGMEQRKIHMLARELLPKKVVCIHN<br>PVLTGLDGEGKMSSSKGNFIAVDDSPEEIRAKIKKAYCPAGVVEGNPIMEIAKYFLEYPLTIKRP<br>EKFGGDLTVNSYEELESLFKNKELHPMRLKNAVAEELIKILEPIRKRL |
| b | AzoF-RS$_{copt}$[a] | MDEFEMIKRNTSEIISEEELREVLKKDEKSAGIGFEPSGKIHLGHYLQIKKMIDLQNAGFDIIIELA<br>DLHAYLNQKGELDEIRKIGDYNKKVFEAMGLKAKYVYGSEAELDKDYTLNVYRLALKTTLKRA<br>RRSMELIAREDENPKVAEVIYPIMQVNGIHYHGVDVAVGGMEQRKIHMLARELLPKKVVCIHN<br>PVLTGLDGEGKMSSSKGNFIAVDDSPEEIRAKIKKAYCPAGVVEGNPIMEIAKYFLEYPLTIKGP<br>EKFGGDLTVNSYEELESLFKNKELHPMRLKNAVAEELIKILEPIRKRL |
| c | ONBY-RS$_{i}$[10] | MDEFEMIKRNTSEIISEEELREVLKKDEKSAAIGFEPSGKIHLGHYLQIKKMIDLQNAGFDIIIALA<br>DLNAYLNQKGELDEIRKIGDYNKKVFEAMGLKAKYVYQSEFALDKDYTLNVYRLALKTTLKRA<br>RRSMELIAREDENPKVAEVIYPIMQVNSAHYAGVDVSVGGMEQRKIHMLQRELLPKKVVCIHN<br>PVLTGLDGEGKMSSSKGNFIAVDDSPEEIRAKIKKAYCPAGVVEGNPIMEIAKYFLEYPLTIKRP<br>EKFGGDLTVNSYEELESLFKNKELHPMRLKNAVAEELIKILEPIRKRL |
| d | ONBY-RS$_{i}$ _R257G | MDEFEMIKRNTSEIISEEELREVLKKDEKSAAIGFEPSGKIHLGHYLQIKKMIDLQNAGFDIIIALA<br>DLNAYLNQKGELDEIRKIGDYNKKVFEAMGLKAKYVYQSEFALDKDYTLNVYRLALKTTLKRA<br>RRSMELIAREDENPKVAEVIYPIMQVNSAHYAGVDVSVGGMEQRKIHMLQRELLPKKVVCIHN<br>PVLTGLDGEGKMSSSKGNFIAVDDSPEEIRAKIKKAYCPAGVVEGNPIMEIAKYFLEYPLTIKGP<br>EKFGGDLTVNSYEELESLFKNKELHPMRLKNAVAEELIKILEPIRKRL |
| e | AzF-RS[11] | MDEFEMIKRNTSEIISEEELREVLKKDEKSATIGFEPSGKIHLGHYLQIKKMIDLQNAGFDIIILLA<br>DLHAYLNQKGELDEIRKIGDYNKKVFEAMGLKAKYVYGSNFQLDKDYTLNVYRLALKTTLKRA<br>RRSMELIAREDENPKVAEVIYPIMQVNPLHYQGVDVAVGGMEQRKIHMLARELLPKKVVCIHN<br>PVLTGLDGEGKMSSSKGNFIAVDDSPEEIRAKIKKAYCPAGVVEGNPIMEIAKYFLEYPLTIKRP<br>EKFGGDLTVNSYEELESLFKNKELHPMRLKNAVAEELIKILEPIRKRL |
| f | Azo-RS 1[12] | MDEFEMIKRNTSEIISEEELREVLKKDEKSAGIGFEPSGKIHLGHYLQIKKMIDLQNAGFDIIIVLA<br>DLHAYLNQKGELDEIRKIGDYNKKVFEAMGLKAKYVYGSEFQLDKDYTLNVYRLALKTTLKRA<br>RRSMELIAREDENPKVAEVIYPIMQVNGYHYSGVDVFVGGMEQRKIHMLARELLPKKVVCIHN<br>PVLTGLDGEGKMSSSKGNFIAVDDSPEEIRAKIKKAYCPAGVVEGNPIMEIAKYFLEYPLTIKGP<br>EKFGGDLTVNSYEELESLFKNKELHPMRLKNAVAEELIKILEPIRKRL |
| g | Azo-RS 4[12] | MDEFEMIKRNTSEIISEEELREVLKKDEKSAGIGFEPSGKIHLGHYLQIKKMIDLQNAGFDIIIVLA<br>DLHAYLNQKGELDEIRKIGDYNKKVFEAMGLKAKYVYGSEFQLDKDYTLNVYRLALKTTLKRA<br>RRSMELIAREDENPKVAEVIYPIMQVNGYHYRGVDVAVGGMEQRKIHMLARELLPKKVVCIHN<br>PVLTGLDGEGKMSSSKGNFIAVDDSPEEIRAKIKKAYCPAGVVEGNPIMEIAKYFLEYPLTIKGP<br>EKFGGDLTVNSYEELESLFKNKELHPMRLKNAVAEELIKILEPIRKRL |
| h | AapF-RS | MDEFEMIKRNTSEIISEEELREVLKKDEKSAAIGFEPSGKIHLGHYLQIKKMIDLQNAGFDIIILLA<br>DLHAYLNQKGELDEIRKIGDYNKKVFEAMGLKAKYVYGSEFMLDKDYTLNVYRLALKTTLKRA<br>RRSMELIAREDENPKVAEVIYPIMQVNAIHYFGVDVAVGGMEQRKIHMLARELLPKKVVCIHNP<br>VLTGLDGEGKMSSSKGNFIAVDDSPEEIRAKIKKAYCPAGVVEGNPIMEIAKYFLEYPLTIKGPE<br>KFGGDLTVNSYEELESLFKNKELHPMRLKNAVAEELIKILEPIRKRL |
| i | AzoF-RS$_{copt}$_Q155V | MDEFEMIKRNTSEIISEEELREVLKKDEKSAGIGFEPSGKIHLGHYLQIKKMIDLQNAGFDIIIELA<br>DLHAYLNQKGELDEIRKIGDYNKKVFEAMGLKAKYVYGSEAELDKDYTLNVYRLALKTTLKRA<br>RRSMELIAREDENPKVAEVIYPIMVVNGIHYHGVDVAVGGMEQRKIHMLARELLPKKVVCIHN<br>PVLTGLDGEGKMSSSKGNFIAVDDSPEEIRAKIKKAYCPAGVVEGNPIMEIAKYFLEYPLTIKGP<br>EKFGGDLTVNSYEELESLFKNKELHPMRLKNAVAEELIKILEPIRKRL |
| j | AzoF-RS$_{copt}$_A167V | MDEFEMIKRNTSEIISEEELREVLKKDEKSAGIGFEPSGKIHLGHYLQIKKMIDLQNAGFDIIIELA<br>DLHAYLNQKGELDEIRKIGDYNKKVFEAMGLKAKYVYGSEAELDKDYTLNVYRLALKTTLKRA<br>RRSMELIAREDENPKVAEVIYPIMQVNGIHYHGVDVVVGGMEQRKIHMLARELLPKKVVCIHN<br>PVLTGLDGEGKMSSSKGNFIAVDDSPEEIRAKIKKAYCPAGVVEGNPIMEIAKYFLEYPLTIKGP<br>EKFGGDLTVNSYEELESLFKNKELHPMRLKNAVAEELIKILEPIRKRL |
| k | AzoF-RS$_{copt}$_A167C | MDEFEMIKRNTSEIISEEELREVLKKDEKSAGIGFEPSGKIHLGHYLQIKKMIDLQNAGFDIIIELA<br>DLHAYLNQKGELDEIRKIGDYNKKVFEAMGLKAKYVYGSEAELDKDYTLNVYRLALKTTLKRA<br>RRSMELIAREDENPKVAEVIYPIMQVNGIHYHGVDVCVGGMEQRKIHMLARELLPKKVVCIHN<br>PVLTGLDGEGKMSSSKGNFIAVDDSPEEIRAKIKKAYCPAGVVEGNPIMEIAKYFLEYPLTIKGP<br>EKFGGDLTVNSYEELESLFKNKELHPMRLKNAVAEELIKILEPIRKRL |
| l | AzoF-RS$_{copt}$_G32A | MDEFEMIKRNTSEIISEEELREVLKKDEKSAAIGFEPSGKIHLGHYLQIKKMIDLQNAGFDIIIELA<br>DLHAYLNQKGELDEIRKIGDYNKKVFEAMGLKAKYVYGSEAELDKDYTLNVYRLALKTTLKRA<br>RRSMELIAREDENPKVAEVIYPIMQVNGIHYHGVDVAVGGMEQRKIHMLARELLPKKVVCIHN<br>PVLTGLDGEGKMSSSKGNFIAVDDSPEEIRAKIKKAYCPAGVVEGNPIMEIAKYFLEYPLTIKGP<br>EKFGGDLTVNSYEELESLFKNKELHPMRLKNAVAEELIKILEPIRKRL |
| m | AzoF-RS$_{copt}$_E65L | MDEFEMIKRNTSEIISEEELREVLKKDEKSAGIGFEPSGKIHLGHYLQIKKMIDLQNAGFDIIILLA<br>DLHAYLNQKGELDEIRKIGDYNKKVFEAMGLKAKYVYGSEAELDKDYTLNVYRLALKTTLKRA<br>RRSMELIAREDENPKVAEVIYPIMQVNGIHYHGVDVAVGGMEQRKIHMLARELLPKKVVCIHN<br>PVLTGLDGEGKMSSSKGNFIAVDDSPEEIRAKIKKAYCPAGVVEGNPIMEIAKYFLEYPLTIKGP<br>EKFGGDLTVNSYEELESLFKNKELHPMRLKNAVAEELIKILEPIRKRL |

| | | |
|---|---|---|
| n | AzoF-RS$_{copt}$_V188P | MDEFEMIKRNTSEIISEEELREVLKKDEKSAGIGFEPSGKIHLGHYLQIKKMIDLQNAGFDIIIELA<br>DLHAYLNQKGELDEIRKIGDYNKKVFEAMGLKAKYVYGSEAELDKDYTLNVYRLALKTTLKRA<br>RRSMELIAREDENPKVAEVIYPIMQVNGIHYHGVDVAVGGMEQRKIHMLARELLPKKPVCIHN<br>PVLTGLDGEGKMSSSKGNFIAVDDSPEEIRAKIKKAYCPAGVVEGNPIMEIAKYFLEYPLTIKGP<br>EKFGGDLTVNSYEELESLFKNKELHPMRLKNAVAEELIKILEPIRKRL |
| o | AzoF-RS$_{copt}$_A108F | MDEFEMIKRNTSEIISEEELREVLKKDEKSAGIGFEPSGKIHLGHYLQIKKMIDLQNAGFDIIIELA<br>DLHAYLNQKGELDEIRKIGDYNKKVFEAMGLKAKYVYGSEFELDKDYTLNVYRLALKTTLKRA<br>RRSMELIAREDENPKVAEVIYPIMQVNGIHYHGVDVAVGGMEQRKIHMLARELLPKKVVCIHN<br>PVLTGLDGEGKMSSSKGNFIAVDDSPEEIRAKIKKAYCPAGVVEGNPIMEIAKYFLEYPLTIKGP<br>EKFGGDLTVNSYEELESLFKNKELHPMRLKNAVAEELIKILEPIRKRL |
| p | AzoF-<br>RS$_{copt}$_E109M | MDEFEMIKRNTSEIISEEELREVLKKDEKSAGIGFEPSGKIHLGHYLQIKKMIDLQNAGFDIIIELA<br>DLHAYLNQKGELDEIRKIGDYNKKVFEAMGLKAKYVYGSEAMLDKDYTLNVYRLALKTTLKRA<br>RRSMELIAREDENPKVAEVIYPIMQVNGIHYHGVDVAVGGMEQRKIHMLARELLPKKVVCIHN<br>PVLTGLDGEGKMSSSKGNFIAVDDSPEEIRAKIKKAYCPAGVVEGNPIMEIAKYFLEYPLTIKGP<br>EKFGGDLTVNSYEELESLFKNKELHPMRLKNAVAEELIKILEPIRKRL |
| q | AzoF-RS$_{copt}$_G158A | MDEFEMIKRNTSEIISEEELREVLKKDEKSAGIGFEPSGKIHLGHYLQIKKMIDLQNAGFDIIIELA<br>DLHAYLNQKGELDEIRKIGDYNKKVFEAMGLKAKYVYGSEAELDKDYTLNVYRLALKTTLKRA<br>RRSMELIAREDENPKVAEVIYPIMQVNAIHYHGVDVAVGGMEQRKIHMLARELLPKKVVCIHN<br>PVLTGLDGEGKMSSSKGNFIAVDDSPEEIRAKIKKAYCPAGVVEGNPIMEIAKYFLEYPLTIKGP<br>EKFGGDLTVNSYEELESLFKNKELHPMRLKNAVAEELIKILEPIRKRL |
| r | AzoF-RS$_{copt}$_H162F | MDEFEMIKRNTSEIISEEELREVLKKDEKSAGIGFEPSGKIHLGHYLQIKKMIDLQNAGFDIIIELA<br>DLHAYLNQKGELDEIRKIGDYNKKVFEAMGLKAKYVYGSEAELDKDYTLNVYRLALKTTLKRA<br>RRSMELIAREDENPKVAEVIYPIMQVNGIHYFGVDVAVGGMEQRKIHMLARELLPKKVVCIHN<br>PVLTGLDGEGKMSSSKGNFIAVDDSPEEIRAKIKKAYCPAGVVEGNPIMEIAKYFLEYPLTIKGP<br>EKFGGDLTVNSYEELESLFKNKELHPMRLKNAVAEELIKILEPIRKRL |
| s | AzoF-RS$_{copt}$_I159Y | MDEFEMIKRNTSEIISEEELREVLKKDEKSAGIGFEPSGKIHLGHYLQIKKMIDLQNAGFDIIIELA<br>DLHAYLNQKGELDEIRKIGDYNKKVFEAMGLKAKYVYGSEAELDKDYTLNVYRLALKTTLKRA<br>RRSMELIAREDENPKVAEVIYPIMQVNGYHYHGVDVAVGGMEQRKIHMLARELLPKKVVCIHN<br>PVLTGLDGEGKMSSSKGNFIAVDDSPEEIRAKIKKAYCPAGVVEGNPIMEIAKYFLEYPLTIKGP<br>EKFGGDLTVNSYEELESLFKNKELHPMRLKNAVAEELIKILEPIRKRL |
| t | AzoF-RS$_{copt}$_I159K | MDEFEMIKRNTSEIISEEELREVLKKDEKSAGIGFEPSGKIHLGHYLQIKKMIDLQNAGFDIIIELA<br>DLHAYLNQKGELDEIRKIGDYNKKVFEAMGLKAKYVYGSEAELDKDYTLNVYRLALKTTLKRA<br>RRSMELIAREDENPKVAEVIYPIMQVNGKHYHGVDVAVGGMEQRKIHMLARELLPKKVVCIHN<br>PVLTGLDGEGKMSSSKGNFIAVDDSPEEIRAKIKKAYCPAGVVEGNPIMEIAKYFLEYPLTIKGP<br>EKFGGDLTVNSYEELESLFKNKELHPMRLKNAVAEELIKILEPIRKRL |
| u | AzoF-RS$_{copt}$_I159F | MDEFEMIKRNTSEIISEEELREVLKKDEKSAGIGFEPSGKIHLGHYLQIKKMIDLQNAGFDIIIELA<br>DLHAYLNQKGELDEIRKIGDYNKKVFEAMGLKAKYVYGSEAELDKDYTLNVYRLALKTTLKRA<br>RRSMELIAREDENPKVAEVIYPIMQVNGFHYHGVDVAVGGMEQRKIHMLARELLPKKVVCIHN<br>PVLTGLDGEGKMSSSKGNFIAVDDSPEEIRAKIKKAYCPAGVVEGNPIMEIAKYFLEYPLTIKGP<br>EKFGGDLTVNSYEELESLFKNKELHPMRLKNAVAEELIKILEPIRKRL |
| v | AzoF-RS$_{copt}$_I159W | MDEFEMIKRNTSEIISEEELREVLKKDEKSAGIGFEPSGKIHLGHYLQIKKMIDLQNAGFDIIIELA<br>DLHAYLNQKGELDEIRKIGDYNKKVFEAMGLKAKYVYGSEAELDKDYTLNVYRLALKTTLKRA<br>RRSMELIAREDENPKVAEVIYPIMQVNGWHYHGVDVAVGGMEQRKIHMLARELLPKKVVCIH<br>NPVLTGLDGEGKMSSSKGNFIAVDDSPEEIRAKIKKAYCPAGVVEGNPIMEIAKYFLEYPLTIKG<br>PEKFGGDLTVNSYEELESLFKNKELHPMRLKNAVAEELIKILEPIRKRL |

[a] aaRS-b is a codon-optimized version (see **Table S8**) of aaRS-a harboring the R257G mutation that improves tRNA binding.[12]

**Table S8.** DNA sequence of the codon-optimized AzoF-RS$_{copt}$ (aaRS-b).

| | |
|---|---|
| AzoF-RS$_{copt}$ | AAAAAGGTCTCACATGGATGAGTTCGAAATGATTAAACGTAACACCAGCGAAATCATCAGCGAAGAAGAACTGCGCGAAGT<br>ACTGAAAAAGGATGAAAAATCAGCCGGTATTGGTTTTGAACCGAGCGGTAAAATTCATCTGGGTCATTATCTGCAAATCAAA<br>AAGATGATCGATCTGCAGAATGCCGGTTTCGATATTATCATTGAACTGGCCGATCTGCATGCATATCTGAATCAGAAAGGTG<br>AACTGGATGAAATTCGCAAAATCGGCGATTACAACAAAAAGGTGTTTGAAGCAATGGGCCTGAAAGCCAAATATGTTTATGG<br>TAGCGAAGCCGAGCTGGATAAAGATTATACCCTGAATGTTTATCGTCTGGCCCTGAAAACCACACTGAAACGTGCCCGTCG<br>TAGCATGGAACTGATTGCACGTGAAGATGAAAATCCGAAAGTTGCCGAAGTGATTTATCCGATTATGCAGGTTAACGGCATT<br>CATTATCATGGTGTTGATGTTGCAGTTGGTGGTATGGAACAGCGCAAAATTCATATGCTGGCACGTGAACTGCTGCCGAAG<br>AAAGTTGTTTGTATTCATAATCCGGTTCTGACCGGTCTGGATGGTGAAGGCAAAATGAGCAGCAGCAAAGGTAACTTTATTG<br>CCGTTGATGATAGTCCGGAAGAAATTCGTGCCAAAATCAAGAAAGCATATTGTCCGGCAGGCGTTGTTGAAGGTAACCCGA<br>TTATGGAAATTGCCAAATACTTTCTGGAATACCCGCTGACCATTAAAGGACCGGAAAAATTTGGTGGTGATCTGACCGTTAA<br>TAGCTATGAAGAACTGGAAAGCCTGTTTAAAAACAAAGAACTGCATCCGATGCGTCTGAAAAATGCAGTTGCGGAAGAACT<br>GATTAAAATCCTGGAACCGATTCGTAAACGTCTGCTCGAGAGACCAAAAA |

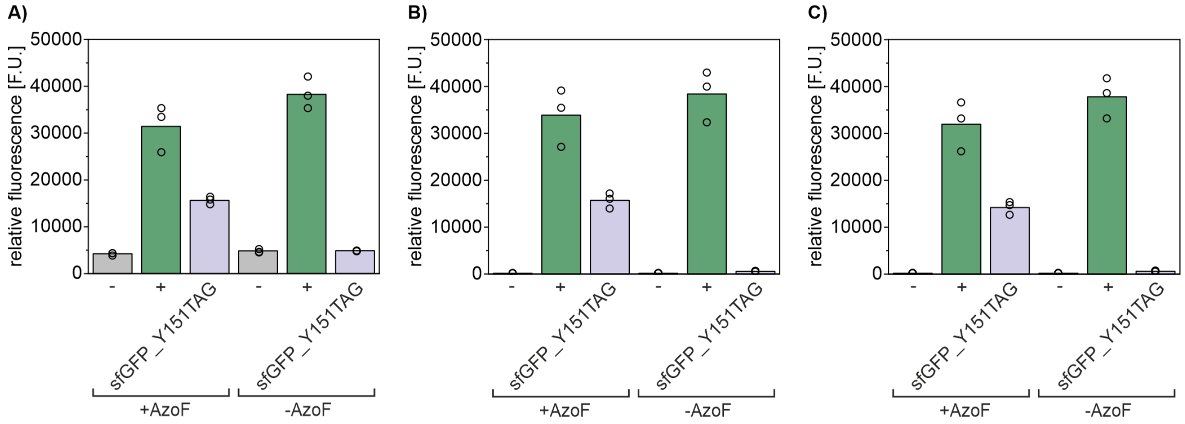

**Figure S14.** Optimization of the aaRS screening procedure. Different conditions for the treatment of the cells before sfGFP measurement were investigated. Relative fluorescence was measured right after expression in the presence and absence of **AzoF** overnight (A), after subsequent centrifugation and resuspension in 50 mM Tris/HCl pH 7.5 (B), and after additional cell lysis with 1x FastBreak$^{TM}$ Cell Lysis Reagent (C). The empty vector pETBAD (corresponding to a former pET24a vector where the arabinose inducible pBAD promoter was inserted) in combination with pGLNS_AzoF-RS$_{copt}$ served as negative control (−), pETBAD_sfGFP_wt in combination with pGLNS_AzoF-RS$_{copt}$ as positive control (+). The sfGFP_Y151TAG gene was co-expressed with pGLNS_AzoF-RS$_{copt}$. Bars represent the mean of biological triplicates with the single values depicted as circles. Depending on the treatment, significant differences in the background fluorescence were observed. At least cell lysis (B) was necessary to sufficiently reduce background fluorescence in the absence of **AzoF**.

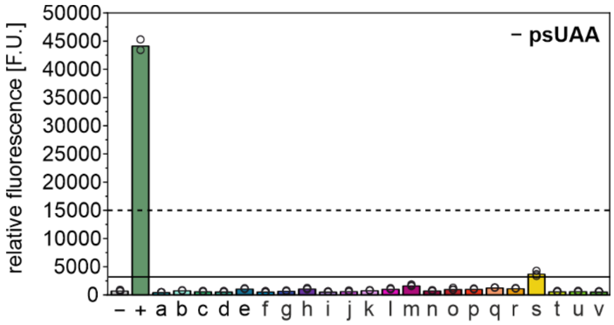

**Figure S15.** Evaluation of misincorporation of natural amino acids in the absence of a psUAA with *Mj*Tyr-RS variants a–v. Orthogonality could be confirmed for each aaRS except aaRS-s. Empty circles: fluorescence intensities of three biological replicates; columns: mean fluorescence intensity; solid line: lower threshold corresponding to the fluorescence intensity with the original AzoF-RS (aaRS-a); dashed line: upper threshold corresponding to ~30% of the wild type sfGFP fluorescence signal (**Figure 4A**).

**Table S9.** Successful psUAA-aaRS combinations found in the synthetase screening.

| psUAA to be incorporated | suitable synthetase as identified from screening |
|---|---|
| **AzoF** | aaRS-g in pGLNS |
| **$NH_2$AzoF** | aaRS-g in pGLNS |
| **MeAapF** | aaRS-p in pGLNS |
| **HAapF** | aaRS-g in pEVOL |
| **AatF** | aaRS-q in pGLNS |
| **HtiF** | aaRS-g in pGLNS |

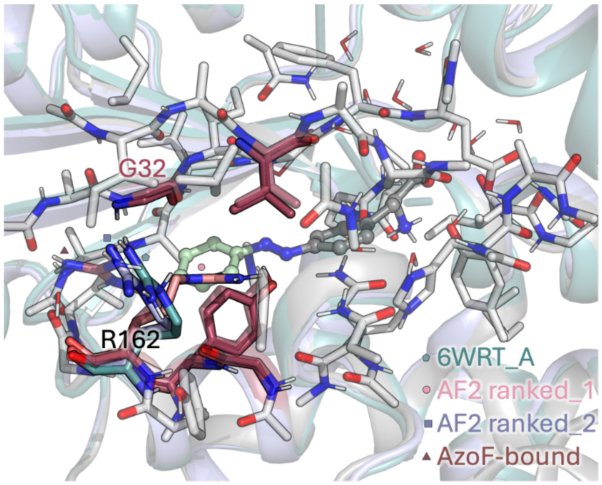

**Figure S16.** Optimized cluster model of aaRS-g with **AzoF** bound in the active site (**AzoF** is shown in gray and green spheres and sticks, whereas the protein residues in gray and raspberry sticks) showing R162 conformation pointing outside the pocket to allow **AzoF** binding. This conformation of R162 has also been reported crystallographically (shown in teal sticks, PDB: 6WRT). AF2 predicts the two possible conformations of R162: model ranked 1 positions R162 in the space left by the Y32G mutation, whereas model ranked 2 places R162 outside the pocket similarly to the conformation adopted in 6WRT crystal structure.

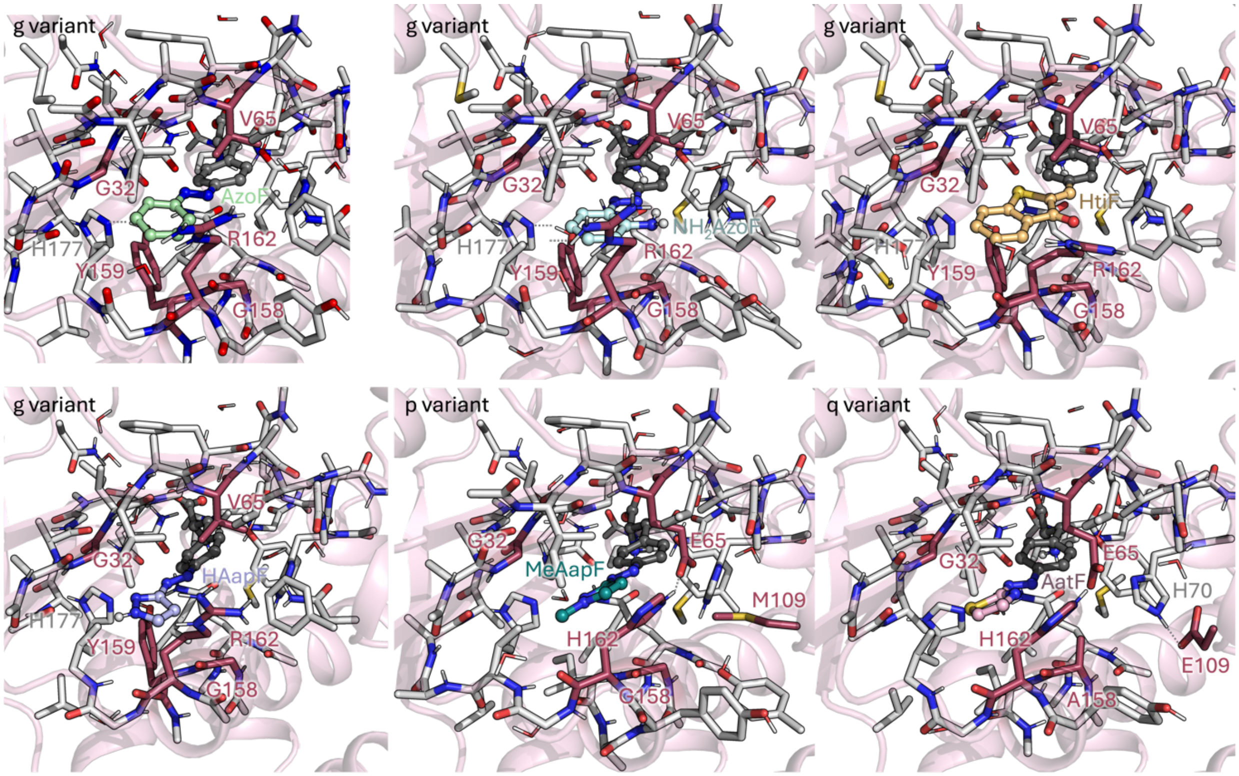

**Figure S17.** Optimized cluster models of aaRS-g, -p, and -q. The optimized geometry of aaRS-g containing **AzoF** (top left, shown in gray and green spheres), **$NH_2$AzoF** (top middle, in gray and cyan), **HtiF** (top right, in gray and gold), and **HAapF** (bottom left, in gray and light blue). The optimized structure of aaRS-p containing **MeAapF** (in gray and teal) bound in the pocket is shown in the bottom middle, and the aaRS-q containing **AatF** (in gray and pink) in the bottom right. All mutations are colored in raspberry and labeled.

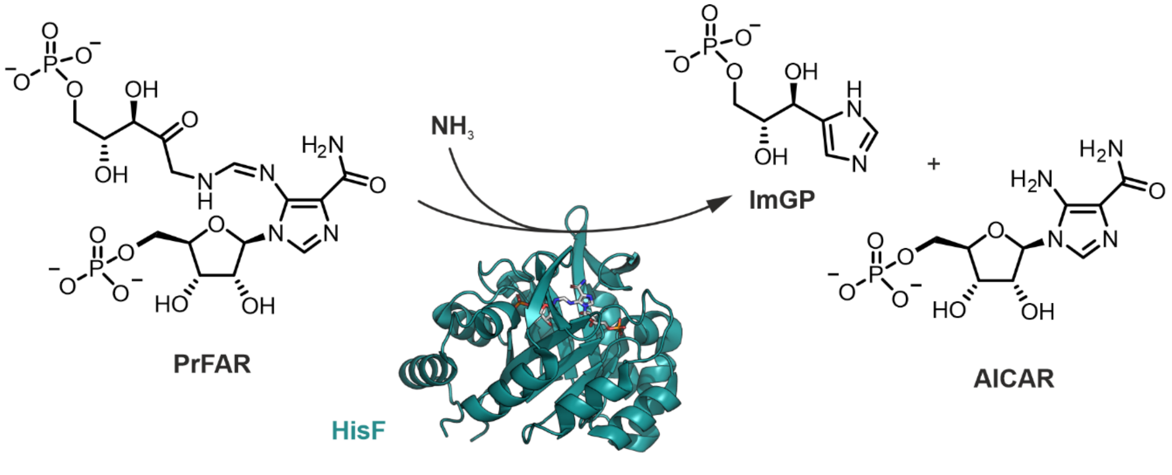

**Figure S18.** Reaction catalyzed by HisF (PDB: 7ac8). Turnover of *N'*-[(5'-phosphoribulosyl)formimino]-5-amino-imidazole-4-carboxamide ribonucleotide (PrFAR) and $NH_3$ to yield imidazole glycerol phosphate (ImGP) and 5-aminoimidazol-4-carboxamidribotide (AICAR).

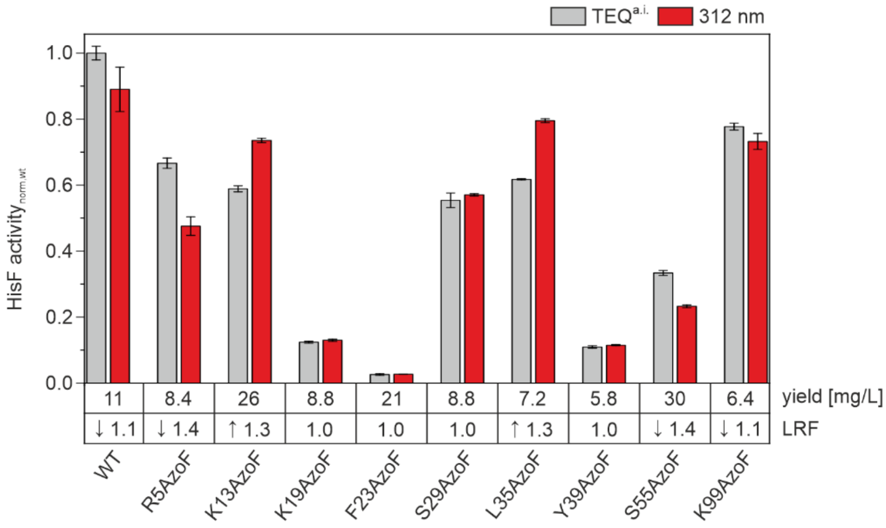

**Figure S19.** Screening of HisF-AzoF variants comparing their activities in the as-isolated, thermally equilibrated state (TEQ$^{a.i.}$, grey) and after irradiation with 312 nm in their PSS$^{312}$. Expression yields in mg per liter expression medium and the LRF values upon irradiation are given for each variant below the bar graph. Reactions were prepared in a 96-well plate at 25 °C and measured in a Tecan Infinite M200 Pro. Reaction conditions: 50 mM Tris acetate pH 8.5, 100 mM ammonium acetate, 1 μM HisA (to convert ProFAR to PrFAR), 70 μM ProFAR (saturation) and 0.2 μM HisF (reaction start). Activities were determined in μM min$^{-1}$ and normalized to the activity of wildtype HisF. WT: wildtype HisF.

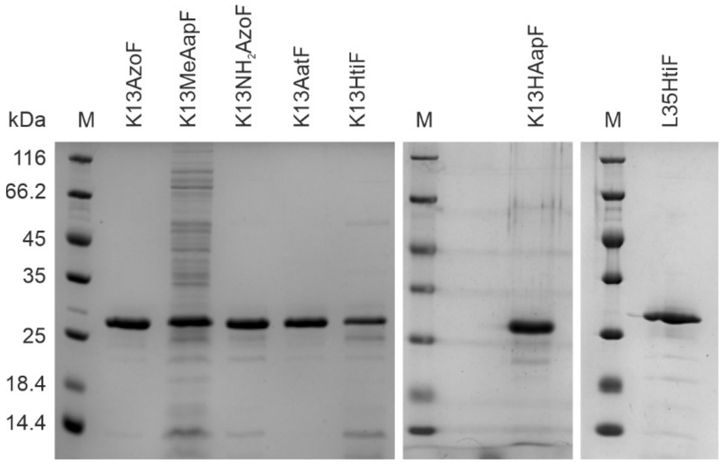

**Figure S20.** SDS-PAGE of the HisF-psUAA variants. 7 μL low-molecular weight marker (M) and 3 μg of each protein were applied to the gel. The variants showed purities of >95% (K13AzoF), >70% (K13MeAapF), >90% (K13$NH_2$AzoF), >95% (K13AatF), >50% (K13HtiF), >95% (K13HAapF), and >95% (L35HtiF, 13 mg/L expression culture).

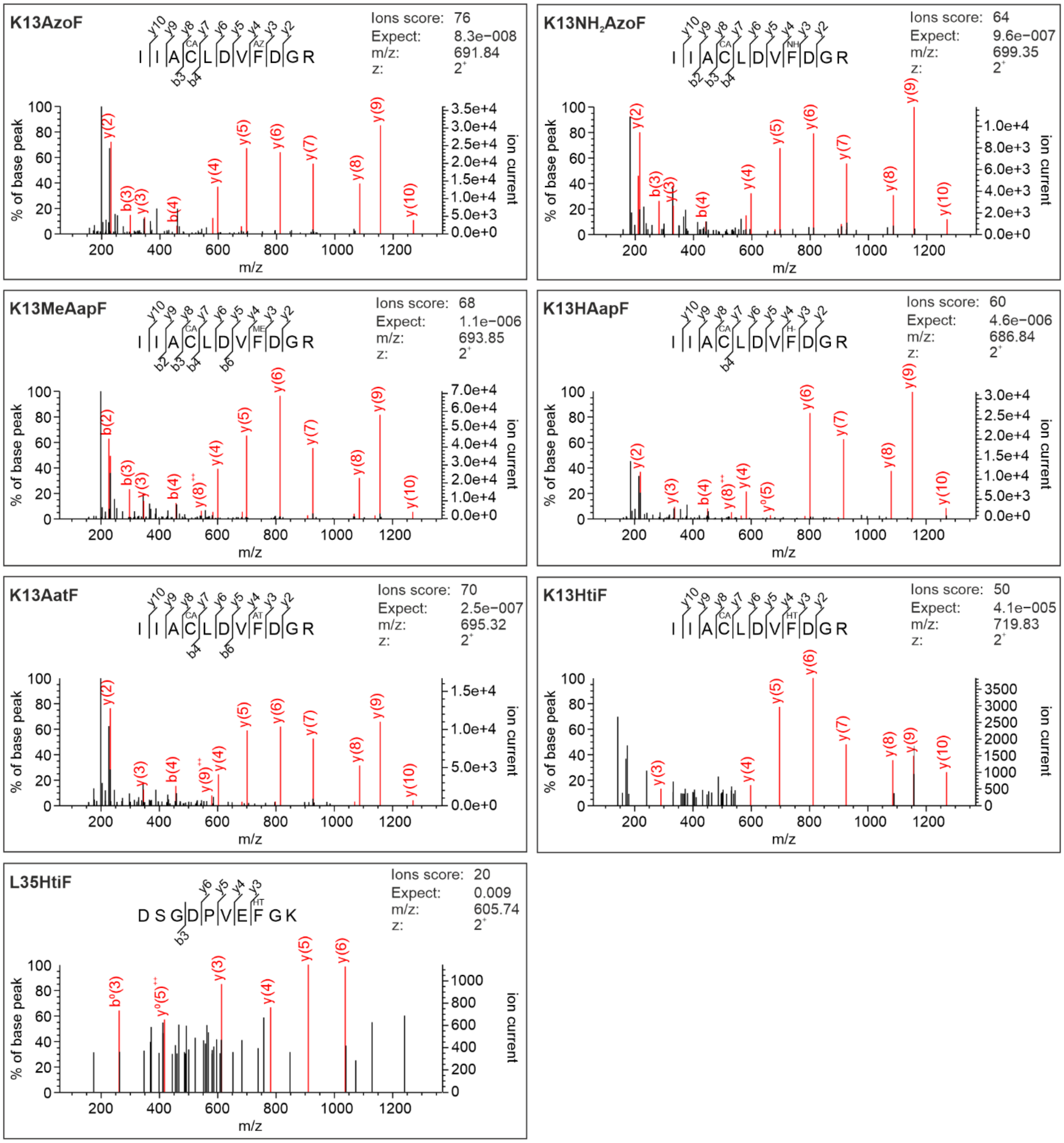

**Figure S21.** Confirmation of correct incorporation of psUAAs in HisF from *Thermotoga maritima* by LC-ESI-MS/MS. Shown are the fragment spectra of tryptic peptides containing **AzoF** (AZ), **NH$_2$AzoF** (NH), **MeAapF** (ME), **HAapF** (H-), **AatF** (AT), and **HtiF** (HT). CA: carbamidomethyl.

**Table S10.** Absorbance maxima $\lambda_{max}$ of each psUAA in 50 mM Tris/HCl pH 7.5 and DMSO.

| psUAA | isomer | $\lambda_{max}$[a] [nm] | $\lambda_{max}$[b] [nm] |
|---|---|---|---|
| **K13AzoF** | *E* | 333 | 425 |
| | *Z* | —[c] | 426 |
| **K13NH$_2$AzoF** | *E* | ~333 | ~420 |
| | *Z* | ~333 | ~420 |
| **K13MeAapF** | *E* | 333 | ~420 |
| | *Z* | —[c] | ~420 |
| **K13HAapF** | *E* | 331 | 419 |
| | *Z* | —[c] | 420 |
| **K13AatF** | *E* | 375 | —[d] |
| | *Z* | 326 | 463 |
| **K13HtiF** | *E* | ~335 | ~448 |
| | *Z* | ~335 | ~448 |

[a] $\pi \rightarrow \pi^*$ transition in case of diazo compounds. [b] $n \rightarrow \pi^*$ transition in case of diazo compounds. [c] cannot be determined due to overlap with protein signal [d] no spectroscopic signal.

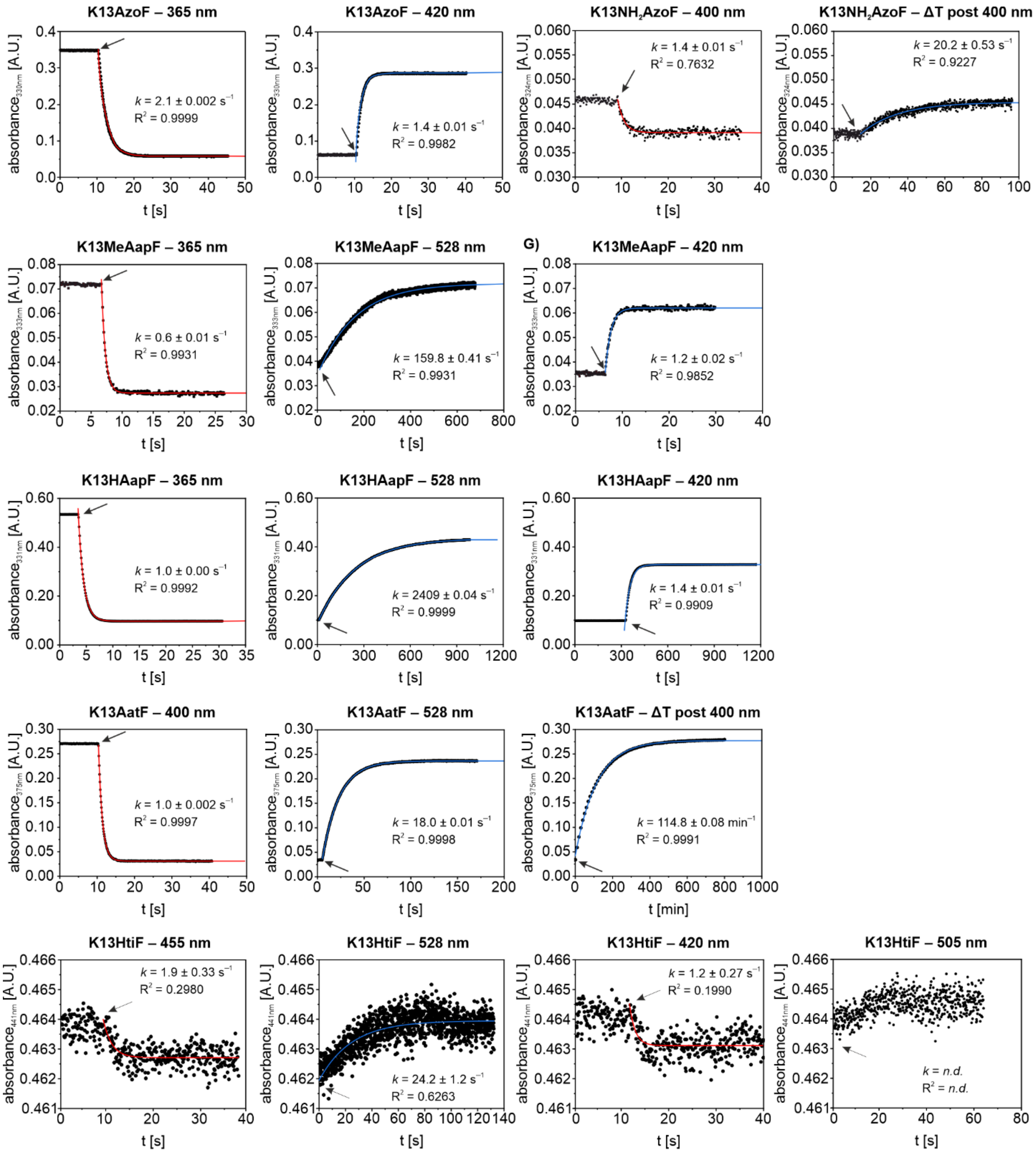

**Figure S22.** Exemplary isomerization rates of each HisF-psUAA in 50 mM Tris/HCl pH 7.5, 100 mM NaCl upon irradiation with the selected wavelengths or through thermal relaxation (*ΔT*). The absorbance was followed at the π→π* or n→π* transition. Irradiation (arrow) was started and either maintained until a plateau was reached (for evaluation of isomerization upon irradiation) or stopped (for evaluation of thermal relaxation). The rates of isomerization *k* were determined by fitting the data with a mono-exponential function (red). For K13HtiF irradiation with 505 nm, the difference in the absorbance signal was insufficient to determine *k*. Half-lives of isomerization $t_{½}$ were deduced from the respective *k* values and are listed in **Table S11**.

**Table S11.** Determination of isomerization half-lives ($t_{½}$) for K13psUAA variants in 50 mM Tris/HCl pH 7.5, 100 mM NaCl at 25 °C.

| Protein | $\lambda$ | selected as best wavelength in …[a] | $t_{½}$ [b] |
|---|---|---|---|
| K13AzoF | 365 nm | Tris/HCl, DMSO | 1.40 ± 0.001 s |
| | 420 nm | Tris/HCl, DMSO | 0.90 ± 0.01 s |
| K13NH$_2$AzoF | 400 nm | Tris/HCl, DMSO | 0.90 ± 0.07 s |
| | $\Delta T$ | | 14.0 ± 0.37 s |
| K13MeAapF | 365 nm | Tris/HCl, DMSO | 0.40 ± 0.01 s |
| | 420 nm | DMSO | 0.80 ± 0.01 s |
| | 528 nm | Tris/HCl | 111 ± 0.28 s |
| K13HAapF | 365 nm | Tris/HCl, DMSO | 0.70 ± 0.00 s |
| | 420 nm | DMSO | 0.95 ± 0.01 s |
| | 528 nm | Tris/HCl | 167.0 ± 0.03 s |
| K13AatF | 400 nm | Tris/HCl, DMSO | 0.70 ± 0.001 s |
| | 528 nm | Tris/HCl, DMSO | 12.5 ± 0.01 s |
| | $\Delta T$ | | 79.6 ± 0.06 min |
| K13HtiF | 455 nm | Tris/HCl | 1.29 ± 0.23 s |
| | 528 nm | Tris/HCl | 16.9 ± 0.83 s |
| | 420 nm | DMSO | 0.83 ± 0.19 |
| | 505 nm | DMSO | –[c] |

[a] cf. **Figure 3** and **Figure S2**. [b] Isomerization half-lives induced by irradiation or thermally were derived from the respective isomerization rates *k* (see **Figure S22**). [c] could not be determined because of insufficient absorbance signal.

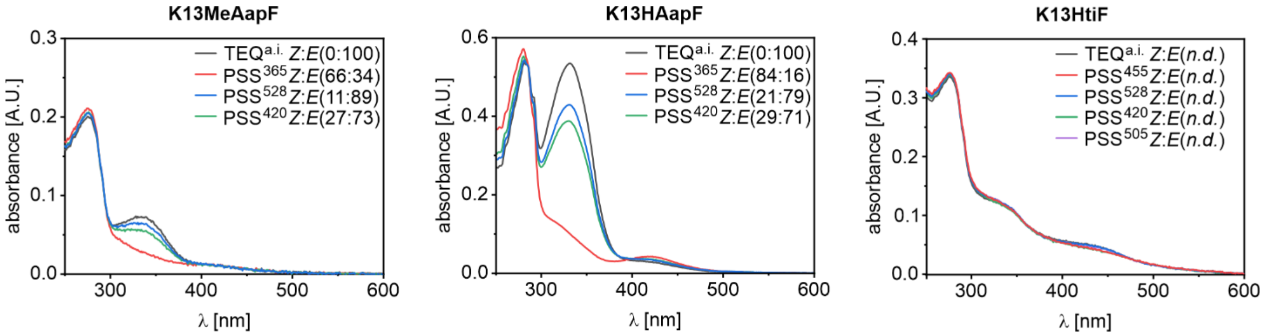

**Figure S23.** Comparison of the most effective wavelengths for the isomerization of K13MeAapF, K13HAapF, and K13HtiF in 50 mM Tris/HCl pH 7.5, 100 mM NaCl. The isolated psUAAs **MeAapF**, **HAapF**, and **HtiF** showed diverging effective wavelengths in buffer and DMSO (cf. **Figures S4–S5**) and thus, these wavelengths had to be re-evaluated in the protein context.

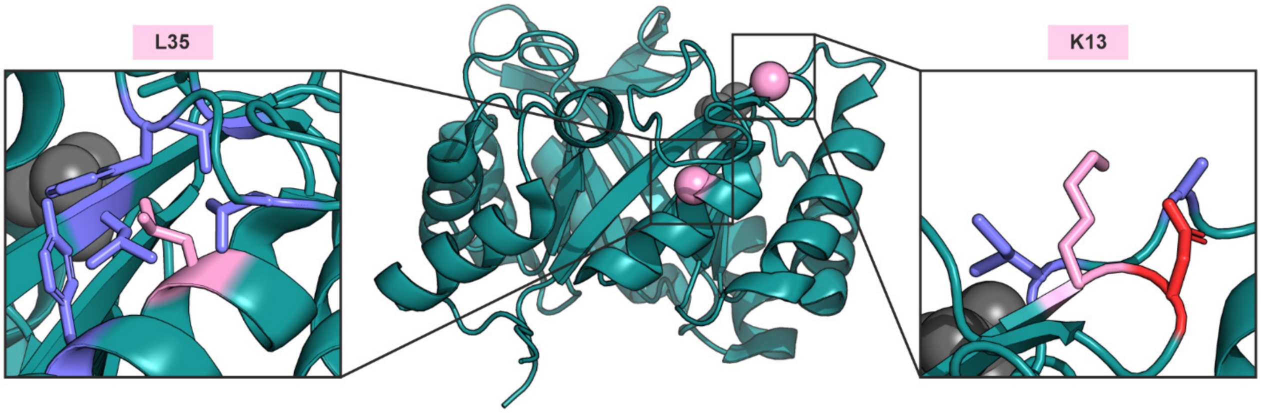

**Figure S24.**Localization of positions K13 and L35 in HisF (PDB: 7ac8). While position K13 points into the solvent and is close to a charged residue (red), L35 is part of a hydrophobic pocket (purple).

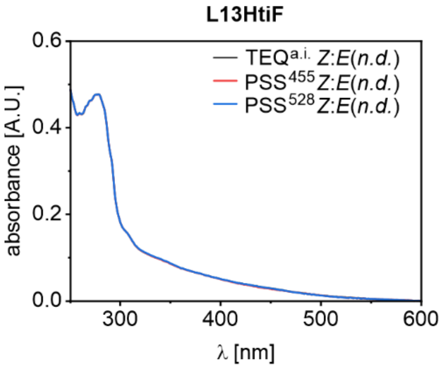

**Figure S25.**Absorbance spectra of 23 µM L35HtiF in 50 mM Tris/HCl pH 7.5, 100 mM NaCl. L35HtiF exhibits light scattering and a less pronounced absorbance signal, with no change upon irradiation.

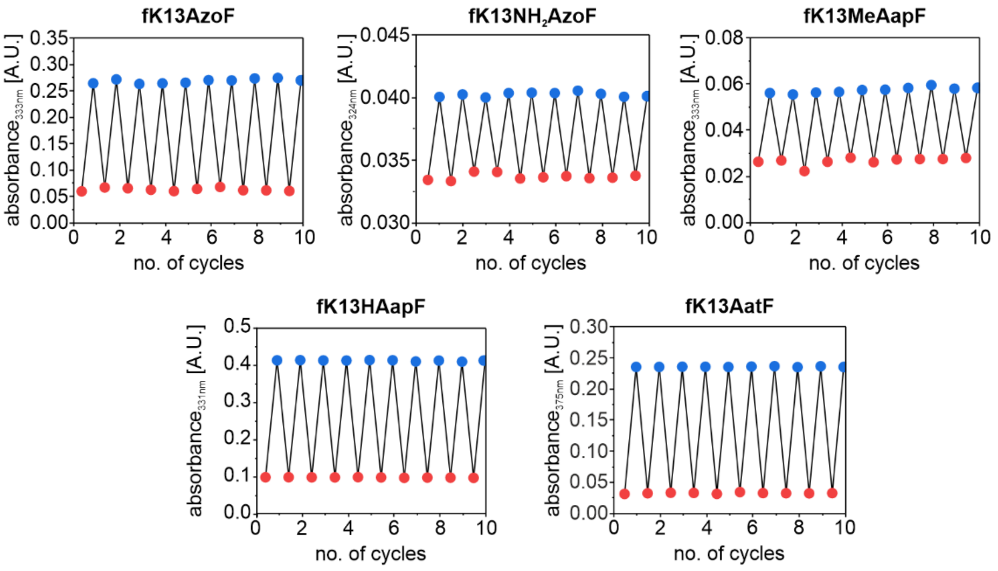

**Figure S26.** Cycle performance of the photoisomerization of HisF-psUAA variants in 50 mM Tris/HCl pH 7.5, 100 mM NaCl. Alternating isomerization was induced between $PSS^{365}$ and $PSS^{420}$ for K13AzoF, $PSS^{385}$ and $TEQ^{post}$ for $NH_2$AzoF, $PSS^{365}$ and $PSS^{420}$ for K13MeAapF, $PSS^{365}$ and $PSS^{505}$ for K13HAapF, and $PSS^{400}$ and $PSS^{528}$ for K13AatF. The absorbance at the characteristic maxima was plotted against the number of cycles (switching to *E* and subsequently to *Z* represents one cycle). All HisF-psUAA variants tolerated repeated photoswitching for ten cycles without photobleaching effects (decrease in absorbance) or light scattering effects (increase in absorbance). For K13MeAapF and K13HAapF, $PSS^{420}$ and $PSS^{505}$ instead of $PSS^{528}$ were used for the sake of time, because considerably less time was required for switching over ten cycles, while simultaneously yielding comparable photoconversion yields (cf. **Figure S23**). For **HtiF**, minor fluctuations in overall absorbance combined with insufficient absorption changes prevented the acquisition of reliable data.

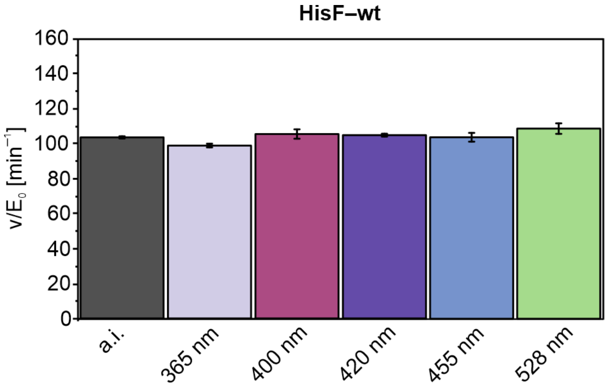

**Figure S27.** HisF-wt activity before and after irradiation with the most effective wavelengths for psUAA isomerization in the presence of 7 µM ProFAR (close to $K_m$ of HisF-wt). To this end, HisF-wt was subjected to the same irradiation conditions verifying that the enzyme system is not impaired by light exposure. Reaction conditions included 50 mM Tris acetate pH 8.5, 100 mM ammonium acetate, 1 µM HisA (to convert ProFAR to PrFAR) and 7 µM ProFAR at 25 °C. Measurements were performed in technical triplicates, which were fitted simultaneously to obtain a mean fit value (column) and the respective fit error (S.E.; error bars). The LRF values comparing $TEQ^{a.i.}$ with either $PSS^{\lambda}$ were obtained by selecting the respective data sets of triplicates and fitting them with a global fit analysis using a linear regression interaction model. Obtained $v/E_0$ ± S.E. and LRF ± S.E. values are listed in **Table S12**.

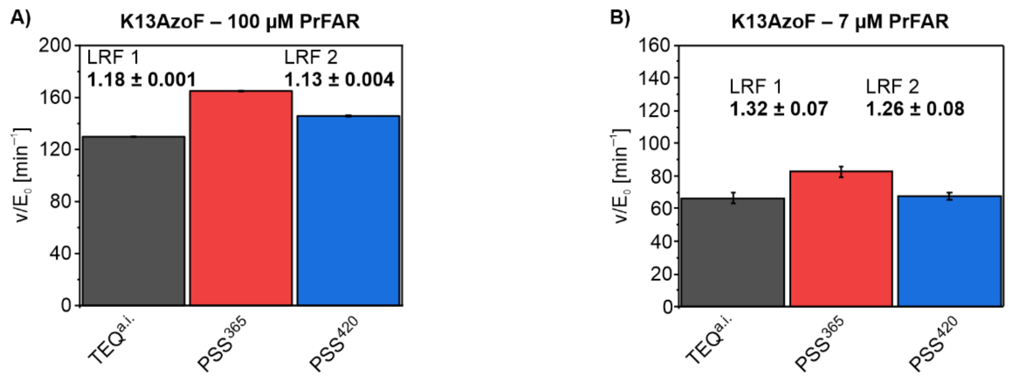

**Figure S28.** Photocontrol of K13AzoF activity in the presence of 100 µM PrFAR (saturation; A) or 7 µM PrFAR (close to $K_m$ of HisF-wt, B). Reaction conditions included 50 mM Tris acetate pH 8.5, 100 mM ammonium acetate, 1 µM HisA (to convert ProFAR to PrFAR) and 7/100 µM ProFAR at 25 °C. The reaction was started with 33 nM K13AzoF either in its $TEQ^{a.i.}$, $PSS^{365}$ or $PSS^{420}$. Measurements were performed in technical triplicates, which were fitted simultaneously to obtain a mean fit value (column) and the respective fit error (S.E.; error bars). The LRF values comparing either $TEQ^{a.i.}$ with $PSS^{365}$ or $PSS^{365}$ with $PSS^{420}$ were obtained by selecting the respective data sets of triplicates and fitting them with a global fit analysis using a linear regression interaction model.

**Table S12.** Photocontrol of HisF-wt and HisF-psUAA activity and deduced LRF values.

| Protein | | $v/E_0$ [$min^{-1}$] | LRF[c] | p-value of LRF[d] |
|---|---|---|---|---|
| HisF-wt[a] | TEQ$^{a.i.}$ | 102.8 ± 0.5 | | |
| | PSS$^{365}$ | 98.2 ± 1.1 | ↓ 1.05 ± 0.01[e] | 2.20E-294 |
| | PSS$^{400}$ | 104.7 ± 2.7 | ↑ 1.02 ± 0.03[e] | 5.93E-147 |
| | PSS$^{420}$ | 104.1 ± 0.8 | ↑ 1.01 ± 0.01[e] | 0 |
| | PSS$^{455}$ | 103.1 ± 2.5 | ↑ 1.00 ± 0.02[e] | 6.41E-155 |
| | PSS$^{528}$ | 108.0 ± 3.0 | ↑ 1.05 ± 0.03[e] | 4.13E-135 |
| K13AzoF (100 µM PrFAR) [b] | TEQ$^{a.i.}$ | 129.9 ± 0.06 | | |
| | PSS$^{365}$ | 165.0 ± 0.31 | ↑ 1.18 ± 0.001[f] | 0 |
| | PSS$^{420}$ | 145.8 ± 0.44 | ↓ 1.13 ± 0.004[f] | 0 |
| K13AzoF[a] | TEQ$^{a.i.}$ | 65.0 ± 2.7 | | |
| | PSS$^{365}$ | 85.5 ± 3.3 | ↑ 1.32 ± 0.07[f] | 6.80E-60 |
| | PSS$^{420}$ | 67.6 ± 2.4 | ↓ 1.26 ± 0.08[f] | 1.62E-52 |
| K13$NH_2$AzoF[a] | TEQ$^{a.i.}$ | 84.4 ± 0.5 | | |
| | PSS$^{400}$ | 100.6 ± 0.8 | ↑ 1.19 ± 0.01[f] | 0 |
| | TEQ$^{post}$ | 87.0 ± 0.8 | ↓ 1.15 ± 0.01[f] | 0 |
| K13MeAapF[a] | TEQ$^{a.i.}$ | 24.6 ± 0.3 | | |
| | PSS$^{365}$ | 33.5 ± 0.5 | ↑ 1.37 ± 0.02[f] | 0 |
| | PSS$^{528}$ | 24.4 ± 0.4 | ↓ 1.37 ± 0.03[f] | 0 |
| K13HAapF[a] | TEQ$^{a.i.}$ | 55.9 ± 0.6 | | |
| | PSS$^{365}$ | 82.5 ± 0.5 | ↑ 1.48 ± 0.03[f] | 0 |
| | PSS$^{528}$ | 59.9 ± 0.6 | ↓ 1.38 ± 0.03[f] | 0 |
| K13AatF[a] | TEQ$^{a.i.}$ | 56.3 ± 0.5 | | |
| | PSS$^{400}$ | 92.9 ± 0.6 | ↑ 1.64 ± 0.02[f] | 0 |
| | PSS$^{528}$ | 67.6 ± 0.8 | ↓ 1.37 ± 0.02[f] | 0 |
| K13HtiF[a] | TEQ$^{a.i.}$ | 23.1 ± 0.6 | | |
| | PSS$^{455}$ | 22.0 ± 0.8 | ↑ 1.05 ± 0.04[f] | 8.75E-95 |
| | PSS$^{528}$ | 22.4 ± 0.6 | ↓ 1.02 ± 0.05[f] | 5.43E-74 |

[a] Reactions were conducted with PrFAR close to $K_m$ as described in **Figure S28**. [b] Reactions were conducted with PrFAR in saturation. [c] generally delineates the activity ratio $\frac{v1}{v2}$ with $v_1 > v_2$ with arrows indicating an activity increase (↑) or decrease (↓). [d] p-values <0.05 indicate a significantly different activity in the two compared data sets. [e] LRF values compare TEQ$^{a.i.}$ with either PSS$^{\lambda}$. [f] LRF values compare either TEQ$^{a.i.}$ with PSS$^{365}$ / PSS$^{400}$ / PSS$^{455}$ or PSS$^{365}$ / PSS$^{400}$ / PSS$^{455}$ with PSS$^{420}$ / TEQ$^{post}$ / PSS$^{528}$.

**Table S13.** Primers used in this work.

| purpose | primers used[a] |
|---|---|
| generation of pGLNS from pEVOL (deletion of pBAD cassette) | GAGCTCCCGGTCATCAATCATCC |
| | CAACTTATATCGTATGGGGCTGAC |
| introduction of restriction sites in pGLNS for subcloning of aaRS | AAAAAA**GGTCTCA**CCATATGGGATTCCTCAAAGCG |
| | AAAAAA**GGTCTCT**GTAACTGCAGTTTCAAACGCTAAATTGC |
| introduction of restriction sites in pBAD for subcloning of sfGFP | AAAAAA**GAAGACAT**CCATAGATCTAATTCCTCCTGTTAGC |
| | AAAAAA**GAAGACAT**GTAATTAAACGGTCTCCAGCTTGGCTG |
| preparation of AzoF-RS$_{copt}$ for pGLNS | AAAAAA**GGTCTCA**AATGGATGAGTTCGAAATGATTAAAC |
| | AAAAAA**GGTCTCT**TTACAGACGTTTACGAATCGGTTCCAGG |
| introduction of Q155V in AzoF-RS$_{copt}$ | GATTATGGTGGTTAACGGCATTCATTATC |
| | GGATAAATCACTTCGGCAACTTTCG |
| introduction of A167V in AzoF-RS$_{copt}$ | GTGTTGATGTTGTGGTTGGTGGTATGG |
| | CATGATAATGAATGCCGTTAACCTG |
| introduction of A167C in AzoF-RS$_{copt}$ | GTGTTGATGTTTGCGTTGGTGGTATGG |
| | CATGATAATGAATGCCGTTAACCTG |
| introduction of V188P in AzoF-RS$_{copt}$ | CGAAGAAACCGGTTTGTATTCATAATCC |
| | GCAGCAGTTCACGTGCCAGC |
| introduction of G32A in AzoF-RS$_{copt}$ | CGCGATTGGTTTTGAACCGAGC |
| | GCTGATTTTTCATCCTTTTTCAGTACTTCG |
| introduction of E65L in AzoF-RS$_{copt}$ | CGATCTGCATGCATATCTGAATCAGAAAGG |
| | GCCAGCAGAATGATAATATCGAAACCG |
| introduction of A108F in AzoF-RS$_{copt}$ | GAATTTGAGCTGGATAAAGATTATACC |
| | GCTACCATAAACATATTTGGCTTTCAGGC |
| introduction of E109M in AzoF-RS$_{copt}$ | GAAGCCATGCTGGATAAAGATTATACC |
| | GCTACCATAAACATATTTGGCTTTCAGGC |
| introduction of G158A in AzoF-RS$_{copt}$ | GCGATTCATTATCATGGTGTTGATGTTGCAG |
| | GTTAACCTGCATAATCGGATAAATCACTTCGG |
| introduction of H162F in AzoF-RS$_{copt}$ | GGCATTCATTATTTTGGTGTTGATGTTGCAG |
| | GTTAACCTGCATAATCGGATAAATCACTTCGG |
| introduction of I159Y in AzoF-RS$_{copt}$ | GCTATCATTATCATGGTGTTGATGTTGC |
| | CGTTAACCTGCATAATCGGATAAATCACTTCG |
| introduction of I159W in AzoF-RS$_{copt}$ | GCTGGCATTATCATGGTGTTGATGTTGC |
| | CGTTAACCTGCATAATCGGATAAATCACTTCG |
| introduction of I159F in AzoF-RS$_{copt}$ | GCTTTCATTATCATGGTGTTGATGTTGC |
| | CGTTAACCTGCATAATCGGATAAATCACTTCG |
| introduction of I159K in AzoF-RS$_{copt}$ | GCAAACATTATCATGGTGTTGATGTTGC |
| | CGTTAACCTGCATAATCGGATAAATCACTTCG |
| introduction of R257G in ONBY-RS$_{i}$ | CATTAAAGGCCCAGAAAAATTTGGTGGTGATCTGAC |
| | GTCAGCGGGTATTCCAGAAAGTATTTGGC |
| generation of Azo-RS 4 from Azo-RS 1[12] | GTGTTGATGTTGCGGTTGGTGGTATGG |
| | CACGATAATGATAGCCGTTAACCTGC |
| introduction of restriction sites in pBAD for subcloning of sfGPF_Y151TAG | AAAAAA**GAAGACAT**CATCGATCTAATTCCTCCTGTTAGC |
| | AAAAAA**GAAGACAT**GTAATTAAACGGTCTCCAGCTTGGCTG |

| | |
|---|---|
| preparation of ONBY-RS$_i$ for subcloning in pGLNS | AAAAAA**GGTCTC**AATGGATGAGTTCGAGATGATTAAACG |
| | AAAAAA**GGTCTCT**TTACAGGCGTTTACGAATCGGTTCC |
| preparation of AzF-RS for subcloning in pGLNS | AAAAAA**GGTCTC**AATGGACGAGTTCGAAATGATTAAAC |
| | AAAAAA**GGTCTCT**TTACAGACGTTTGCGAATTGGTTCCAGAATC |
| preparation of pET24a for subcloning of pBAD_sfGFP_Y151TAG_rrnB | AAAAAA**GAAGACAT**AAGGAGGAACTATATCCGG |
| | AAAAAA**GAAGACAT**CGCAACGCAATTAATGTAAGTTAG |
| preparation of araBAD_sfGFP_Y151TAG_rrnB for subcloning in pET24a | AAAAAAA**GAAGACAT**TGCGCTTATTAATCAGATAAAATATTTC |
| | AAAAAAA**GAAGACAT**CCTTGCACACGGTCACACTG |
| introduction of TAG151Y in sfGFP_Y151TAG | GTCTATATCACCGCAGATAAACAGAAAAACG |
| | GTTGTGGCTGTTGAAATTATATTCCAGTTTGTGG |

[a] recognition sites for BsaI or BbsI are shown in bold and the resulting four base pair overhangs are highlighted in red. Point mutations are underlined.

### 2. Materials and methods

#### 2.1 Synthesis of photoswitchable UAAs

AzoF was synthesized according to a previously published protocol.[13] Detailed synthesis schemes of psUAAs and corresponding procedures are given below. Starting materials and reagents were purchased from commercial suppliers (Sigma Aldrich, Alfa Aesar, Acros, Fluka, TCI or VWR) and used without further purification. Solvents were used as p.a. grade or distilled. Dry solvents were dried over molecular sieves. For automated flash column chromatography industrial grade of solvents was used. All NMR spectra were measured at room temperature using a Bruker Avance 300 (300 MHz for $^{1}$H, 75 MHz for $^{13}$C, 282 MHz for $^{19}$F) or a Bruker Avance 400 (400 MHz for $^{1}$H, 101 MHz for $^{13}$C, 376 MHz for $^{19}$F) NMR spectrometer. All chemical shifts are reported in $\delta$-scale as parts per million [ppm] (multiplicity, coupling constant J, number of protons) relative to the solvent residual peaks as the internal standard. Coupling constants J are given in Hertz [Hz]. Abbreviations used for signal multiplicity: $^{1}$H NMR: b = broad, s = singlet, d = doublet, t = triplet, q = quartet, hept = heptet dd = doublet of doublets, dt = doublet of triplets, dq = doublet of quartets, and m = multiplet; $^{13}$C NMR: (+) = primary/tertiary, (-) = secondary, (Cq) = quaternary carbon. HRMS (high resolution mass spectra) and LRMS (low resolution mass spectra) were measured at the Central Analytical Laboratory of the University of Regensburg. These mass spectra were recorded on a Finnigan MAT 95, ThermoQuest Finnigan TSQ 7000, Finnigan MAT SSQ 710 A or an Agilent Q-TOF 6540 UHD instrument. Analytical TLC was performed on silica gel coated alumina plates (MN TLC sheets ALUGRAM® Xtra SIL G/UV254). Visualization was done by UV light (254 or 366 nm). If necessary, ninhydrin, vanillin or permanganate were used for chemical staining. Purification by column chromatography was performed with silica gel 60 M (40-63 µm, 230-440 mesh, Merck) or with a pre-packed Biotage® Sfär C18 D Duo 100 Å 30 µm column on a Biotage® IsoleraTM Spektra One device.

##### 2.1.1 Schemes

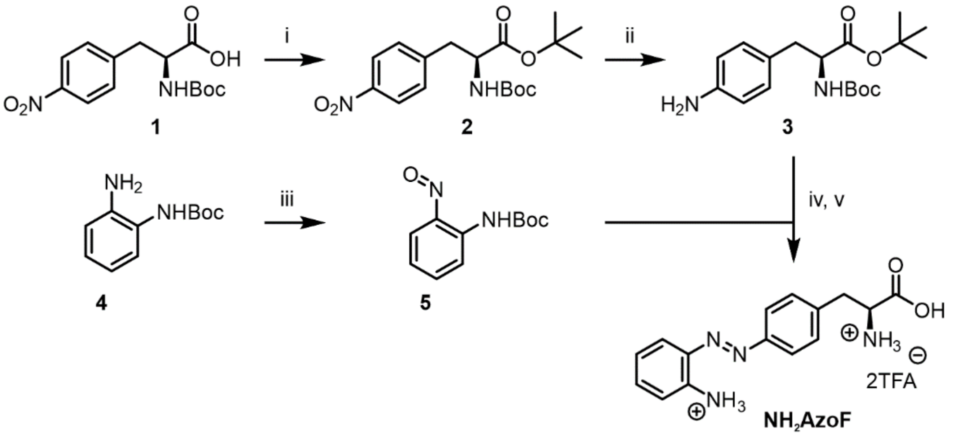

**Scheme S1:** Synthesis of **NH$_2$AzoF**. Reaction conditions: (i) Boc$_2$O, DMAP, tBuOH; (ii) H$_2$, Pd/C; (iii) oxone; iv) AcOH; v) TFA. Overall yield 10%.

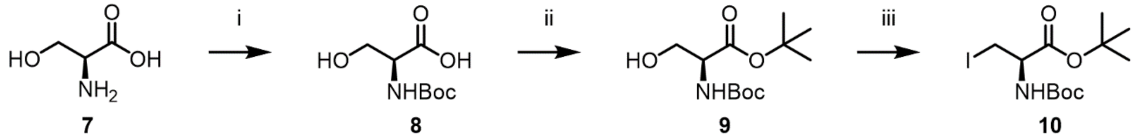

**Scheme S2:** Synthesis of *tert*-butyl (*R*)-2-((*tert*-butoxycarbonyl)amino)-3-iodopropanoate as precursor for the syntheses of **MeAapF**, **HtiF**, and **SpiroY**. Reaction condtions: (i) Boc$_2$O, NaOH; (ii) tert-butyl N,N'-diisopropylcarbamimidate; (iii) $I_2$, $PPh_3$, imidazole.

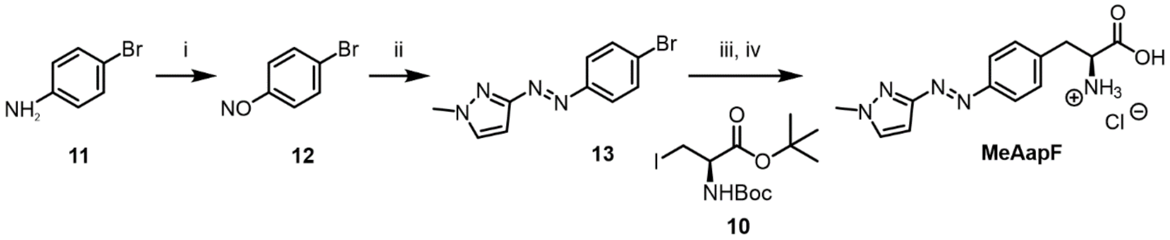

**Scheme S3:** Synthesis of **MeAapF**. Reaction conditions: (i) Oxone; (ii) 1-methyl-3-aminopyrazole, AcOH; (iii) Zn, 1,2.dibromoethane, TMSCl, Pd$_2$(dba)$_3$, SPhos; (iv) TFA. Synthesis of the precursor **10** is shown in Scheme 2. Overall yield 16%.

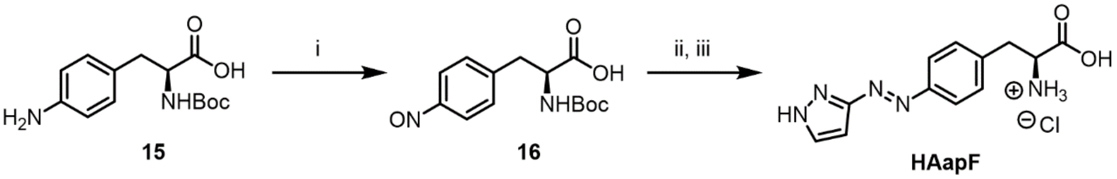

**Scheme S4:** Synthesis of **HAapF**. Reaction conditions: (i) Oxone; (ii) 3-aminopyrazole, AcOH; (iii) HCl. Overall yield 3%.

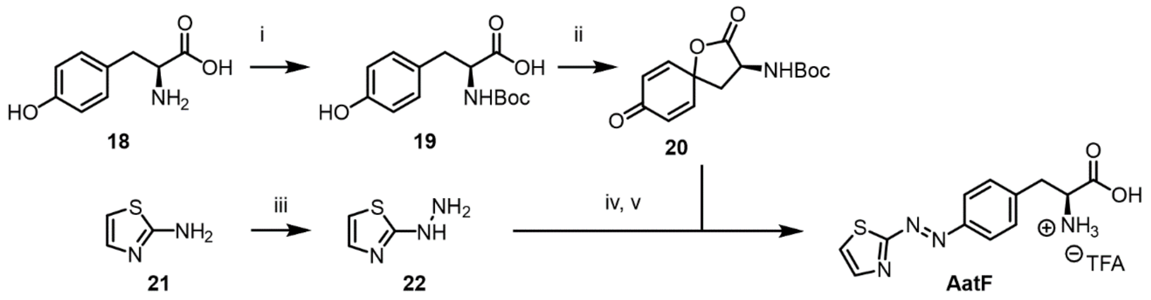

**Scheme S5:** Synthesis of **AatF**. Reaction conditions: (i) $Boc_2O$, $Et_3N$; (ii) $PhI(OAc)_2$; (iii) HCl, $NaNO_2$, $SnCl_2$; (iv) $Ce(NO_3)_6(NH_4)_2$; (v) TFA. Overall yield 6%.

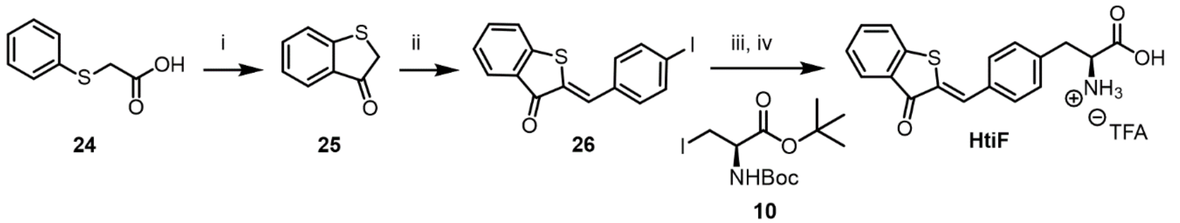

**Scheme S6:** Synthesis of **HtiF**. Reaction conditions: (i)TfOH; (ii) *p*-iodobenzaldehyde, piperidine; (iii) Zn, 1,2.dibromoethane, TMSCl, $Pd_2(dba)_3$, SPhos; (iv) TFA. Synthesis of the precursor **10** is shown in Scheme 2. Overall yield 4%.

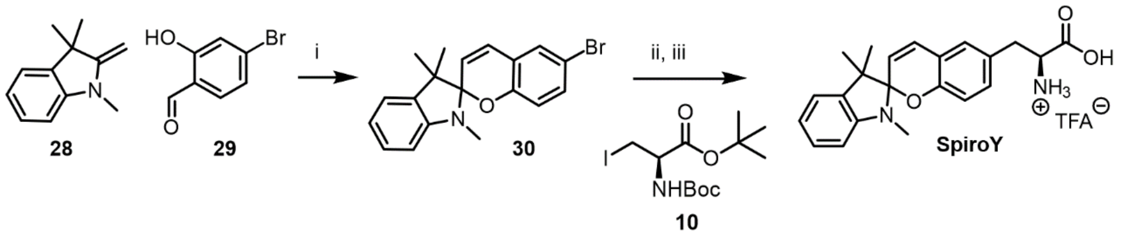

**Scheme S6:** Synthesis of **SpiroY**. Reaction conditions: (i) piperidine; (ii) Zn, 1,2.dibromoethane, TMSCl, $Pd_2(dba)_3$, SPhos; (iii) TFA. Synthesis of the precursor **10** is shown in Scheme 2. Overall yield 71%.

##### 2.1.2 Procedures

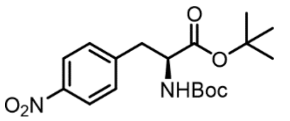

***tert*-butyl (*S*)-2-((*tert*-butoxycarbonyl)amino)-3-(4-nitrophenyl)propanoate 2:** Synthesized according to literature reports.[15] To Boc-4-nitrophenylalanine **1** (6.45 mmol, 1 eq) in 10 mL *t*-BuOH:DCM 1:2, DMAP (0.3 eq) Boc anhydride (1.1 eq) was added. Reaction mixture was stirred at room temperature (rt) overnight. All solvents were evaporated *in vacuo* and the product was purified by silica gel flash column chromatography (PE:EtOAc 9:1). Colorless oil turned to sticky white solid overnight, 87% yield. $^1$H-NMR (300 MHz, Chloroform-*d*): $\delta$ 8.16 (d, J = 8.7 Hz, 2H), 7.35 (d, J = 8.7 Hz, 2H), 5.07 (d, J = 7.7 Hz, 1H), 4.49 (q, J = 6.7 Hz, 1H), 3.36 - 2.93 (m, 2H), 1.41 (d, J = 1.2 Hz, 18H).

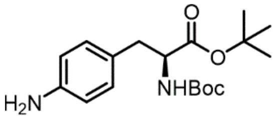

***tert*-butyl (*S*)-3-(4-aminophenyl)-2-((*tert*-butoxycarbonyl)amino)propanoate 3:** Compound **2** (1.4 mmol, 1 eq) was dissolved in 10 mL EtOH and Pd/C (0.05 eq) was added. After 3 vacuum/$H_2$ cycles, the reaction was left at room temperature overnight with a $H_2$ balloon. Filtered through Celite® and dried. Pinkish solid, 100% yield. $^1$H-NMR (300 MHz, Chloroform-*d*): $\delta$ 6.94 (d, J = 8.4 Hz, 2H), 6.61 (d, J = 8.4 Hz, 2H), 4.94 (d, J = 8.4 Hz, 1H), 4.48 - 4.29 (m, 1H), 3.61 (s, 2H), 2.94 (d, J = 5.9 Hz, 2H), 1.41 (d, J = 1.9 Hz, 18H).

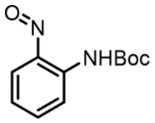

***tert*-butyl (2-nitrosophenyl)carbamate 5**: Synthesized according to literature reports.[16] To a solution of *tert*-butyl-N-(2-aminophenyl)carbamate **4** (2.4 mmol, 1 eq) in DCM (5 mL) a solution of Oxone® (2 eq) in water (20 mL) was added and stirred vigorously at ambient temperature for 1 h. The organic phase was extracted with DCM and the organic layers were combined, washed with 1 M HCl and saturated $NaHCO_3$, dried over $MgSO_4$, filtered and concentrated under reduced pressure. The resulting brown solid (43% yield) was directly used for the next step. $^1$H-NMR (300 MHz, Chloroform-*d*): $\delta$ 10.20 (s, 1H), 8.58 (dd, J = 8.6, 1.2 Hz, 1H), 7.65 (dddd, J = 8.8, 7.1, 1.7, 0.6 Hz, 1H), 7.49 - 7.30 (m, 1H), 7.06 (ddd, J = 8.2, 7.1, 1.2 Hz, 1H), 1.56 (s, 9H).

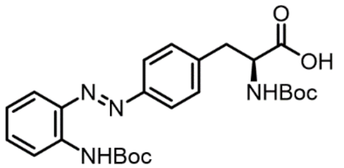

***tert*-butyl (*S*,*E*)-2-((*tert*-butoxycarbonyl)amino)-3-(4-((2-((*tert*-butoxy-carbonyl)amino)phenyl)diazenyl)phenyl)propanoate 6:** Compounds **3** (1.5 mmol, 1 eq) and **5** (1 eq) were mixed in 15 mL AcOH and stirred at rt overnight. The reaction was quenched with saturated $NaHCO_3$ and the organic phase was extracted with EtOAc, washed with brine and dried over $MgSO_4$. Separation by column chromatography (PE:EtOAc 95:5) obtained an orange oil in 25% yield. $^1$H-NMR (300 MHz, Chloroform-*d*): $\delta$ 9.21 (s, 1H), 8.38 (dd, J = 8.5, 1.4 Hz, 1H), 7.88 - 7.71 (m, 3H), 7.43 (ddd, J = 8.6, 7.3, 1.6 Hz, 1H), 7.39 - 7.31 (m, 2H), 7.08 (ddd, J = 8.2, 7.1, 1.2 Hz, 1H), 5.06 (d, J = 7.8 Hz, 1H), 4.51 (q, J = 6.2 Hz, 1H), 3.29 - 3.00 (m, 2H), 1.57 (s, 9H), 1.43 (d, J = 1.4 Hz, 18H).

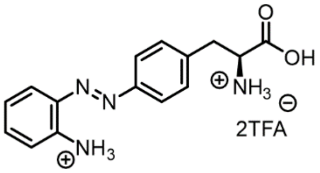

**(S,E)-2-amino-3-(4-((2-aminophenyl)diazenyl)phenyl)propanoic acid $NH_2$AzoF:** 0.37 mmol (1 eq) of **6** was dissolved in 5 mL DCM and excess TFA was added before the reaction was allowed to stir at rt for 2 days. The solvent was removed by rotatory evaporation and the residue was dissolved in a minimum amount of water and lyophilized. The compound was then triturated with $Et_2O$ and dried in vacuum to obtain the final product as a double TFA salt. Red solid in 97% yield. $^1$H-NMR (400 MHz, DMSO-*d$_6$*): $\delta$ 10.01 (s, 3H), 8.38 (s, 3H), 7.87 (d, J = 8.1 Hz, 2H), 7.64 (dd, J = 8.1, 1.6 Hz, 1H), 7.43 (d, J = 8.1 Hz, 2H), 7.28 - 7.09 (m, 1H), 6.88 (d, J = 8.3 Hz, 1H), 6.65 (t, J = 7.5 Hz, 1H), 4.26 (s, 1H), 3.19 (dd, J = 6.4, 2.7 Hz, 2H). $^{13}$C-NMR (101 MHz, DMSO-*d$_6$*): $\delta$ 170.4, 151.8, 145.1, 137.0, 135.9, 132.7, 130.5, 124.6, 122.3, 117.3, 115.8, 53.1, 35.7. HRMS (ESI-TOF) m/z calculated for $[C_{15}H_{17}N_4O_2]^+$ $[MH]^+$ 285.1347, found 285.1363.

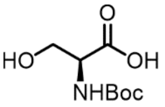

**(*tert*-butoxycarbonyl)-*L*-serine 8**: Synthesized according to a reported procedure.[1] A solution of *L*-serine **7** (47.5 mmol, 1 eq) in 1 M NaOH (50 mL) and dioxane (25 mL) at 0 °C was treated with $Boc_2O$ (1.2 eq) and the mixture was allowed to warm to rt and stirred for 24 h. The dioxane was then evaporated and the aqueous layer was washed with $Et_2O$. EtOAc was then added to the aqueous layer and the pH was adjusted with 1 M HCl to pH 2-3. The organic layer was extracted, and the aqueous layer was saturated with NaCl and extracted with EtOAc. The combined organic layers were dried and filtered, and the solvent was removed in vacuum to obtain a thick colorless oil in 100% yield. $^1$H-NMR (400 MHz, Methanol-*d$_4$*): $\delta$ 4.19 (t, J = 4.3 Hz, 1H), 3.83 (qd, J = 11.3, 4.3 Hz, 2H), 1.45 (s, 9H).

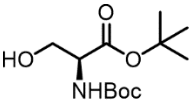

***tert*-butyl (*tert*-butoxycarbonyl)-*L*-serinate 9**: Synthesized according to a reported procedure.[2] **8** (4.9 mmol, 1 eq) was dissolved in DCM and cooled to 0 °C under $N_2$. *tert*-butyl N,N'-diisopropylcarbamimidate (3 eq) was added in three portions over 5 min. The reaction was stirred overnight at rt before it was stirred with hexane for 30 min, then filtered and concentrated under reduced pressure. The crude was purified by column chromatography (PE:EtOAc 3:1 to 1:1) to yield a colorless oil/sticky solid, with 57% yield. $^1$H-NMR (300 MHz, Chloroform-*d*): $\delta$ 5.42 (s, 1H), 4.25 (s, 1H), 3.89 (d, J = 3.8 Hz, 2H), 1.48 (s, 9H), 1.45 (s, 9H).

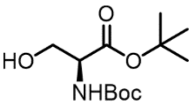

***tert*-butyl (*R*)-2-((*tert*-butoxycarbonyl)amino)-3-iodopropanoate 10**: Synthesized according to a reported procedure.[2] To a solution of triphenylphosphine (1.3 eq) and imidazole (1.3 eq) in DCM (20 mL) at 0°C was added iodine (1.3 eq) in 3 portions over 30 min. A solution of **9** (2.8 mmol, 1 eq) in DCM (20 mL) was added dropwise. The resultant solution was stirred at 0°C for 1 h before being warmed to room temperature for 2 h. Methanol (10 mL) was added, and the solvent was removed *in vacuo*. Purification by flash column chromatography (PE:EtOAc, 9:1). Colorless oil, 49% yield. $^1$H-NMR (300 MHz, Chloroform-*d*): $\delta$ 5.34 (d, J = 6.6 Hz, 1H), 4.34 (dt, J = 7.4, 3.5 Hz, 1H), 3.56 (d, J = 3.8 Hz, 2H), 1.50 (s, 9H), 1.45 (s, 9H).

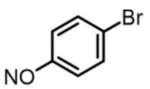
**1-bromo-4-nitrosobenzene 12**: Synthesized according to literature report.[10] A solution of Oxone® (2 eq) in water was added to a solution of 4-bromoaniline **11** (4.6 mmol, 1 eq) in DCM with vigorous stirring. The reaction mixture was stirred at room temperature for 3h. After disappearance of starting materials (indicated by TLC), the reaction was extracted with DCM. The combined organic layers were dried over $MgSO_4$ and concentrated under reduced pressure to give a light brown solid in quantitative yield. $^1$H-NMR (400 MHz, Chloroform-*d*): $\delta$ 7.78 (s, 4H) ppm.

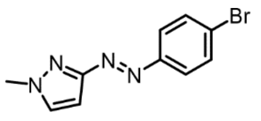
**(*E*)-3-((4-bromophenyl)diazenyl)-1-methyl-1*H*-pyrazole 13**: A solution of **12** and N-methyl-3-aminopyrazole in a mixture of DCM:AcOH 1:1 was stirred at rt overnight. The reaction was quenched with $NaHCO_3$, diluted with water and extracted with EtOAc, separated by column chromatography and dried in high vacuum. Yellow solid, 28% yield. $^1$H-NMR (400 MHz, Chloroform-*d*): $\delta$ 7.91 - 7.73 (m, 2H), 7.70 - 7.54 (m, 2H), 7.40 (d, J = 2.4 Hz, 1H), 6.64 (d, J = 2.5 Hz, 1H), 4.02 (s, 3H).

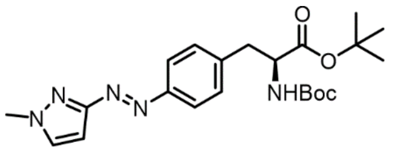
***tert*-butyl (*S*,*E*)-2-((*tert*-butoxycarbonyl)amino)-3-(4-((1-methyl-1*H*-pyrazol-3-yl)diazenyl)phenyl)propanoate 14**: Procedure adapted from literature.[11] Zinc (3 eq) was added to a dry crimp capped vial and purged with three cycles of vacuum/$N_2$. Dry DMF (1 mL), 1,2-dibromoethane (0.08 eq), and TMSCl (0.04 eq) were added to the flask. Stirred reaction was heated to 60°C under $N_2$ for 5 min, and then it was allowed to cool to rt before the addition of **10** (1.2 mmol, 1.5 eq). The sealed reaction was again purged with three cycles of vacuum/$N_2$ and sonicated without stirring for 30 min at 35°C. **13** (0.80 mmol, 1 eq), $Pd_2dba_3$ (0.025 eq.) and SPhos (0.05 eq.) were added in another crimp capped vial and purged with three cycles of vacuum/$N_2$ before 1 mL DMF was added. The organozinc mixture was carefully syringed, with minimal transfer of unreacted zinc, in the second vial and stirred overnight at 70°C under $N_2$. Reaction mixture was quenched with saturated $NH_4Cl$ and extracted with EtOAc. Organic phase was washed with brine, dried over $MgSO_4$, filtered, concentrated and dried under high vacuum. Crude was purified by flash chromatography (PE:EtOAc 95:5 to 90:10) to obtain a yellow oil in 63% yield. $^1$H-NMR (400 MHz, Chloroform-*d*): $\delta$ 7.88 (d, J = 8.5 Hz, 2H), 7.39 (d, J = 2.5 Hz, 1H), 7.31 (d, J = 8.5 Hz, 2H), 6.64 (d, J = 2.5 Hz, 1H), 5.05 (d, J = 7.5 Hz, 1H), 4.49 (d, J = 6.8 Hz, 1H), 4.03 (s, 3H), 3.13 (d, J = 5.7 Hz, 2H), 1.43 (s, 9H), 1.39 (s, 9H).

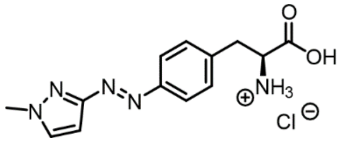
**(*S*,*E*)-2-amino-3-(4-((1-methyl-1*H*-pyrazol-3-yl)diazenyl)phenyl) propanoic acid MeAapF:** Compound **14** (0.57 mmol, 1 eq) was dissolved in dioxane:HCl 4M 1:1 and stirred for 2 days at rt. The solvent was evaporated, and the residue was lyophilized, then triturated with $Et_2O$ and dried in *vacuo* to obtain the final product as a HCl salt. Yellow solid, 91% yield. $^1$H-NMR (400 MHz, DMSO-$d_6$): $\delta$ 8.53 (dd, J = 4.8, 1.6 Hz, 3H), 7.84 (d, J = 2.4 Hz, 1H), 7.82 - 7.75 (m, 2H), 7.56 - 7.44 (m, 2H), 6.53 (d, J = 2.5 Hz, 1H), 4.23 (d, J = 5.5 Hz, 1H),

3.97 (s, 3H), 3.24 (d, J = 6.4 Hz, 2H). $^{13}$C-NMR (101 MHz, DMSO-$d_6$): $\delta$ 170.3, 163.0, 151.4, 138.5, 133.2, 130.7, 122.4, 94.5, 53.0, 35.5. HRMS (ESI-TOF) m/z calculated for $[C_{13}H_{16}N_5O_2]^+$ $[MH]^+$ 274.1299, found 274.1302.

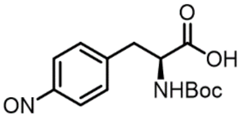

**(*S*)-2-((tert-butoxycarbonyl)amino)-3-(4-nitrosophenyl)propanoic acid 16:** To a solution of amino acid (7.13 mmol, 1 eq.) in DCM (15.0 mL/mmol) at 0 °C, Oxone® (7.13 mmol, 1.0 eq) in water (15.0 mL/mmol) was added. The orange, biphasic was allowed to stir for 1 h at 0°C, before the ice bath was removed and the mixture was stirred for another 1 h. Afterwards, the aqueous phase was decanted. To the remaining organic layer in the round bottom flask, water (2x 20 mL) was added and once again decanted off. The resulting organic DCM mixture in the round bottom flask was used directly in the next step. The water layer was further extracted with 1% AcOH (10 ml) and DCM (2x 10 ml). Without further drying, the organic phase was used directly in the next step.

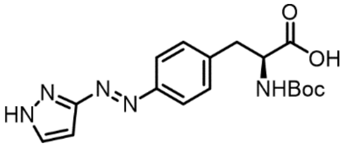

**(*S*,*E*)-3-(4-((1*H*-pyrazol-3-yl)diazenyl)phenyl)-2-((*tert*-butoxycarbonyl) amino)propanoic acid 17:** Following nitroso formation, 3-aminopyrazole (7.48 mmol, 1.10 eq.) and DCM/ glacial AcOH (15 mL/mmol) were added to the nitroso intermediate **16**. The orange solution was allowed to stir for 15 hours under $N_2$ atmosphere in the dark. The solvent was removed *in vacuo* and the crude product was subjected to reverse phase chromatography (solvent A: $H_2O$ + 0.05% TFA, solvent B: MeCN, gradient: 0–100% B) The product was obtained as a yellow solid (4%). $^1$H NMR (400 MHz, MeOD) δ 7.72 (d, J = 8.2 Hz, 2H), 7.59 (t, J = 4.9 Hz, 1H), 7.31 (d, J = 8.2 Hz, 2H), 6.54 (d, J = 2.5 Hz, 1H), 4.31 (dd, J = 9.1, 5.0 Hz, 1H), 3.23-3.13 (m, 3.2 Hz, 2H), 2.92-2.86 (m, 1H), 1.25 (s, 9H).

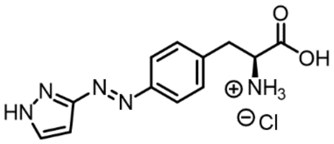

**(*S*,*E*)-3-(4-((1*H*-pyrazol-3-yl)diazenyl)phenyl)-2-aminopropanoic acid HAapF:** The Boc protected amine **17** (0.139 mmol, 1.0 eq.) was dissolved in dioxane (4.0 mL/mmol) at 0 °C and 4 N HCl (1.33 mL/mmol) was added dropwise. The reaction mixture was allowed to stir at room temperature for 5 h. Toluene (2.00 mL/mmol) was added, and the solvents were removed in *vacuo*. Subsequently, the product was triturated with diethyl ether (3x 1.0 mL/mmol) to obtain the target compound as a yellow solid (83%). $^1$H-NMR (400 MHz, MeOD): δ (ppm) = 7.91 (d, J = 8.40 Hz, 2H), 7.73 (d, J = 2.50 Hz, 1H), 7.49 (d, J = 8.40 Hz, 2H), 6.67 (d, J = 2.52 Hz, 1H), 4.30 (dd, J = 7.7, 5.6 Hz, 1H), 3.40 (dd, J = 14.5, 5.6 Hz, 1H), 3.25 (dd, J = 14.5, 7.7 Hz, 1H). $^{13}$C-NMR (101 MHz, MeOD): δ = 171.27, 164.71, 153.57, 139.38, 132.47, 131.48, 124.31, 95.41, 55.06, 37.20. ESI-MS (m/z): $[M^+H]^+$ calculated: 260.1142; found: 260.1143.

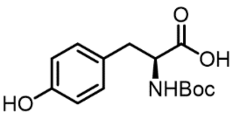

**(*tert*-butoxycarbonyl)-*L*-tyrosine 19**: Synthesized according to literature report.[12] To a solution of *L*-tyrosine (11 mmol, 1 eq) in 12 mL 1:1 dioxane/water was added $Et_3N$ (1.5 eq). The reaction flask was cooled to 0°C with an ice bath and Boc anhydride (1.1 eq) was added. After 1 h, the cold bath was removed, and the reaction mixture was stirred at ambient temperature for 20 h. The reaction mixture was then concentrated on a rotary evaporator and the residue diluted with water and ethyl acetate. The aqueous layer was acidified to pH 1 with 1 M HCl and extracted with ethyl acetate. The organic layer was washed with brine, dried over $MgSO_4$ and evaporated and a white solid was obtained with 74% yield. $^1$H-NMR (300 MHz, Chloroform-*d*): $\delta$ 7.01 (d, J = 8.0 Hz, 2H), 6.74 (d, J = 8.1 Hz, 2H), 4.99 (d, J = 7.8 Hz, 1H), 4.55 (d, J = 7.4 Hz, 1H), 3.05 (s, 2H), 1.43 (s, 9H).

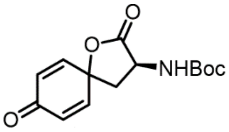

***tert*-butyl (*S*)-(2,8-dioxo-1-oxaspiro[4.5]deca-6,9-dien-3-yl)carbamate 20**: Synthesized according to literature report.[13] Phenyliodonium diacetate (1.1 eq) was added to a round-bottom flask containing 10 mL acetonitrile, and the mixture was stirred at room temperature. A solution of **19** (5 mmol, 1 eq) in 10 mL acetonitrile was added to the above round-bottom flask dropwise over 20 min. The mixture was allowed to stir for overnight before diluting with 20 mL ethyl acetate and 20 mL brine. The reaction mixture was extracted with ethyl acetate, and the organic layers were combined, dried over $MgSO_4$ and concentrated under reduced pressure. The crude mixture was purified by column PE:EtOAc 3:1 to 1:1 and a yellowish solid was obtained with 20% yield. $^1$H-NMR (300 MHz, Chloroform-*d*): $\delta$ 6.96 - 6.75 (m, 2H), 6.30 (td, J = 10.4, 2.1 Hz, 2H), 5.15 (s, 1H), 4.53 (s, 1H), 2.76 (dd, J = 12.9, 9.1 Hz, 1H), 2.53 - 2.34 (m, 1H), 1.46 (s, 9H).

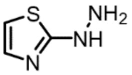

**2-hydrazineylthiazole 22**: A solution of sodium nitrite (1 eq) in water was added dropwise to a suspension of 2-aminothiazole (20 mmol 1 eq) in HCl (16mL) at –10°C. The cold diazonium salt solution was added dropwise to a solution of stannous chloride (2 eq) in HCl (20mL) at –10°C. The reaction mixture was stirred for 1 h, neutralized with NaOH 4 M, and filtered. The filtrate was brought to pH>9 extracted with EtOAc, dried over $MgSO_4$ and dried in vacuum. Grey solid was obtained with 11% yield. $^1$H-NMR (400 MHz, DMSO-$d_6$): $\delta$ 8.34 (s, 1H), 7.02 (d, J = 3.6 Hz, 1H), 6.63 (d, J = 3.6 Hz, 1H), 4.76 (s, 2H).

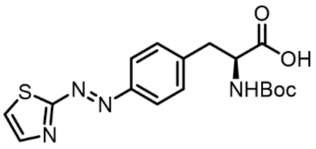

**(*S*,*E*)-2-((*tert*-butoxycarbonyl)amino)-3-(4-(thiazol-2-yldiazenyl)phenyl) propanoic acid 23**: Adapted from reported procedure.[14] **20** (1.1 mmol, 1 eq) and **22** (0.9 eq) were mixed in 10 mL acetonitrile in the presence of 3 mol% cerium(IV) ammonium nitrate at room temperature overnight. The solution was concentrated under reduced pressure. The crude product was purified by flash chromatography DCM:MeOH 100:0 to 90:10. Part of nonreacted spirolactone was also recovered. Orange solid was obtained with 55% yield. $^1$H-NMR (400 MHz, Chloroform-*d*): $\delta$ 8.02 (d, J = 3.3 Hz, 1H), 7.92 (d, J = 8.3 Hz, 2H), 7.41

(d, J = 3.4 Hz, 1H), 7.38 (d, J = 8.4 Hz, 2H), 5.21 (d, J = 8.1 Hz, 1H), 4.76 - 4.53 (m, 1H), 3.38 - 3.09 (m, 2H), 1.42 (s, 9H).

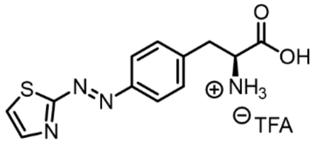

**(*S*,*E*)-2-amino-3-(4-(thiazol-2-yldiazenyl)phenyl)propanoic acid AatF:** Compound **23** (0.23 mmol, 1 eq) was dissolved in 5 mL DCM and excess TFA is added before the solution was stirred at rt for 2 days. The solvent was removed by rotatory evaporation and the residue was dissolved in a minimum amount of water and lyophilized. The compound was then triturated with $Et_2O$ and dried in vacuum to obtain the final product as a TFA salt. A red solid was obtained with 88% yield. $^1$H-NMR (400 MHz, DMSO-*d$_6$*): δ 8.15 (d, J = 3.3 Hz, 1H), 7.96 (d, J = 3.3 Hz, 1H), 7.93 (d, J = 8.4 Hz, 2H), 7.54 (d, J = 8.5 Hz, 2H), 4.16 (t, J = 6.6 Hz, 1H), 3.21 (qd, J = 14.4, 6.7 Hz, 2H). $^{13}$C-NMR (101 MHz, DMSO-*d$_6$*): δ 176.1, 170.2, 150.2, 144.4, 141.5, 131.0, 123.5, 123.4, 53.3, 36.0. HRMS (ESI-TOF) m/z calculated for $[C_{12}H_{13}N_4O_2S]^+$ $[MH]^+$ 277.0754, found 277.0760.

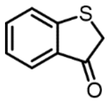

**benzo[*b*]thiophen-3(2*H*)-one 25**: Synthesized according to reported procedure.[9] To a flame-dried Schlenk flask equipped with a magnetic stir bar thiophenoxy acetic acid (3 mmol, 1 eq) and 10 mL DCM were added. To this solution, TfOH (5eq) was carefully added. Subsequently, the flask was sealed, placed into a prewarmed oil bath set to 40°C and stirred for 7 hours. After 7 hours, the reaction mixture was cooled to room temperature and carefully poured into ice/water. The organic layer was separated, and the aqueous layer was further extracted with DCM. The combined organic layers were washed with saturated $NaHCO_3$ and brine, before they were dried over anhydrous $MgSO_4$, filtered and concentrated under reduced pressure by rotary evaporation. The the crude product was obtained as a yellow solid in 52% yield and was immediately used without further purification. $^1$H-NMR (300 MHz, Chloroform-*d*): δ 7.78 (dd, J = 7.8, 1.2 Hz, 1H), 7.55 (ddd, J = 8.2, 7.2, 1.3 Hz, 1H), 7.43 (d, J = 8.0 Hz, 1H), 7.25 - 7.17 (m, 1H), 3.79 (s, 2H).

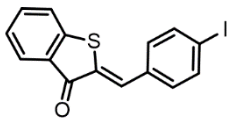

**(*Z*)-2-(4-iodobenzylidene)benzo[*b*]thiophen-3(2*H*)-one 26**: To a flame-dried 5 mL microwave vial 2**5** (1.25 eq), *p*-iodobenzaldehyde (1.1 mmol, 1 eq) and 1mL DMF were added. Then, piperidine (0.2 eq) was added to the solution. The reaction was stirred for five hours and then diluted with chloroform (20 mL) and washed with saturated aqueous $NH_4Cl$ (50 mL) and $H_2O$ (100 mL). The aqueous layer was then extracted with chloroform. The combined organic layers were then washed with brine (60 mL) and dried over anhydrous $Na_2SO_4$, filtered and concentrated. Purification by flash column chromatography. A yellow solid was obtained in 33% yield. $^1$H-NMR (300 MHz, Chloroform-*d*): δ 7.95 (ddd, J = 7.8, 1.4, 0.7 Hz, 1H), 7.89 - 7.78 (m, 3H), 7.60 (ddd, J = 8.5, 7.1, 1.4 Hz, 1H), 7.51 (dt, J = 8.0, 1.0 Hz, 1H), 7.47 - 7.37 (m, 2H), 7.32 (ddd, J = 8.1, 7.1, 1.0 Hz, 1H).

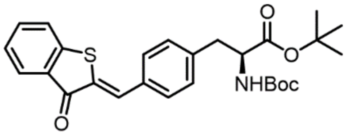

***tert*-butyl (*S,Z*)-2-((*tert*-butoxycarbonyl)amino)-3-(4-((3-oxobenzo[*b*]-thiophen-2(3*H*)-ylidene)methyl)phenyl)propanoate 27**: Same procedure as **14** but with **26** (0.33 mmol, 1 eq) instead of **13**. Yellow oil was obtained in 76% yield. $^{1}$H-NMR (400 MHz, Chloroform-*d*): *δ* 7.97 - 7.90 (m, 2H), 7.68 - 7.61 (m, 2H), 7.58 (ddd, J = 8.3, 7.1, 1.4 Hz, 1H), 7.53 - 7.48 (m, 1H), 7.34 - 7.27 (m, 3H), 5.05 (d, J = 8.1 Hz, 1H), 4.55 - 4.42 (m, 1H), 3.12 (h, J = 7.9, 7.2 Hz, 2H), 1.42 (d, J = 6.4 Hz, 18H).

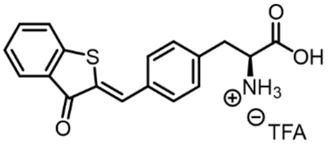

**(*S*,*Z*)-2-amino-3-(4-((3-oxobenzo[*b*]thiophen-2(3*H*)-ylidene)methyl) phenyl)propanoic acid HtiF**: **27** (0.25 mmol, 1 eq) was dissolved in 5 mL DCM and excess TFA was added before the solution was stirred at rt for 2 days. The solvent was removed *in vacuo* and the residue was dissolved in a minimum amount of water, followed by lyophilization. The compound was then triturated with $Et_2O$ and dried *in vacuo* to obtain the final product as a TFA salt. Orange solid in 73% yield. $^{1}$H-NMR (400 MHz, DMSO-$d_6$): *δ* 7.95 (s, 1H), 7.92 - 7.86 (m, 1H), 7.83 - 7.79 (m, 1H), 7.79 - 7.72 (m, 3H), 7.50 - 7.45 (m, 2H), 7.43 (td, J = 7.2, 1.2 Hz, 1H), 4.10 (t, J = 6.6 Hz, 1H), 3.23 - 3.07 (m, 2H). $^{13}$C-NMR (101 MHz, DMSO-$d_6$): *δ* 136.1, 131.1, 130.5, 129.6, 129.5, 126.6, 124.6. HRMS (ESI-TOF) m/z calculated for $[C_{18}H_{16}NO_3S]^+$ $[MH]^+$ 326.0846, found 326.0847.

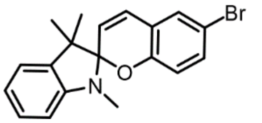

**6-bromo-1',3',3'-trimethylspiro[chromene-2,2'-indoline] 30**: To a solution of 1,3,3-trimethyl-2-methyleneindoline (1 eq) in EtOH, was added 5-bromosalicylaldehyde. The reaction mixture was refluxed overnight before being concentrated *in vacuo*. The crude product was purified by flash column chromatography to obtain a red oil in quantitative yield. $^{1}$H-NMR (300 MHz, Chloroform-*d*): *δ* 7.22 - 7.13 (m, 3H), 7.07 (ddd, J = 7.3, 1.4, 0.5 Hz, 1H), 6.85 (td, J = 7.4, 1.0 Hz, 1H), 6.78 (dd, J = 10.3, 0.8 Hz, 1H), 6.60 (dd, J = 9.1, 0.8 Hz, 1H), 6.56 - 6.48 (m, 1H), 5.73 (d, J = 10.3 Hz, 1H), 2.72 (s, 3H), 1.29 (s, 3H), 1.16 (s, 3H).

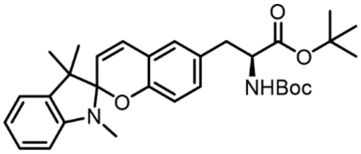

***tert*-butyl (2*S*)-2-((*tert*-butoxycarbonyl)amino)-3-(1',3',3'-trimethylspiro[chromene-2,2'-indolin]-6-yl)propanoate 31**: Same procedure as **14** but with **30** (1.3 mmol, 1 eq) instead of **13**. Red oil was obtaned in 71% yield. $^{1}$H-NMR (400 MHz, Chloroform-*d*): *δ* 7.18 (td, J = 7.5, 1.2 Hz, 1H), 7.07 (dd, J = 7.2, 1.4 Hz, 1H), 6.93 - 6.82 (m, 3H), 6.80 (d, J = 10.4 Hz, 1H), 6.63 (d, J = 8.0 Hz, 1H), 6.52 (d, J = 7.0 Hz, 1H), 5.67 (d, J = 10.2 Hz, 1H), 5.01 (d, J = 8.1 Hz, 1H), 4.39 (q, J = 6.5 Hz, 1H), 2.94 (d, J = 6.2 Hz, 2H), 2.72 (d, J = 2.1 Hz, 3H), 1.43 (s, 9H), 1.40 (s, 9H), 1.29 (s, 3H), 1.16 (s, 3H).

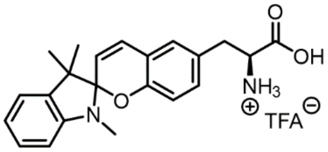

**(2*S*)-2-amino-3-(1',3',3'-trimethylspiro[chromene-2,2'-indolin]-6-yl) propanoic acid SpiroY: 31** (0.1 mmol, 1 eq) was dissolved in 5 mL DCM and excess TFA is added before the solution was stirred at rt for 2 days. The solvent was removed *in vacuo* and the residue was dissolved in a minimum amount of water followed by lyophilization. The compound was then triturated with $Et_2O$ and dried *in vacuo* to obtain the final product. The product was a mixture of spyropyran TFA salt and merocyanine double TFA salt. Red solid was obtained in quantitative yield. $^1$H-NMR (400 MHz, DMSO-$d_6$): $\delta$. (spiropyran) 8.38 - 8.12 (m, 3H), 7.13 - 7.02 (m, 3H), 6.97 (td, J = 7.0, 6.0, 3.4 Hz, 2H), 6.77 (t, J = 7.3 Hz, 1H), 6.65 (d, J = 8.3 Hz, 1H), 6.55 (d, J = 7.7 Hz, 1H), 5.77 (d, J = 10.3 Hz, 1H), 4.17 - 4.11 (m, 1H), 3.09 - 2.88 (m, 2H), 2.64 (d, J = 3.4 Hz, 3H), 1.21 (d, J = 2.7 Hz, 3H), 1.09 (s, 3H). (merocyanine) 8.47 (d, J = 16.4 Hz, 1H), 8.42 (s, 3H), 8.00 (d, J = 2.3 Hz, 1H), 7.95 - 7.82 (m, 2H), 7.73 - 7.56 (m, 3H), 7.35 (dd, J = 8.5, 2.2 Hz, 1H), 7.04 (d, J = 8.5 Hz, 1H), 4.20 (s, 1H), 4.09 (s, 3H), 3.19 - 3.07 (m, 2H), 1.76 (s, 6H). $^{13}$C-NMR (101 MHz, DMSO-$d_6$): $\delta$ 181.7, 170.5, 170.4, 158.5, 158.4, 158.1, 153.2, 148.4, 147.8, 143.2, 141.9, 136.5, 136.3, 130.9, 129.3, 129.2, 129.0, 128.0, 127.5, 126.4, 122.9, 121.5, 121.2, 119.6, 119.0, 118.6, 118.4, 117.1, 115.1, 114.6, 112.0, 106.8, 104.0, 88.7, 53.2, 51.9, 51.3, 35.0, 34.2, 28.6, 26.0, 25.7, 20.0. HRMS (ESI-TOF) m/z calculated for $[C_{22}H_{25}N_2O_3]^+$ $[MH]^+$ 365.1860, found 365.1862 (z=1), 183.0973 (z=2).

##### 2.1.3 $^1$H-NMR Spectra

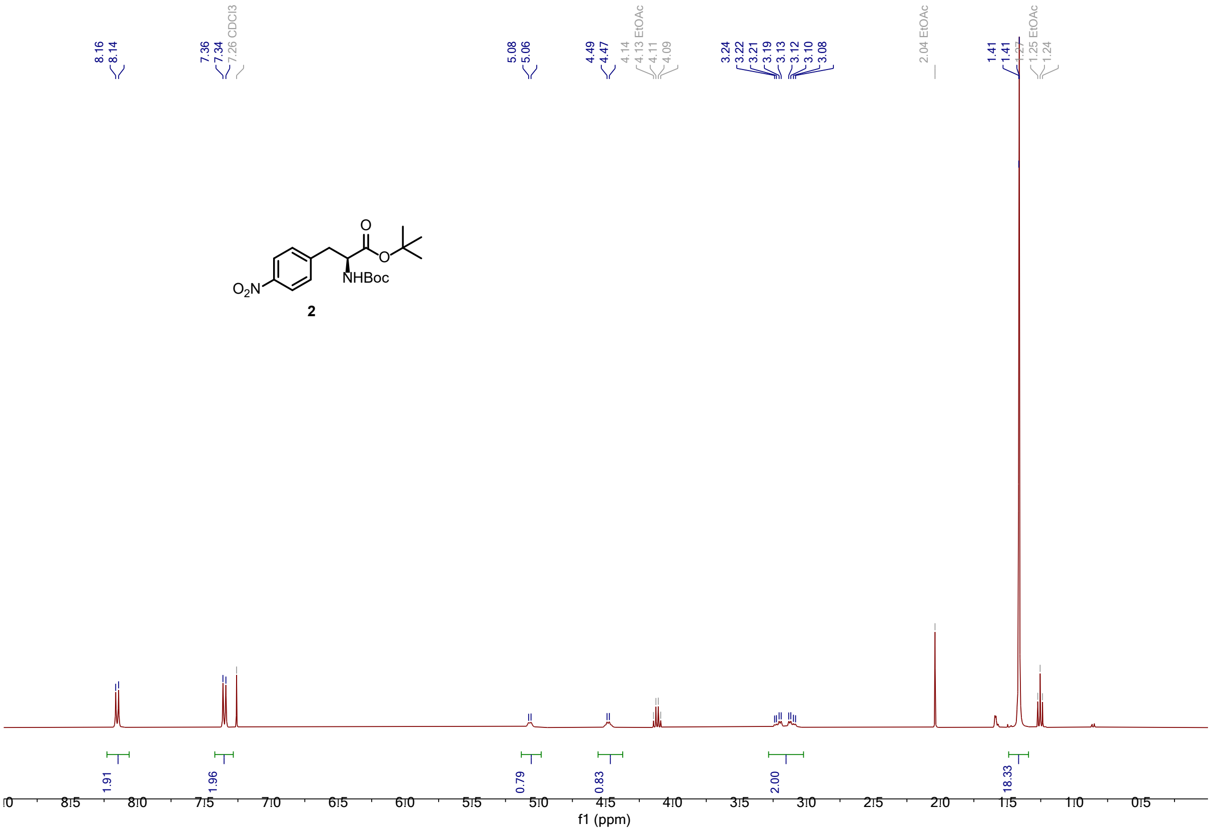

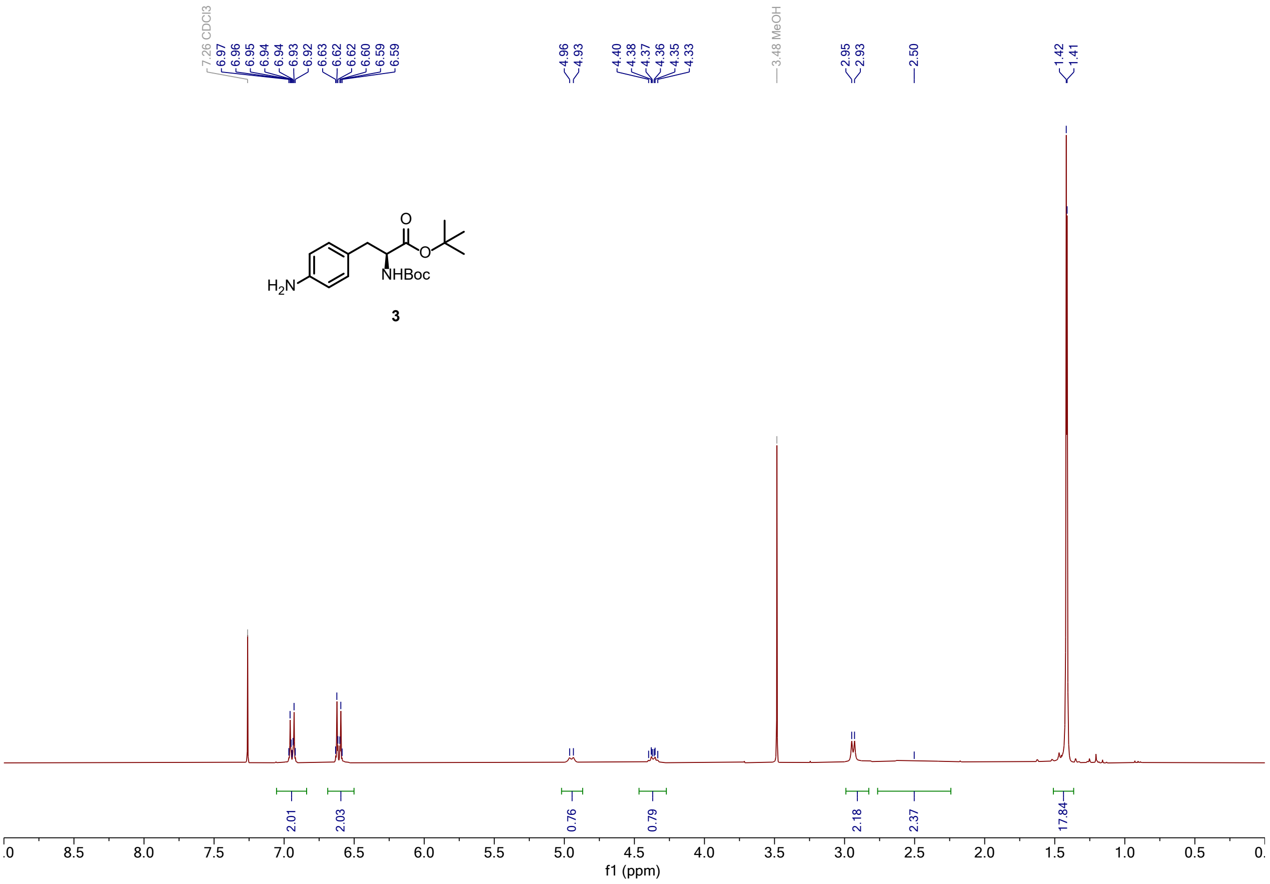

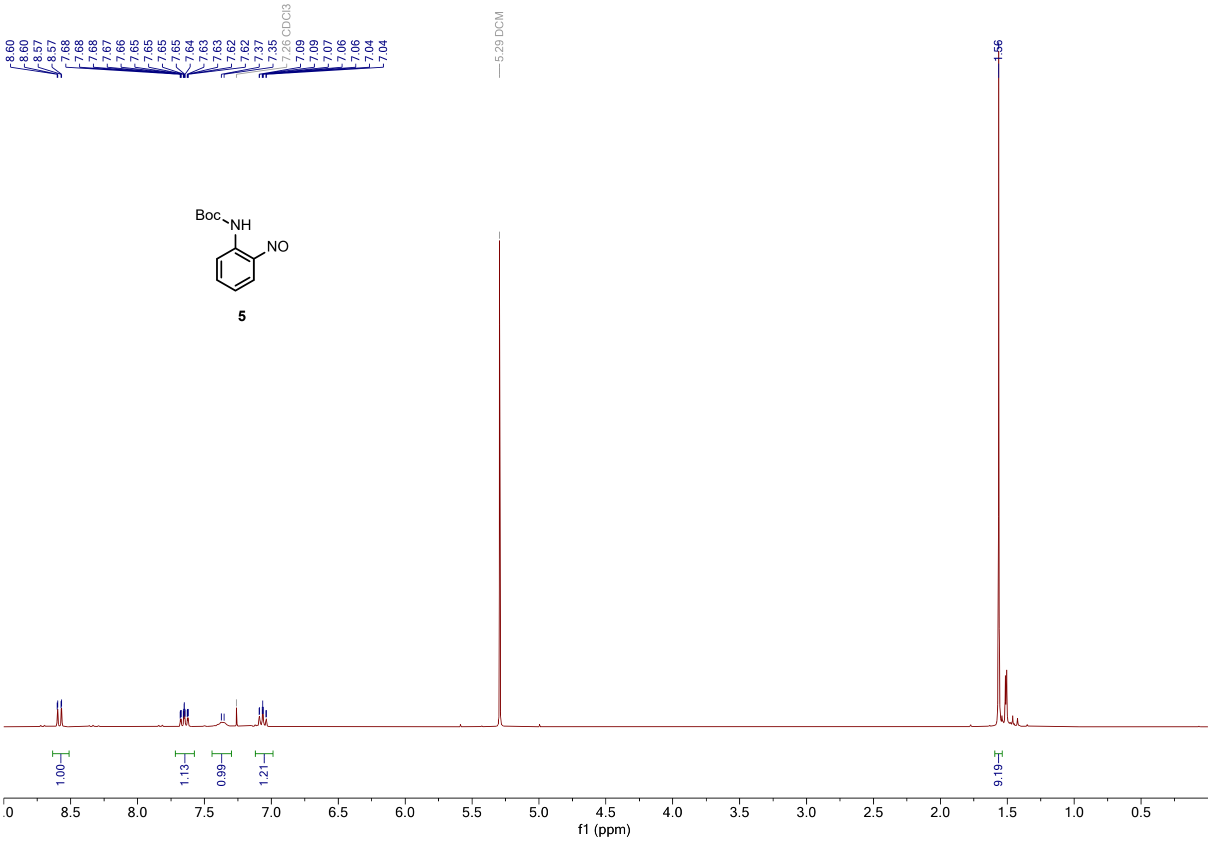

Boc
NH
NO
5
f1 (ppm)

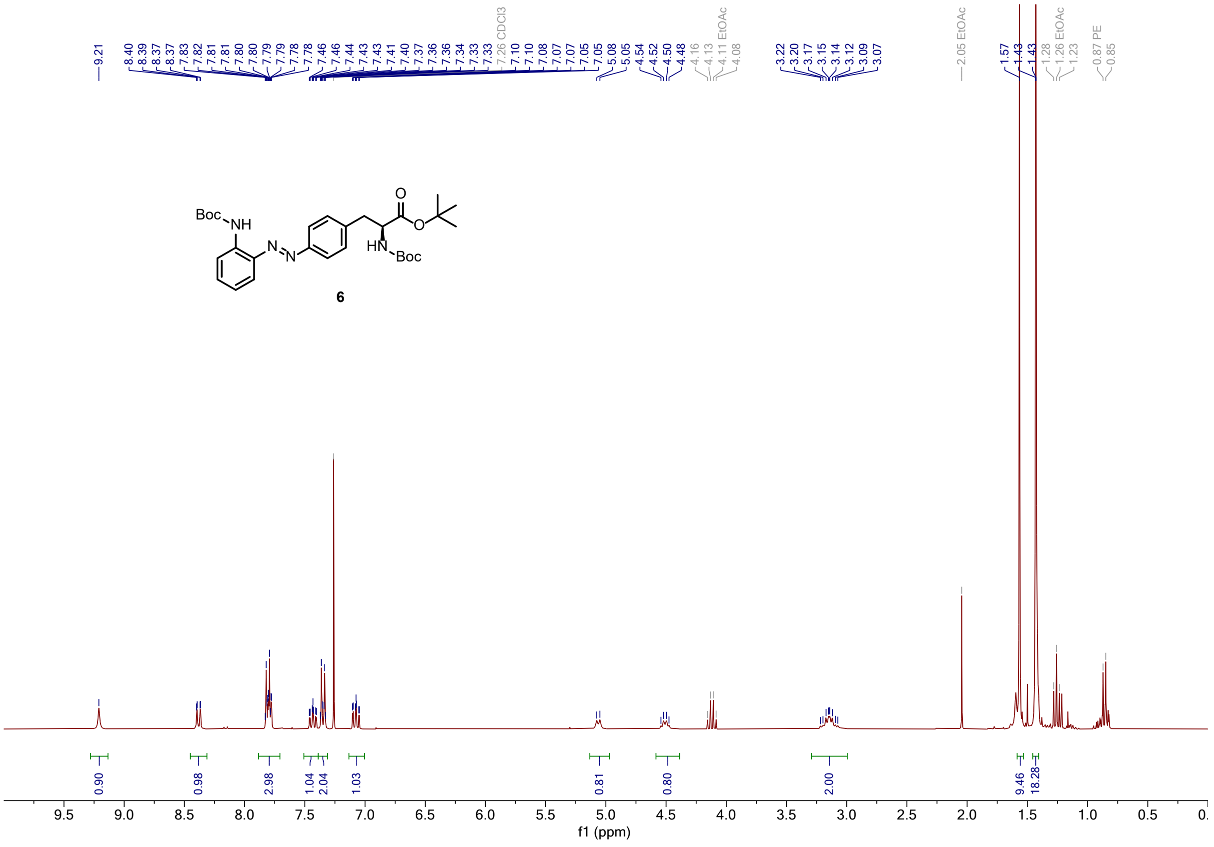

Boc
NH
N
N
O
O
HN
Boc
6
f1 (ppm)

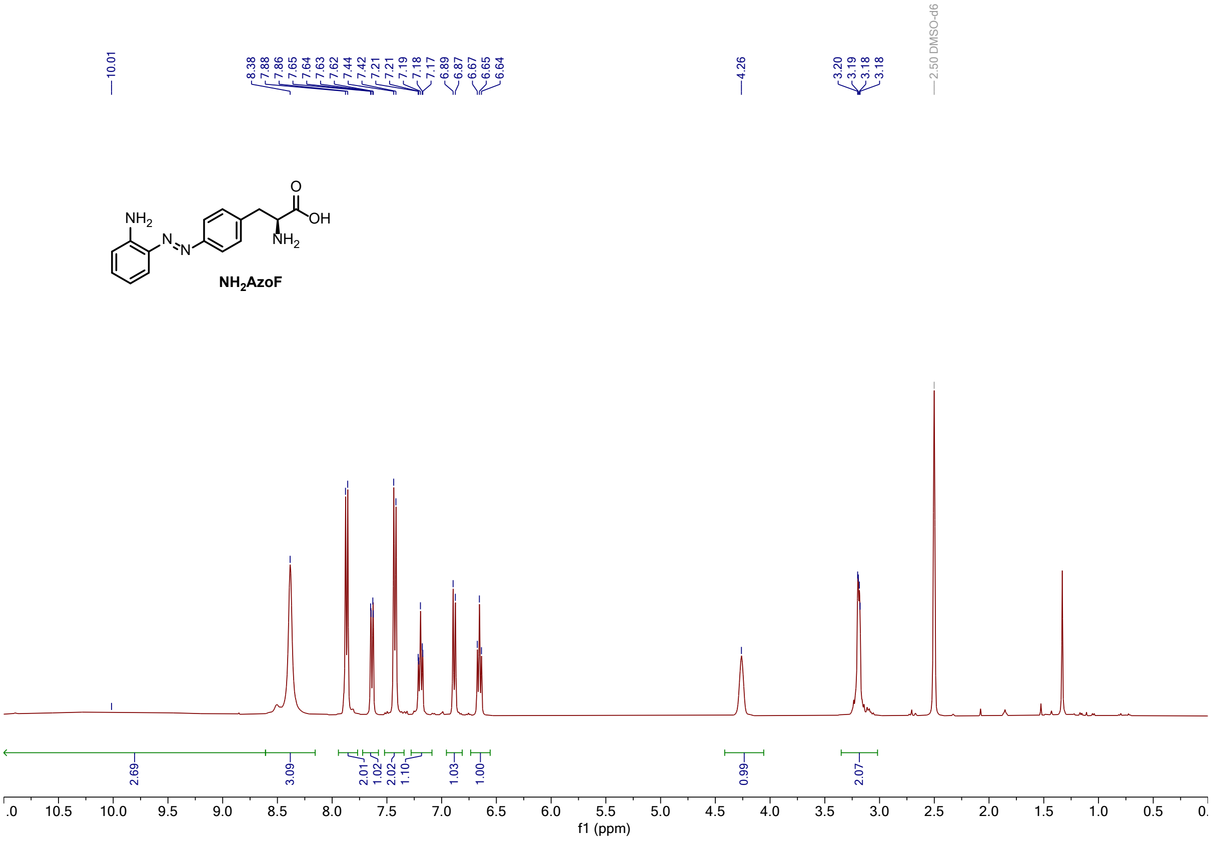
10.01
8.38
7.88
7.86
7.65
7.64
7.63
7.62
7.44
7.42
7.21
7.21
7.19
7.18
7.17
6.89
6.87
6.67
6.65
6.64
4.26
3.20
3.19
3.18
3.18
2.50 DMSO-d6
NH2
N
N
O
OH
NH2
NH2AzoF
2.69
3.09
2.01
1.02
2.02
1.10
1.03
1.00
0.99
2.07
f1 (ppm)

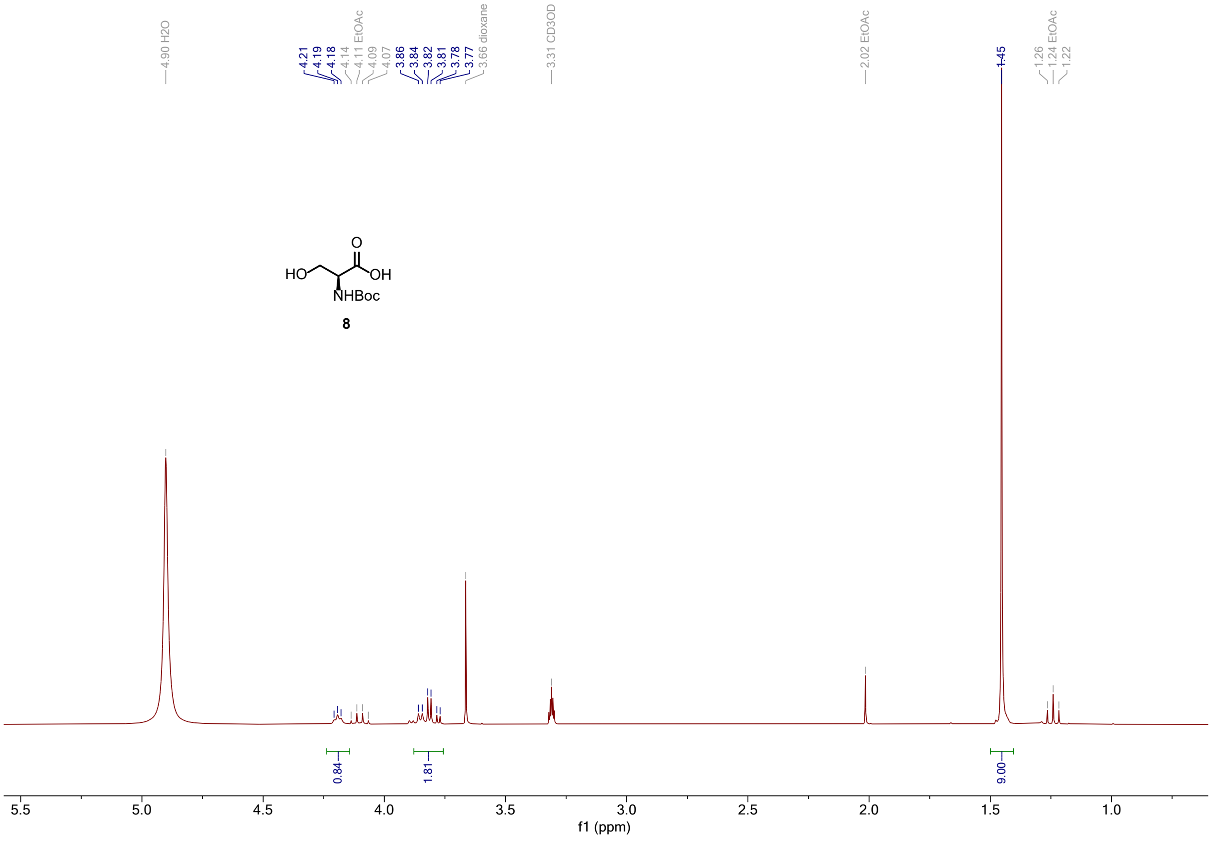
4.90 H2O
4.21
4.19
4.18
4.14
4.11 EtOAc
4.09
4.07
3.86
3.84
3.82
3.81
3.78
3.77
3.66 dioxane
3.31 CD3OD
2.02 EtOAc
1.45
1.26
1.24 EtOAc
1.22
O
HO
OH
NHBoc
8
0.84
1.81
9.00
f1 (ppm)

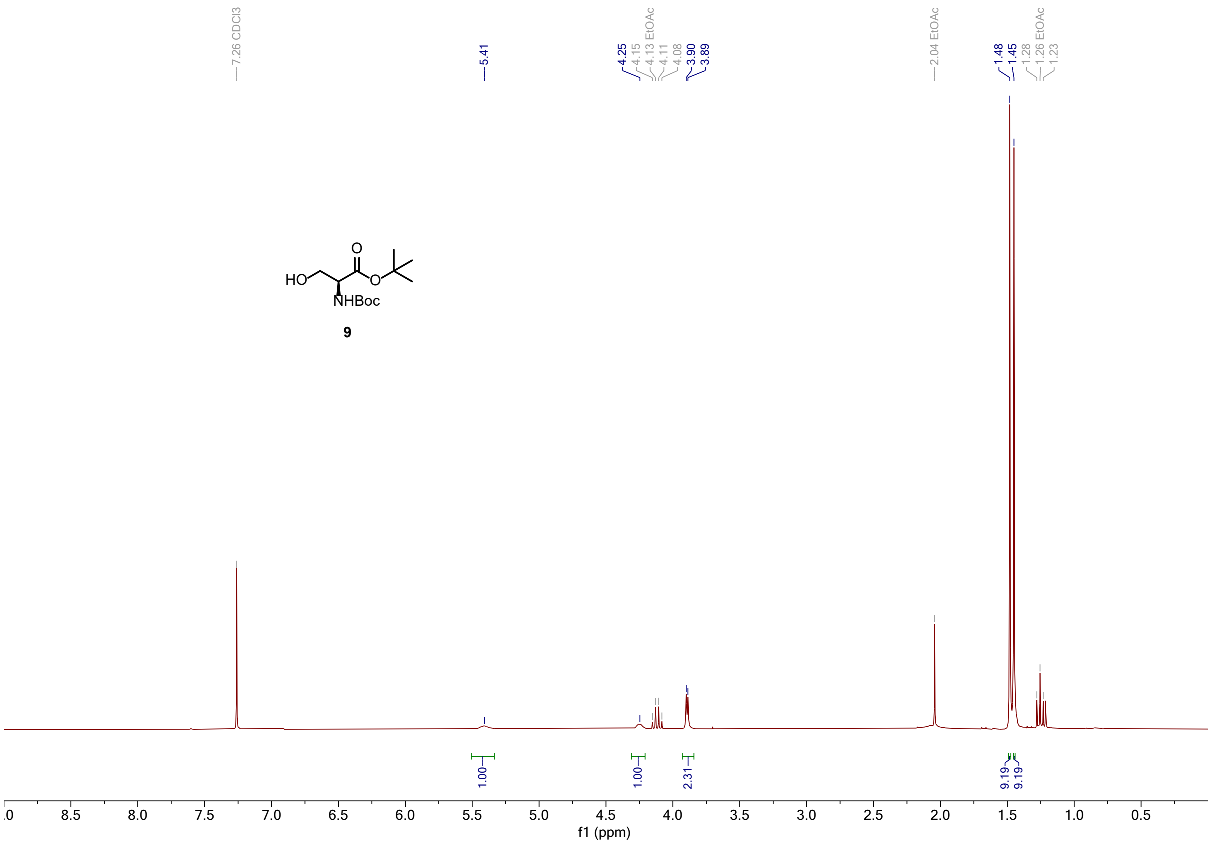
7.26 CDCl3
5.41
4.25
4.15
4.13 EtOAc
4.11
4.08
3.90
3.89
2.04 EtOAc
1.48
1.45
1.28
1.26 EtOAc
1.23
HO
O
O
NHBoc
9
1.00
1.00
2.31
9.19
9.19
.0 8.5 8.0 7.5 7.0 6.5 6.0 5.5 5.0 4.5 4.0 3.5 3.0 2.5 2.0 1.5 1.0 0.5
f1 (ppm)

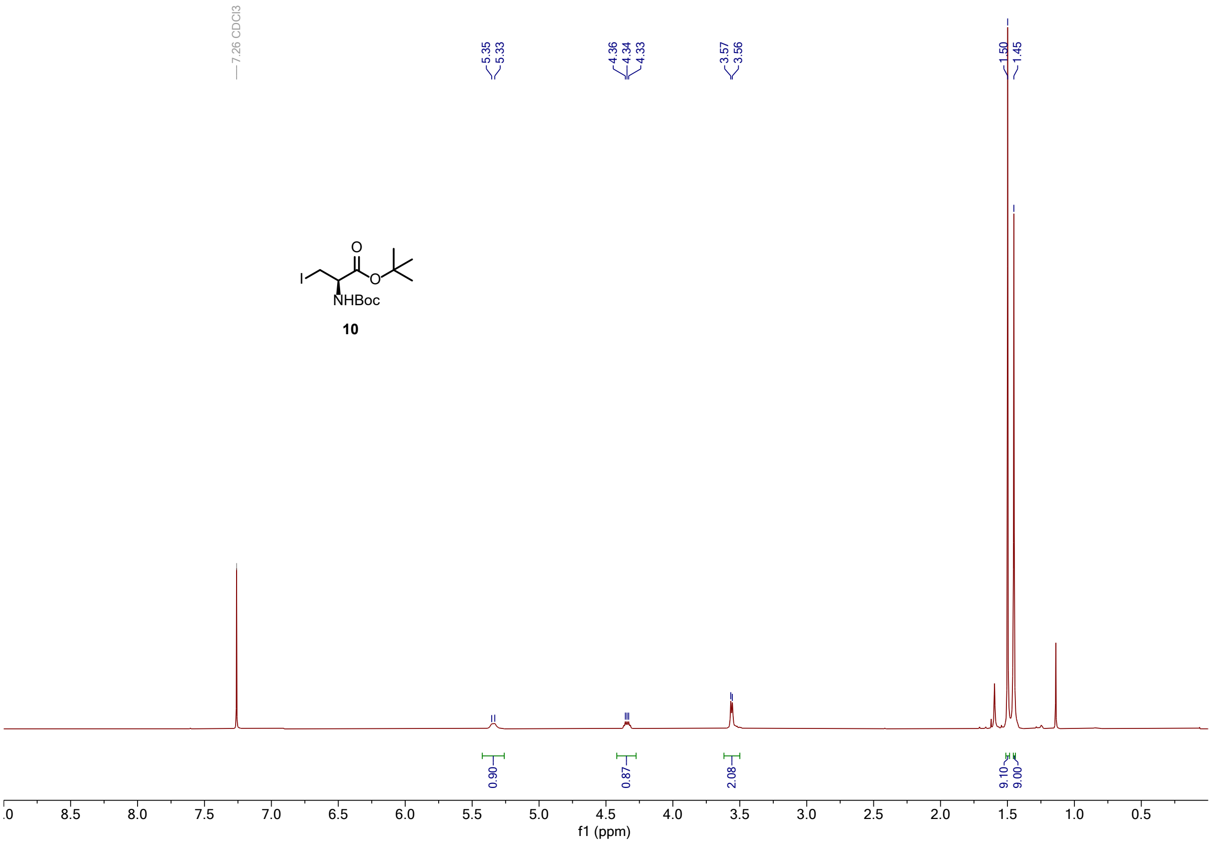
7.26 CDCl3
5.35
5.33
4.36
4.34
4.33
3.57
3.56
1.50
1.45
I
O
O
NHBoc
10
0.90
0.87
2.08
9.10
9.00
.0 8.5 8.0 7.5 7.0 6.5 6.0 5.5 5.0 4.5 4.0 3.5 3.0 2.5 2.0 1.5 1.0 0.5
f1 (ppm)

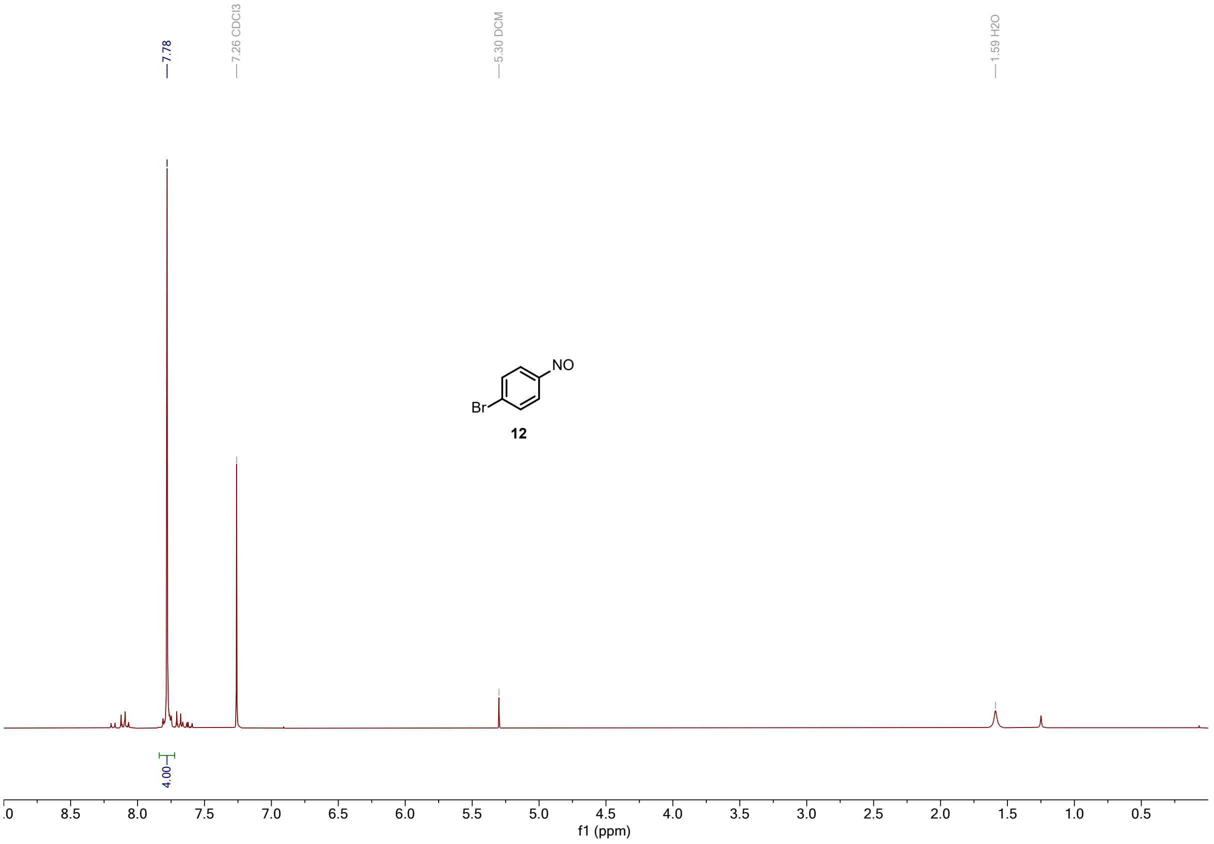
7.78
7.26 CDCl3
5.30 DCM
1.59 H2O
NO
Br
12
4.00
f1 (ppm)

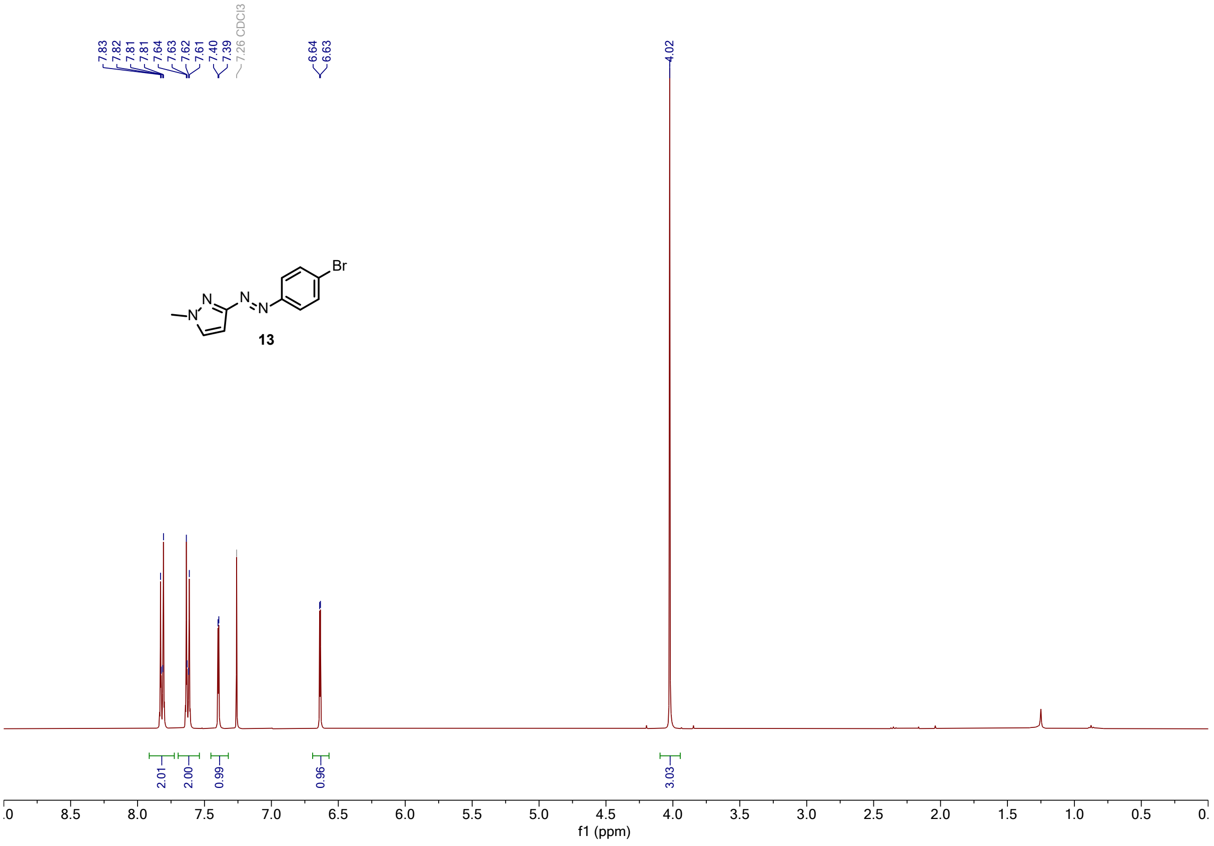
7.83
7.82
7.81
7.81
7.64
7.63
7.62
7.61
7.40
7.39
7.26 CDCl3
6.64
6.63
4.02
N
N
N
N
Br
13
2.01
2.00
0.99
0.96
3.03
f1 (ppm)

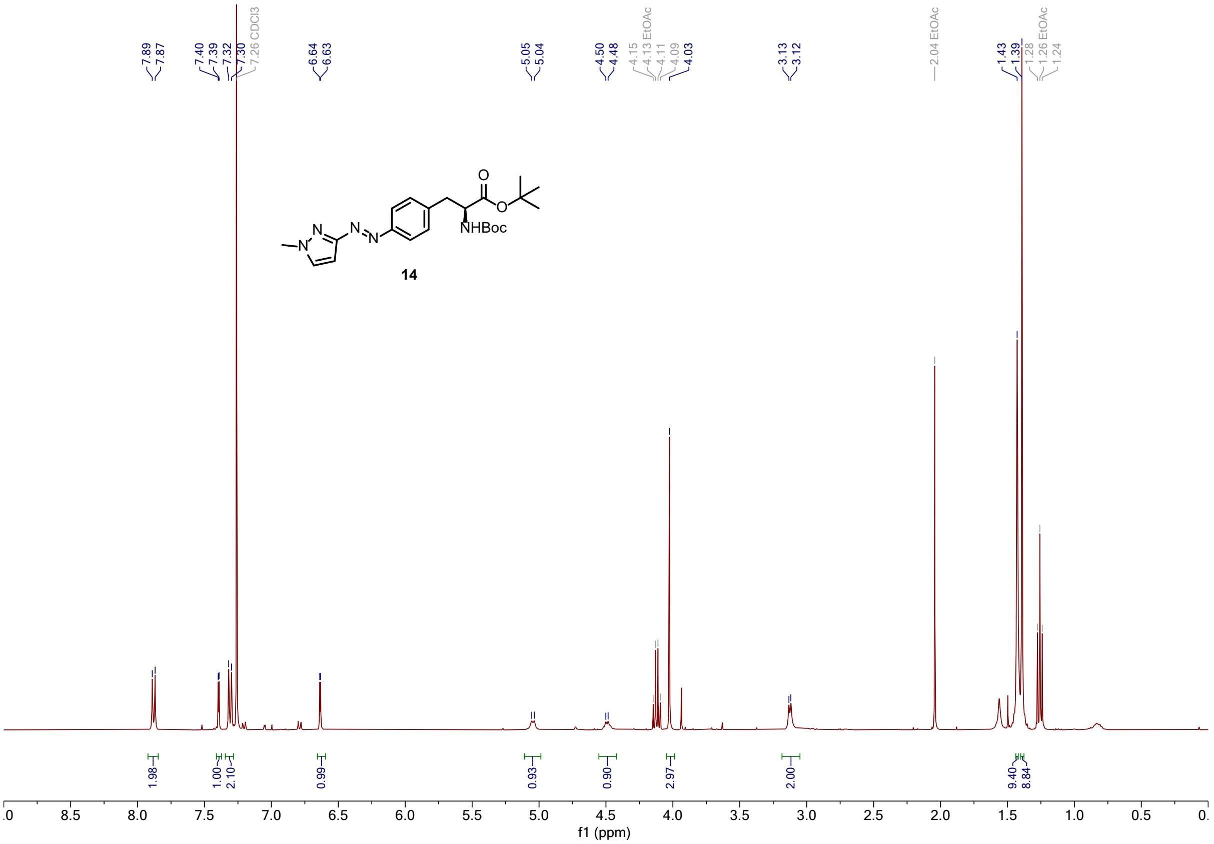
7.89
7.87
7.40
7.39
7.32
7.30
7.26 CDCl3
6.64
6.63
5.05
5.04
4.50
4.48
4.15
4.13 EtOAc
4.11
4.09
4.03
3.13
3.12
2.04 EtOAc
1.43
1.39
1.28
1.26 EtOAc
1.24
NHBoc
14
1.98
1.00
2.10
0.99
0.93
0.90
2.97
2.00
9.40
8.84
f1 (ppm)

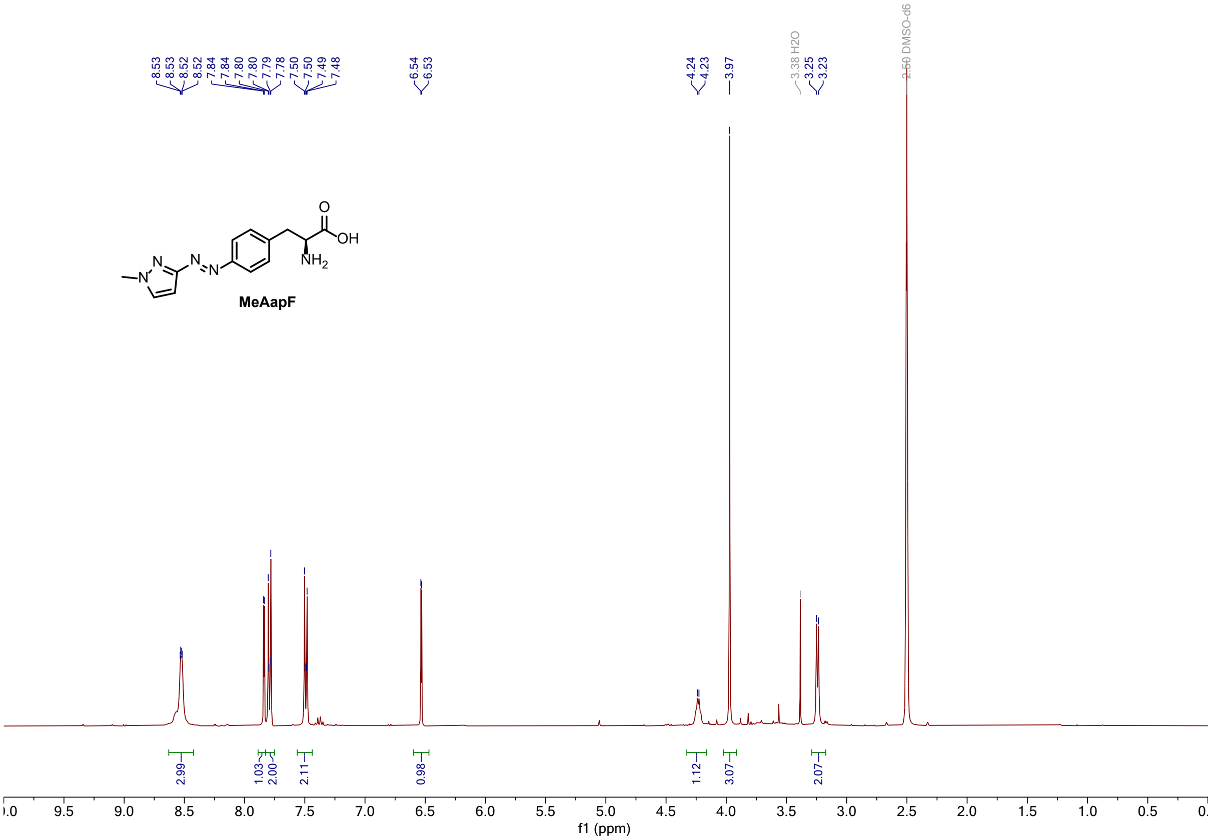
8.53
8.53
8.52
8.52
7.84
7.84
7.80
7.79
7.78
7.50
7.50
7.49
7.48
6.54
6.53
4.24
4.23
3.97
3.38 H2O
3.25
3.23
2.50 DMSO-d6
OH
MeAapF
2.99
1.03
2.00
2.11
0.98
1.12
3.07
2.07
f1 (ppm)

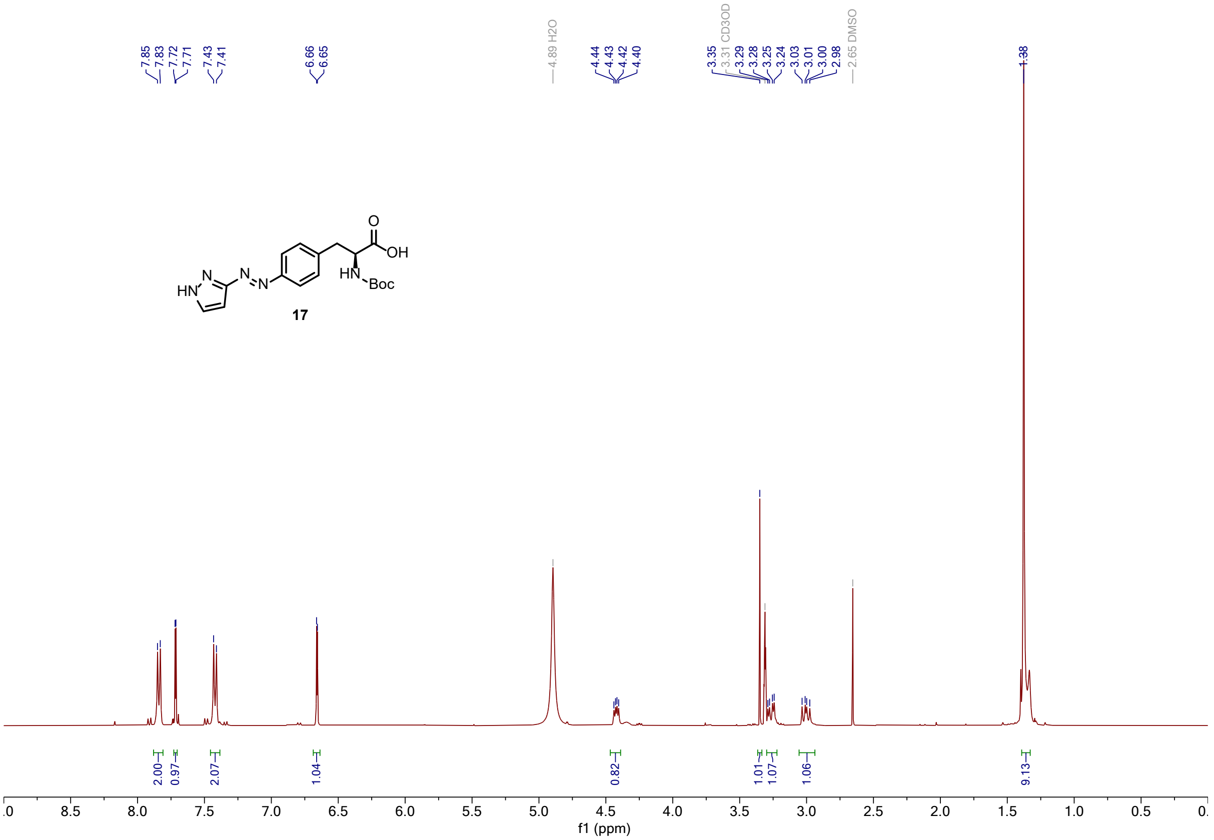

7.85 7.83 7.72 7.71 7.43 7.41
6.66 6.65
4.89 H2O
4.44 4.43 4.42 4.40
3.35 3.31 CD3OD 3.29 3.28 3.25 3.24 3.03 3.01 3.00 2.98
2.65 DMSO
1.38
HN N N N O OH HN Boc
17
2.00 0.97 2.07 1.04 0.82 1.01 1.07 1.06 9.13
f1 (ppm)

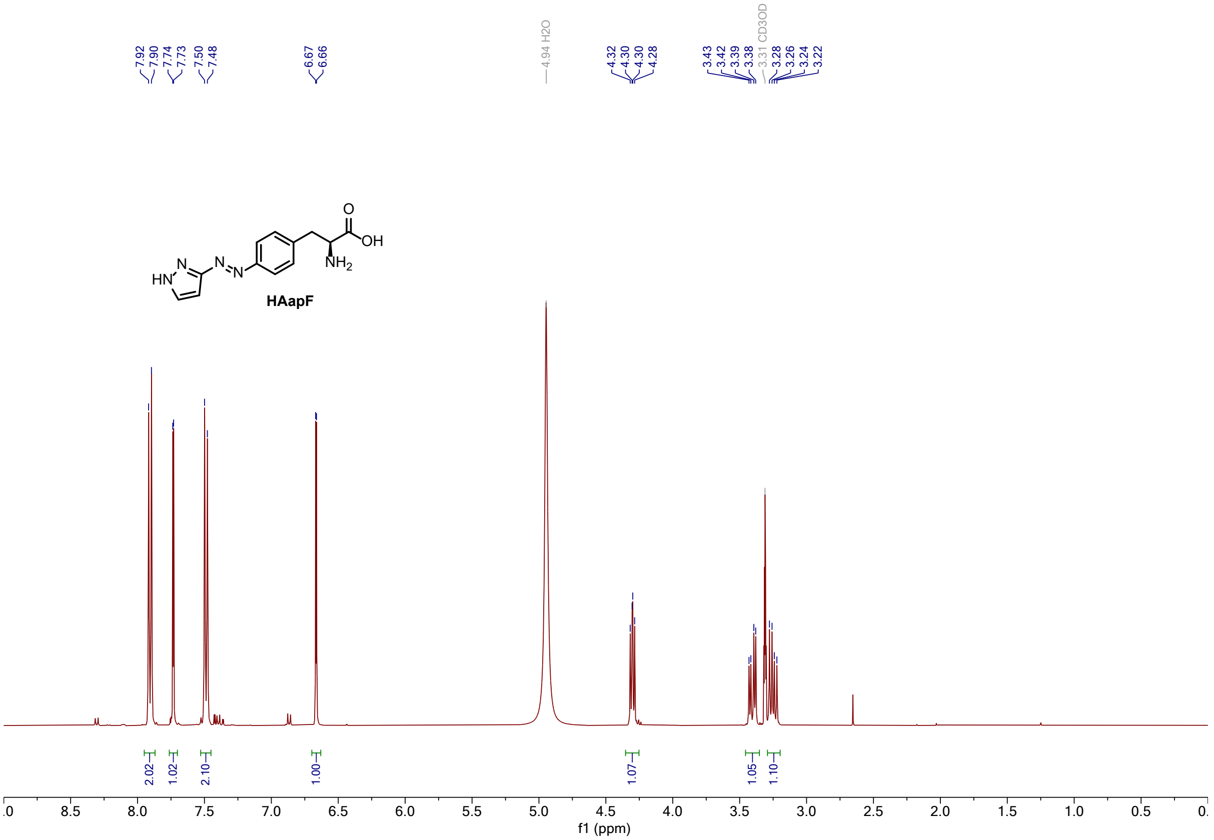

7.92 7.90 7.74 7.73 7.50 7.48
6.67 6.66
4.94 H2O
4.32 4.30 4.30 4.28
3.43 3.42 3.39 3.38 3.31 CD3OD 3.28 3.26 3.24 3.22
HN N N N O OH NH2
HAapF
2.02 1.02 2.10 1.00 1.07 1.05 1.10
f1 (ppm)

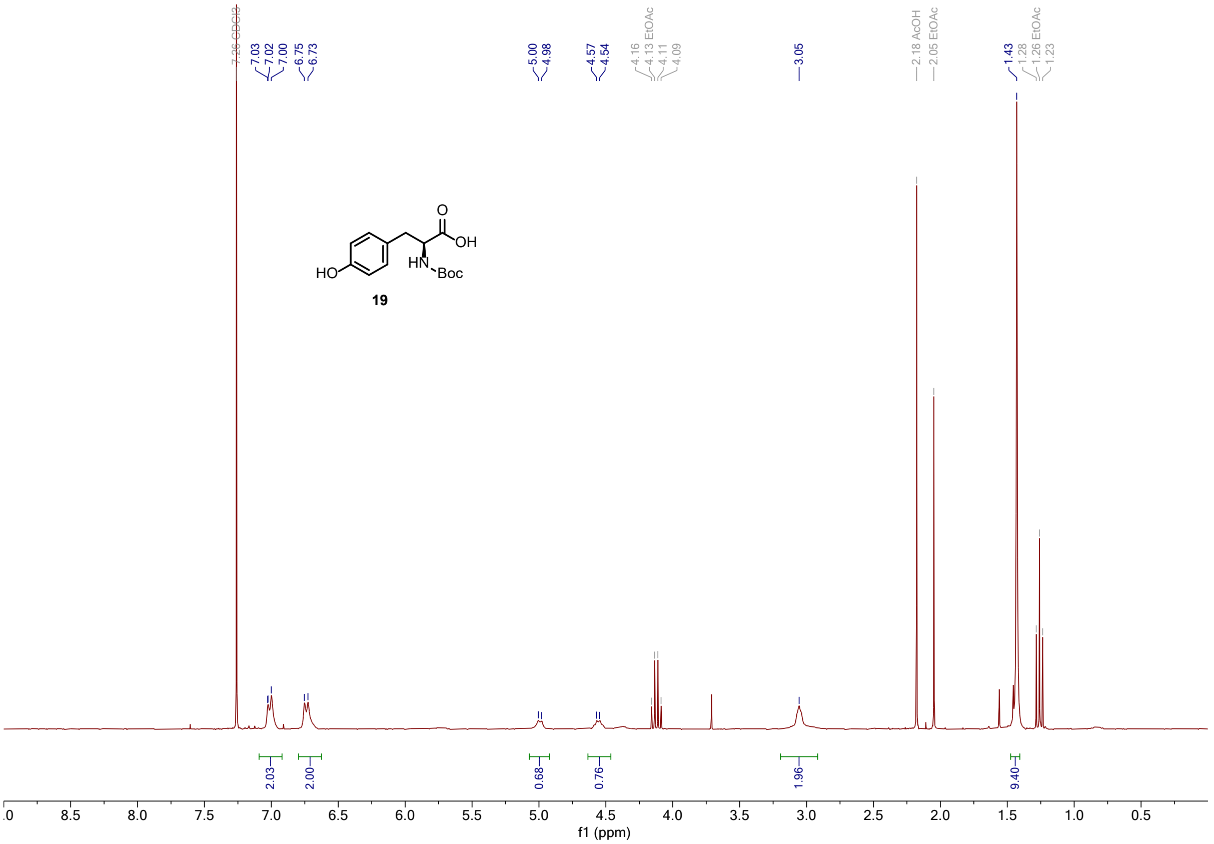
7.26 CDCl3
7.03
7.02
7.00
6.75
6.73
5.00
4.98
4.57
4.54
4.16
4.13 EtOAc
4.11
4.09
3.05
2.18 AcOH
2.05 EtOAc
1.43
1.28
1.26 EtOAc
1.23
O
OH
HO
HN
Boc
19
2.03
2.00
0.68
0.76
1.96
9.40
f1 (ppm)

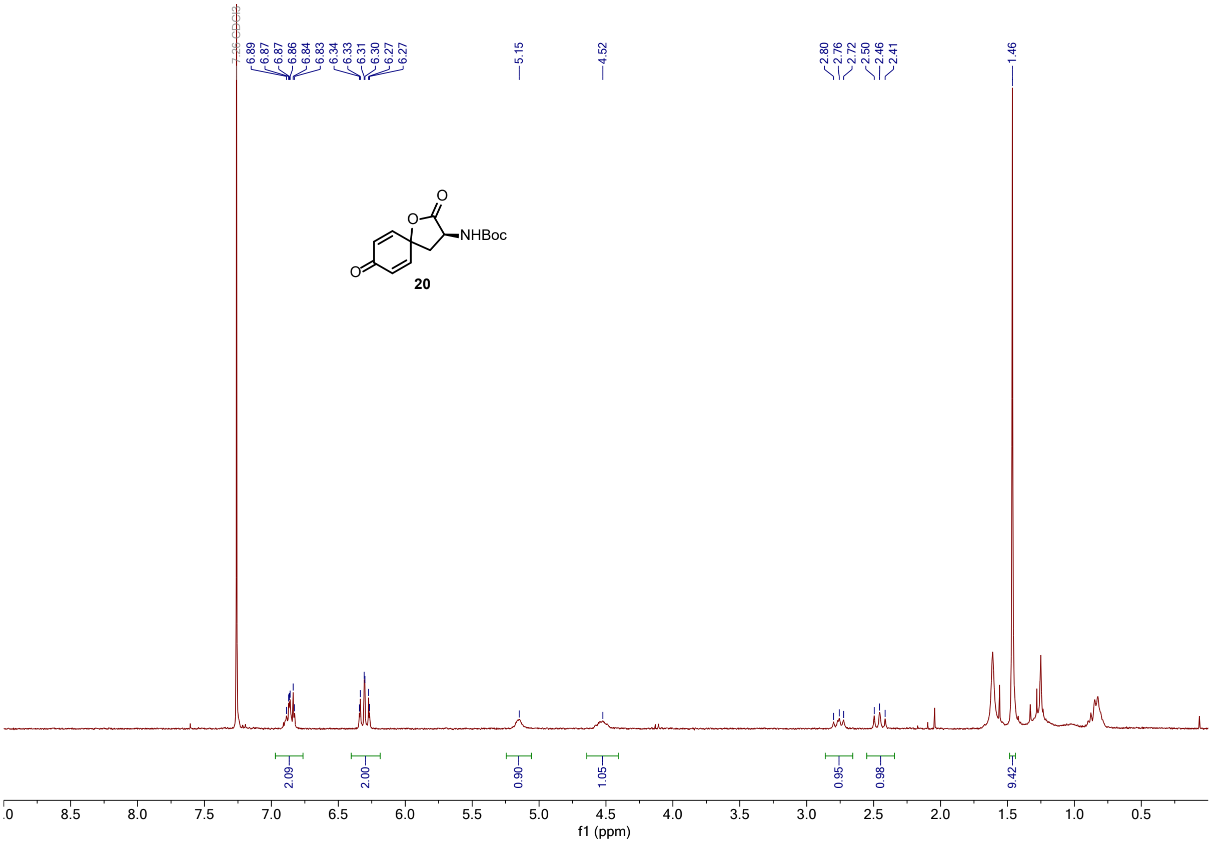
7.26 CDCl3
6.89
6.87
6.87
6.86
6.84
6.83
6.34
6.33
6.31
6.30
6.27
6.27
5.15
4.52
2.80
2.76
2.72
2.50
2.46
2.41
1.46
O
O
NHBoc
O
20
2.09
2.00
0.90
1.05
0.95
0.98
9.42
f1 (ppm)

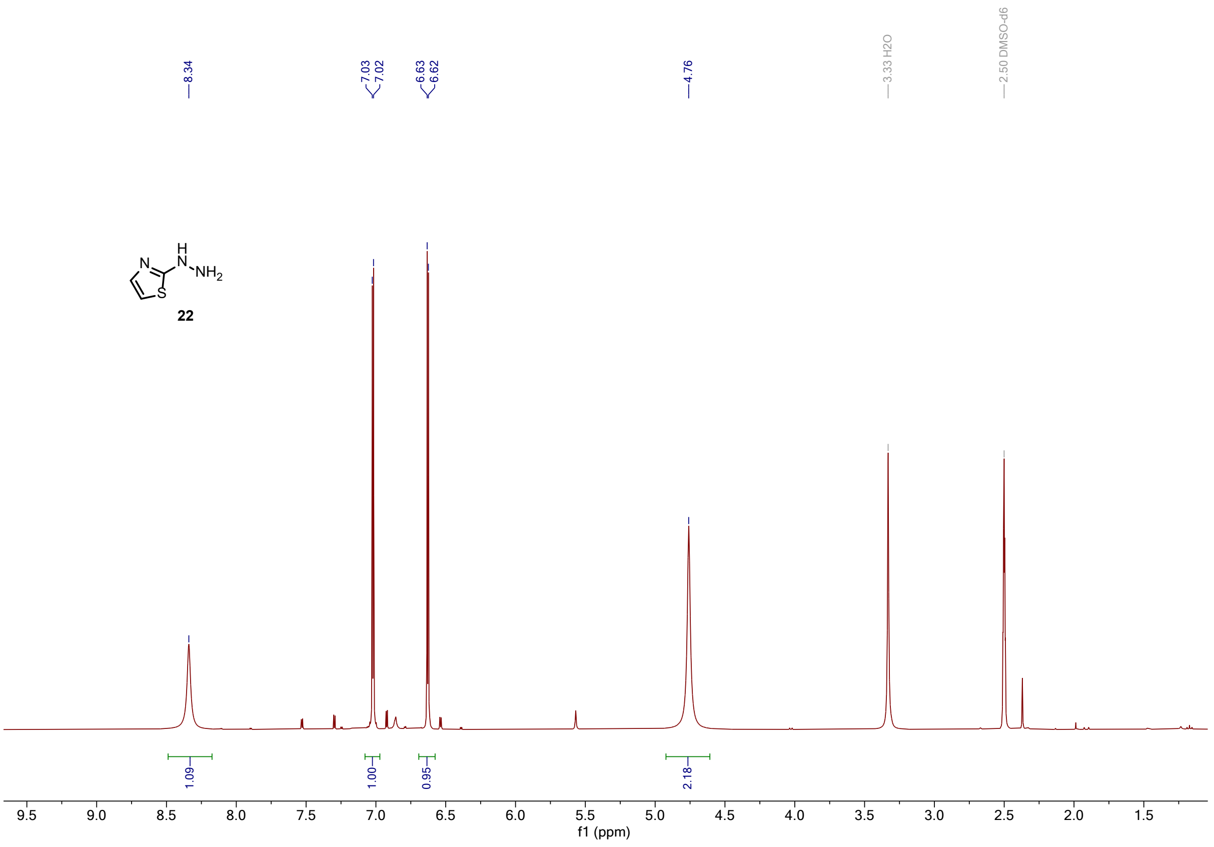
8.34
7.03
7.02
6.63
6.62
4.76
3.33 H2O
2.50 DMSO-d6
22
1.09
1.00
0.95
2.18
f1 (ppm)

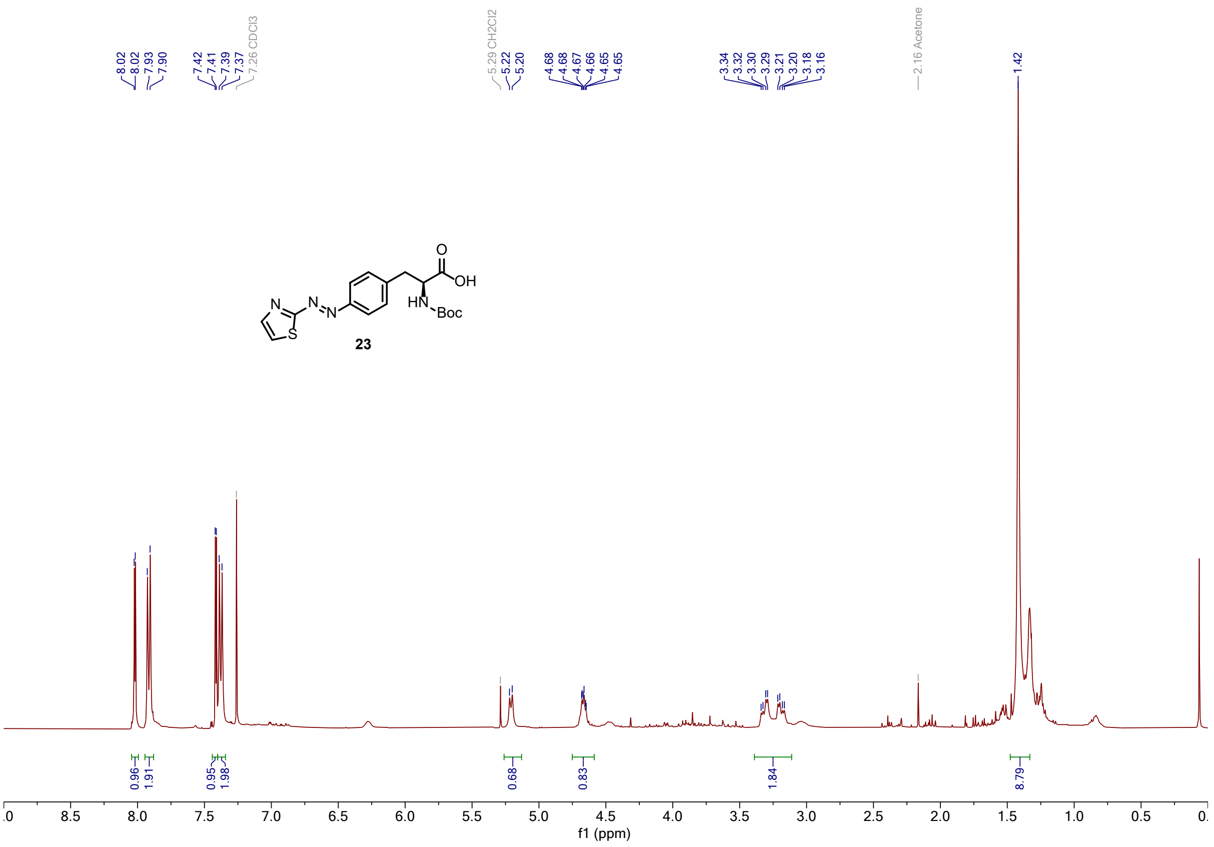
8.02
8.02
7.93
7.90
7.42
7.41
7.39
7.37
7.26 CDCl3
5.29 CH2Cl2
5.22
5.20
4.68
4.68
4.67
4.66
4.65
4.65
3.34
3.32
3.30
3.29
3.21
3.20
3.18
3.16
2.16 Acetone
1.42
23
0.96
1.91
0.95
1.98
0.68
0.83
1.84
8.79
f1 (ppm)

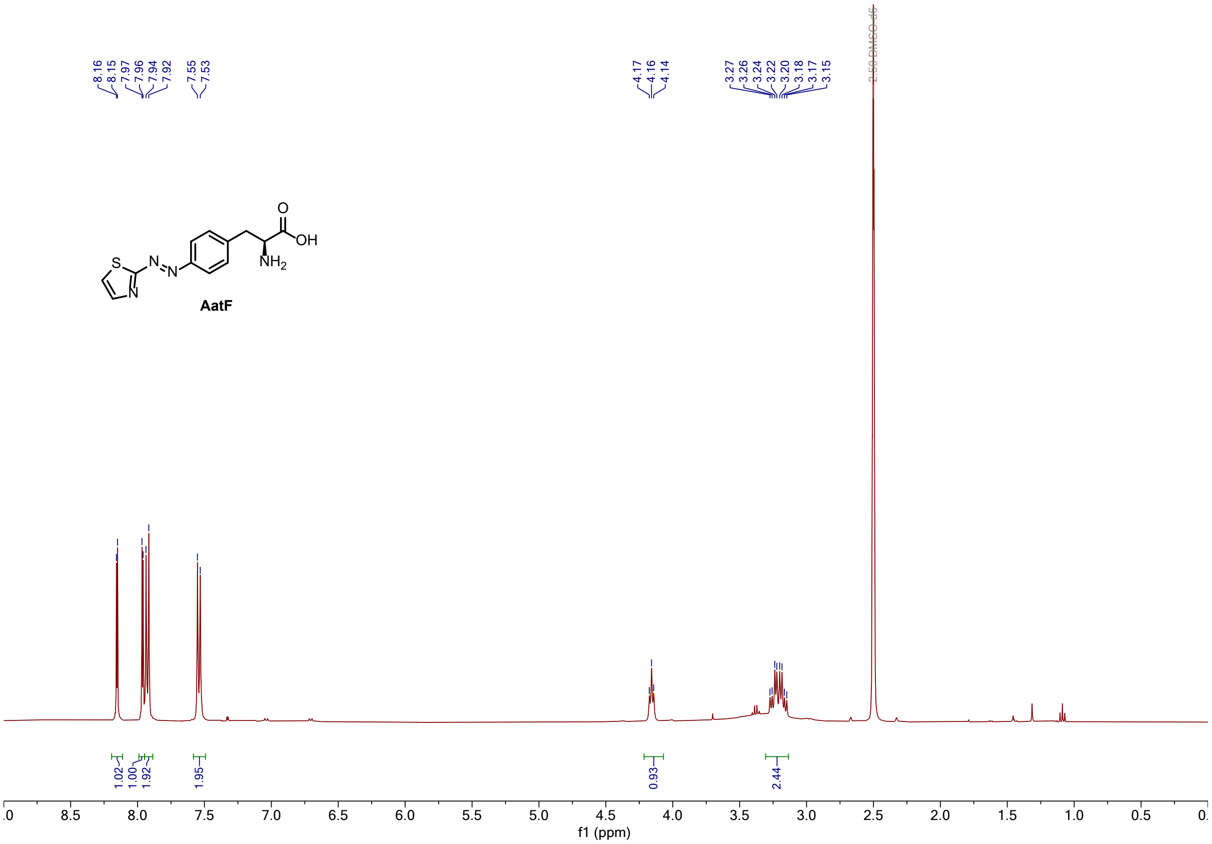
8.16
8.15
7.97
7.96
7.94
7.92
7.55
7.53
4.17
4.16
4.14
3.27
3.26
3.24
3.22
3.20
3.18
3.17
3.15
2.50 DMSO-d6
S
N
N
N
O
OH
NH2
AatF
1.02
1.00
1.92
1.95
0.93
2.44
f1 (ppm)

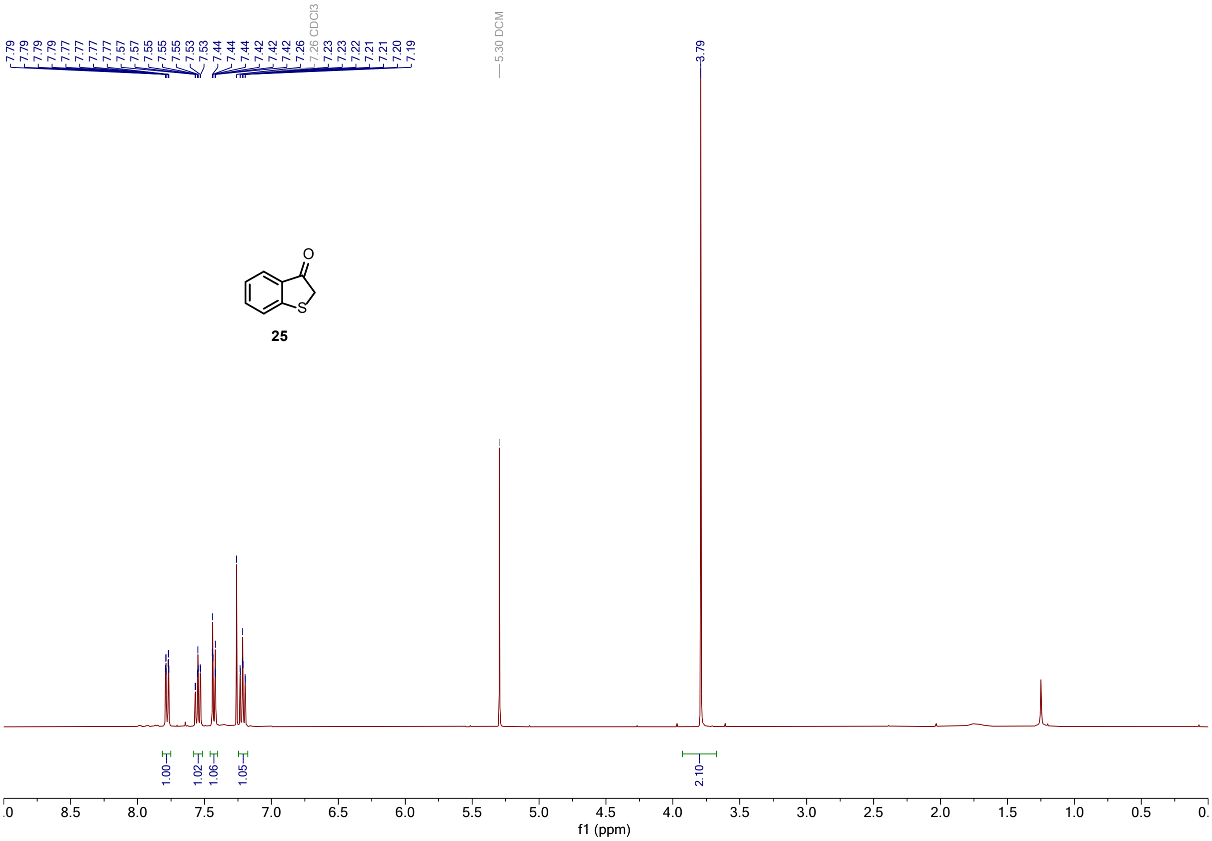
7.79
7.79
7.79
7.79
7.77
7.77
7.77
7.77
7.57
7.57
7.55
7.55
7.53
7.53
7.44
7.44
7.44
7.42
7.42
7.42
7.26 CDCl3
7.23
7.23
7.22
7.21
7.21
7.20
7.19
5.30 DCM
3.79
O
S
25
1.00
1.02
1.06
1.05
2.10
f1 (ppm)

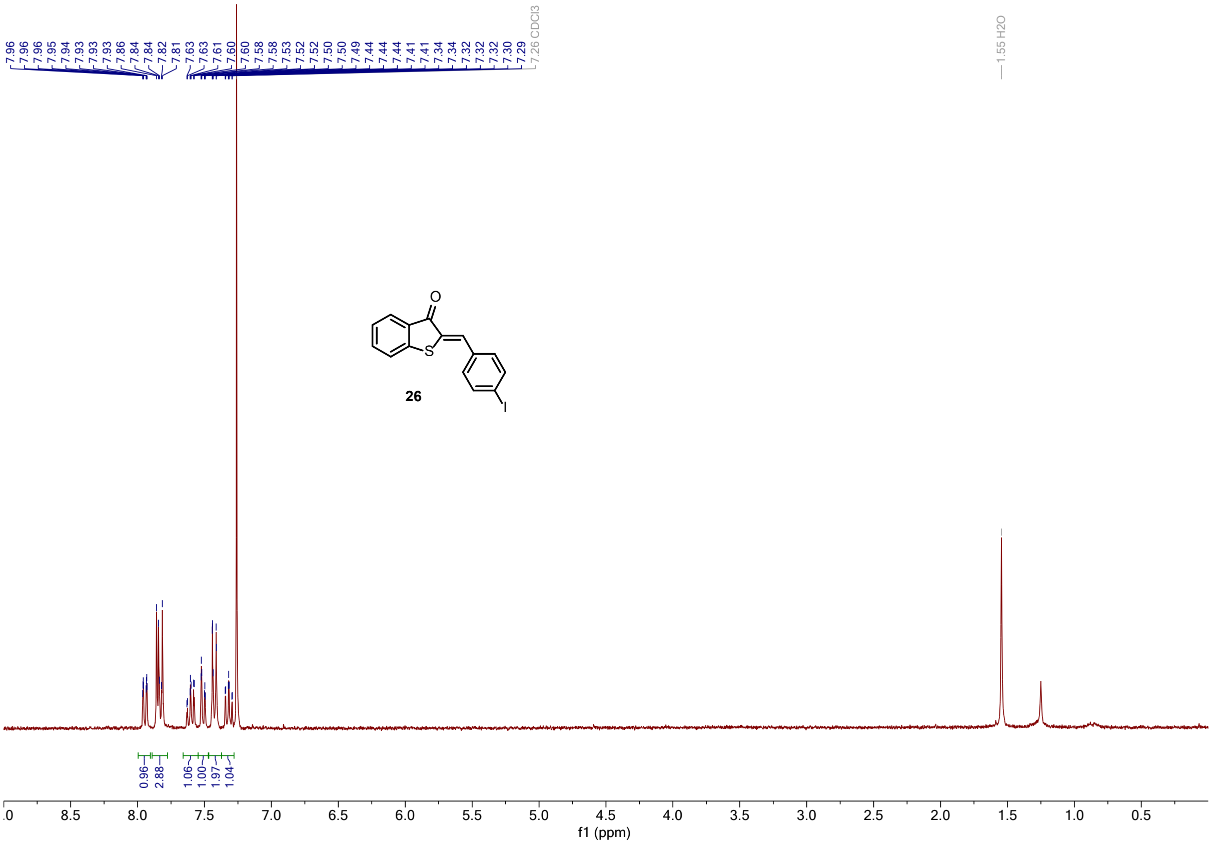
7.96 7.96 7.96 7.95 7.94 7.93 7.93 7.93 7.86 7.84 7.84 7.82 7.81 7.63 7.63 7.61 7.60 7.60 7.60 7.58 7.58 7.53 7.52 7.52 7.50 7.50 7.49 7.44 7.44 7.44 7.41 7.41 7.34 7.34 7.32 7.32 7.30 7.29
7.26 CDCl3
1.55 H2O
O
S
I
26
0.96 2.88 1.06 1.00 1.97 1.04
f1 (ppm)

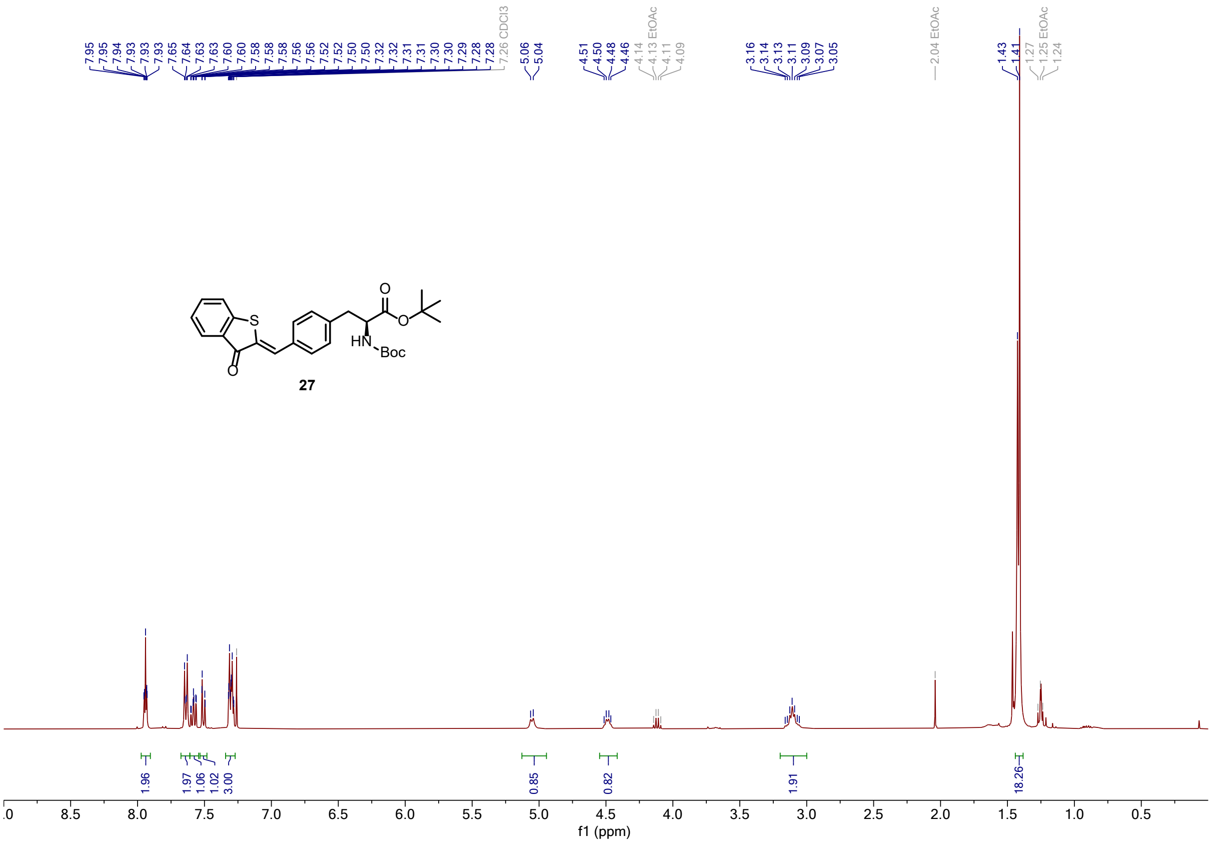
7.95 7.95 7.94 7.93 7.93 7.65 7.64 7.63 7.63 7.60 7.60 7.58 7.58 7.56 7.52 7.52 7.50 7.50 7.32 7.32 7.31 7.30 7.29 7.28 7.28
7.26 CDCl3
5.06 5.04
4.51 4.50 4.48 4.46
4.14 4.13 EtOAc 4.11 4.09
3.16 3.14 3.13 3.11 3.09 3.07 3.05
2.04 EtOAc
1.43 1.41
1.27 1.25 EtOAc 1.24
S
O
HN
Boc
27
1.96 1.97 1.06 1.02 3.00 0.85 0.82 1.91 18.26
f1 (ppm)

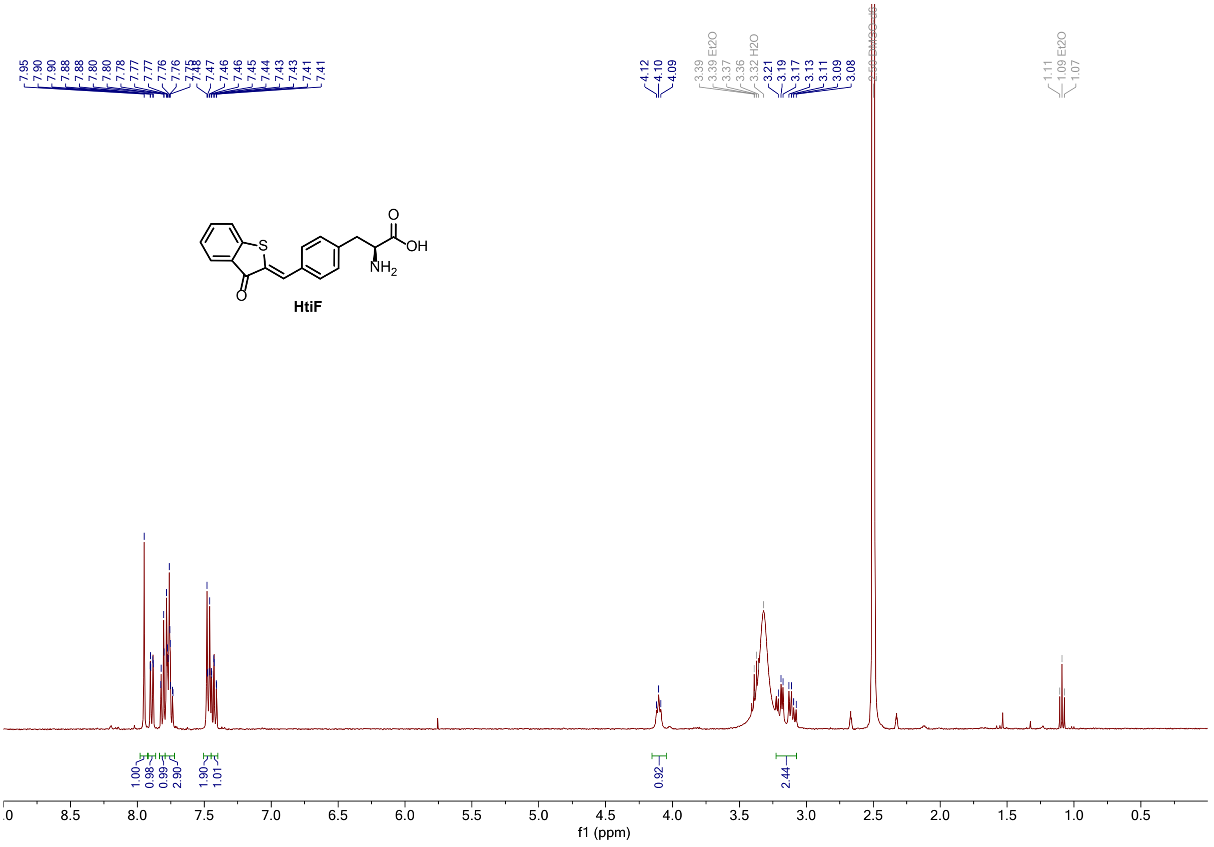

HtiF

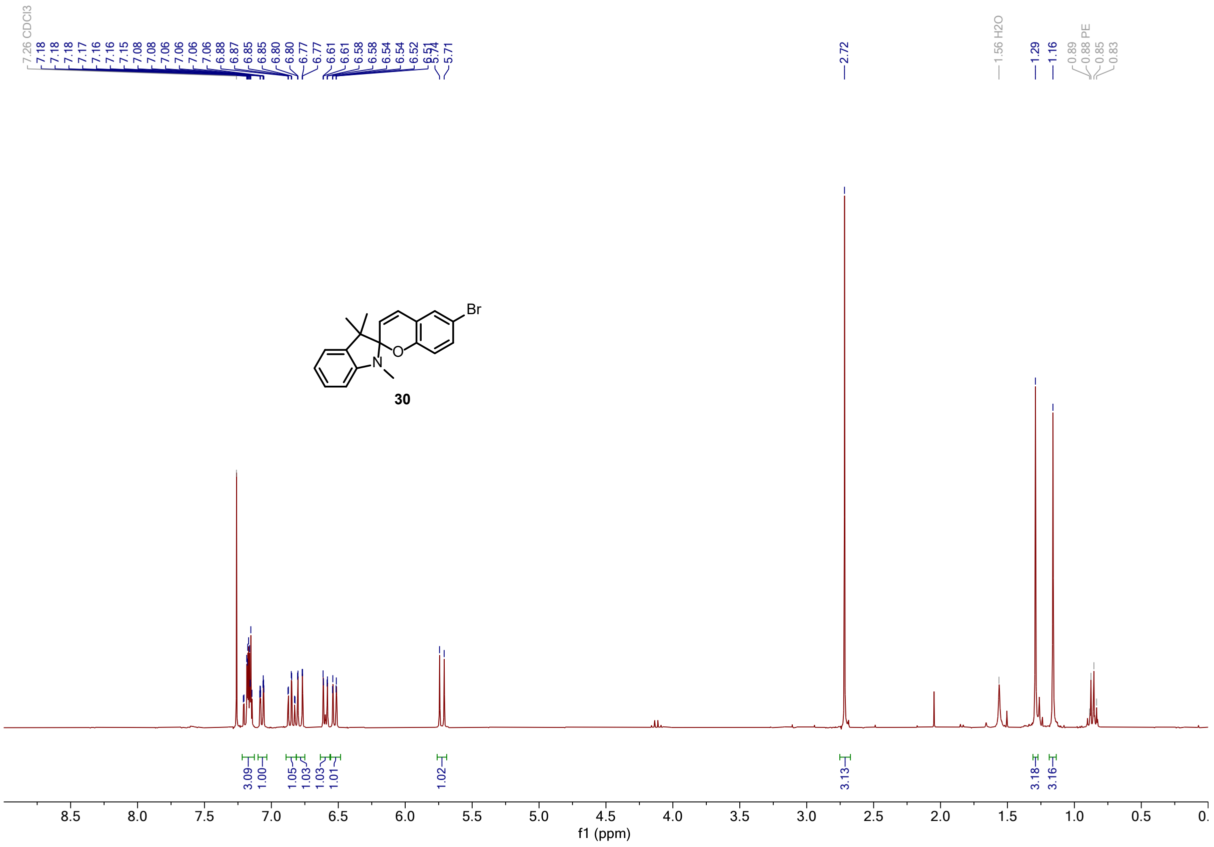

30

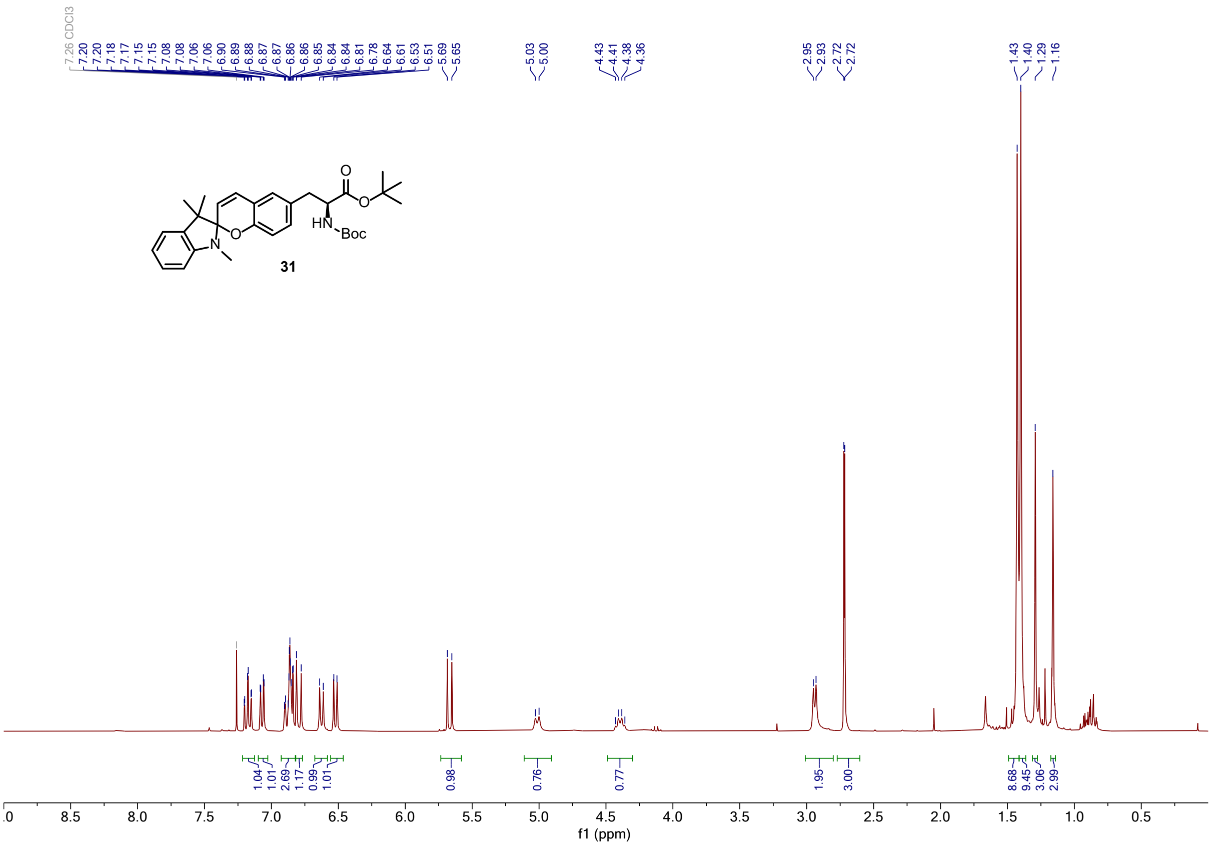
7.26 CDCl3
HN
Boc
31
f1 (ppm)

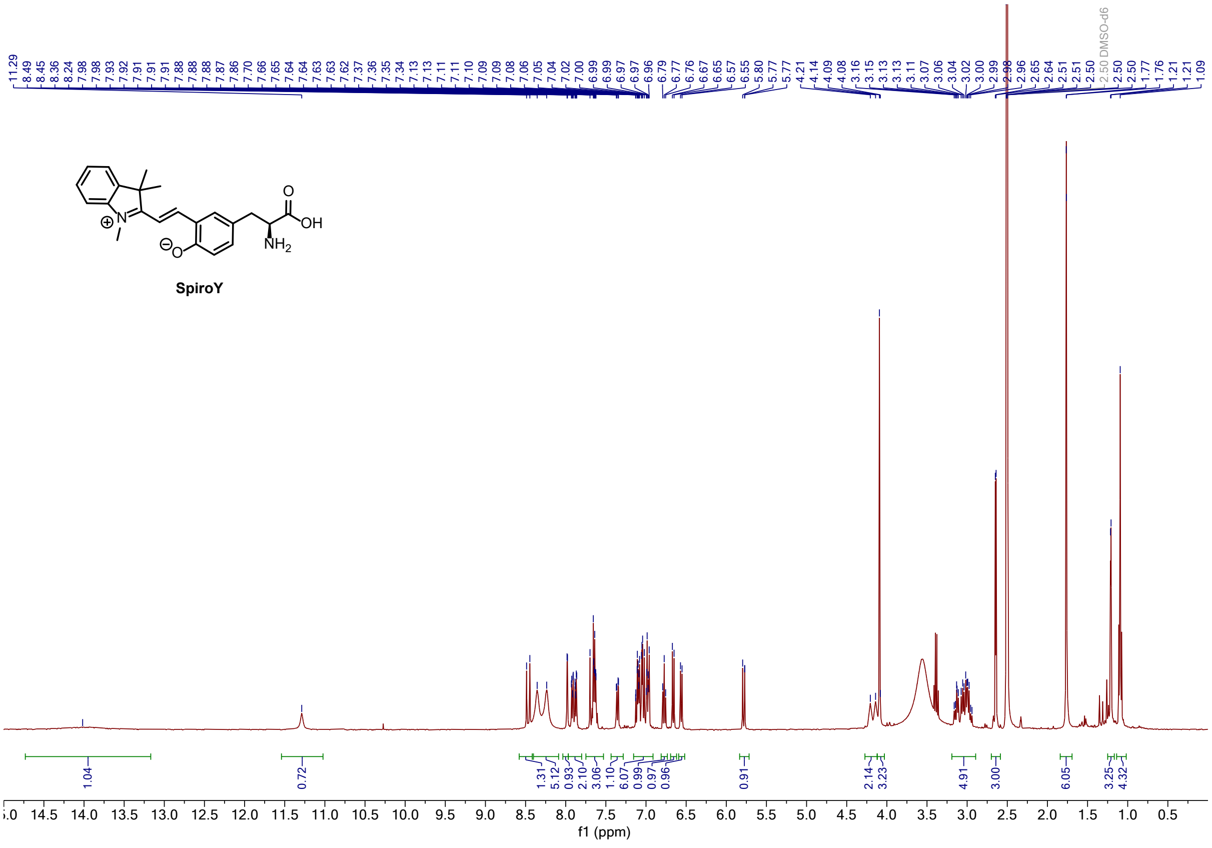
2.50 DMSO-d6
OH
SpiroY
f1 (ppm)

#### 2.1.4 13C-NMR Spectra

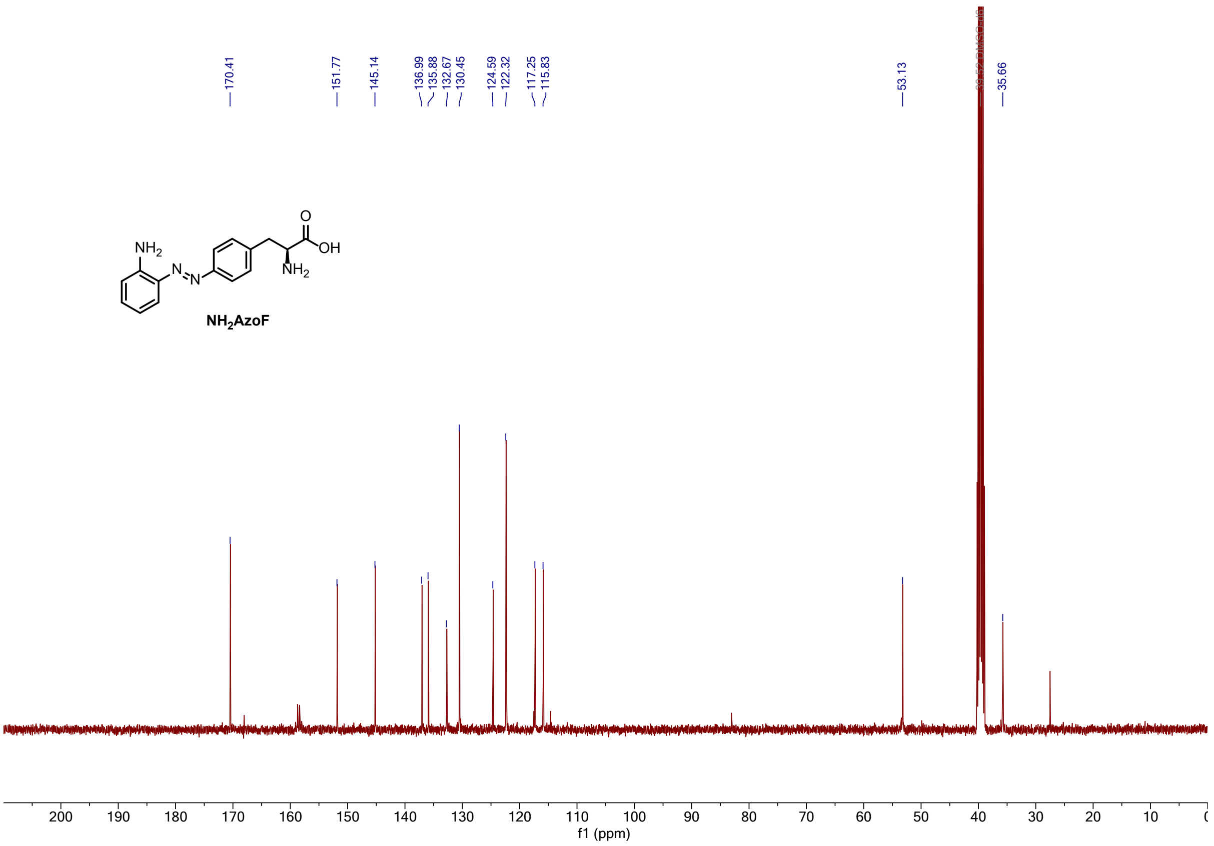

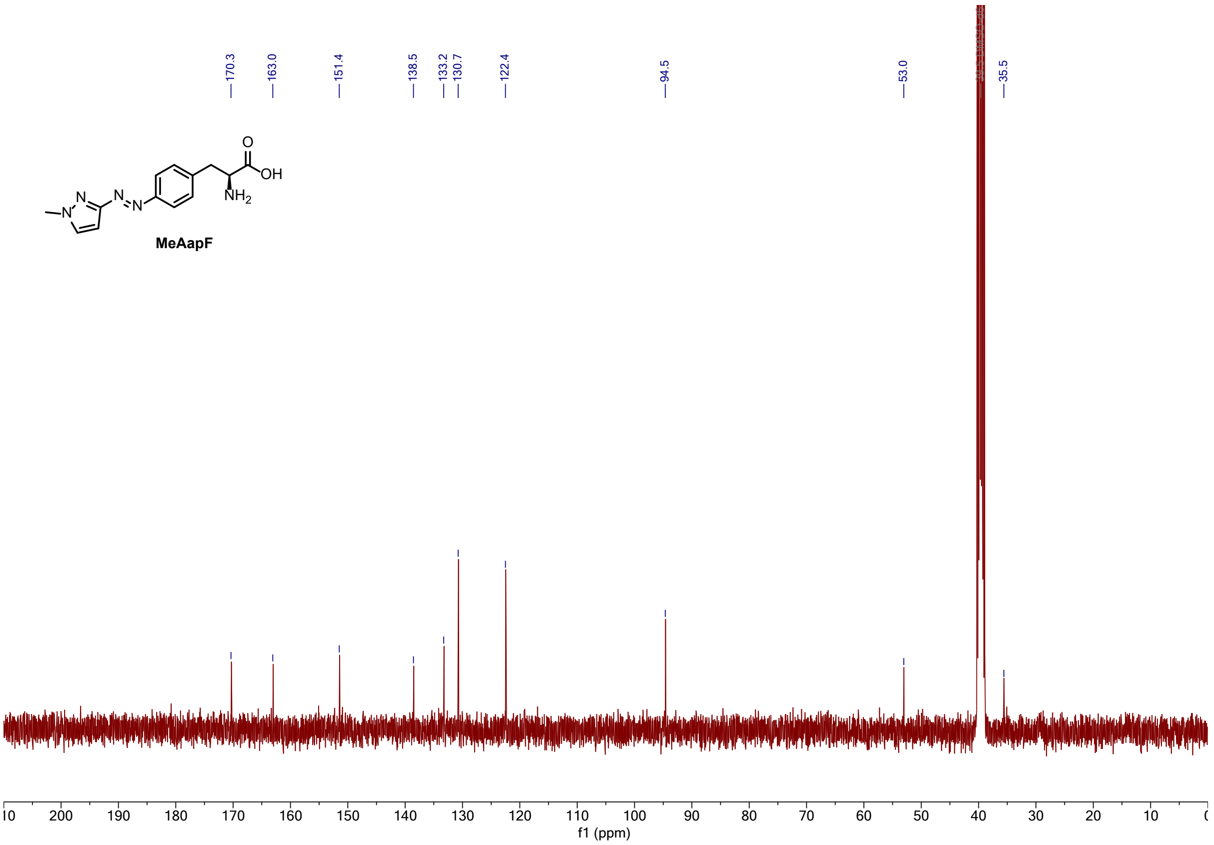

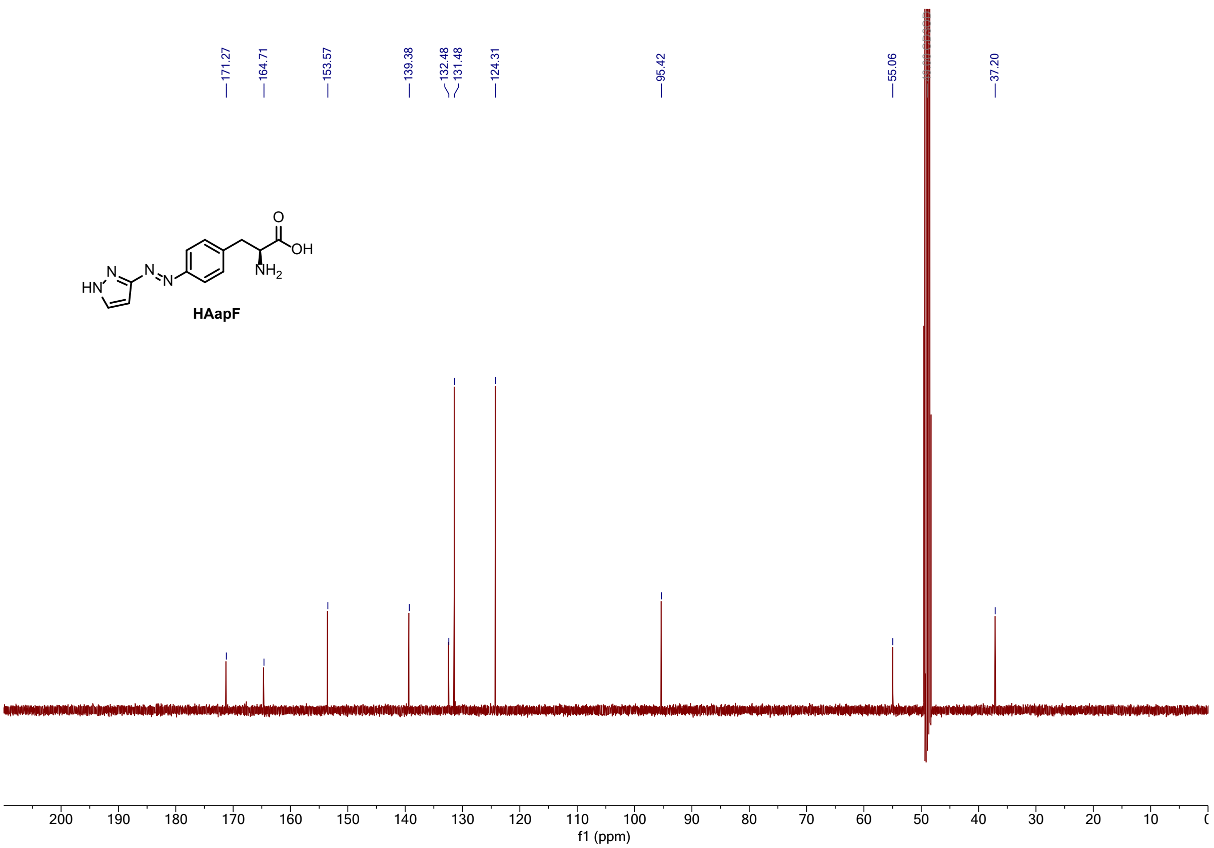
171.27
164.71
153.57
139.38
132.48
131.48
124.31
95.42
55.06
37.20
HAapF
200 190 180 170 160 150 140 130 120 110 100 90 80 70 60 50 40 30 20 10
f1 (ppm)

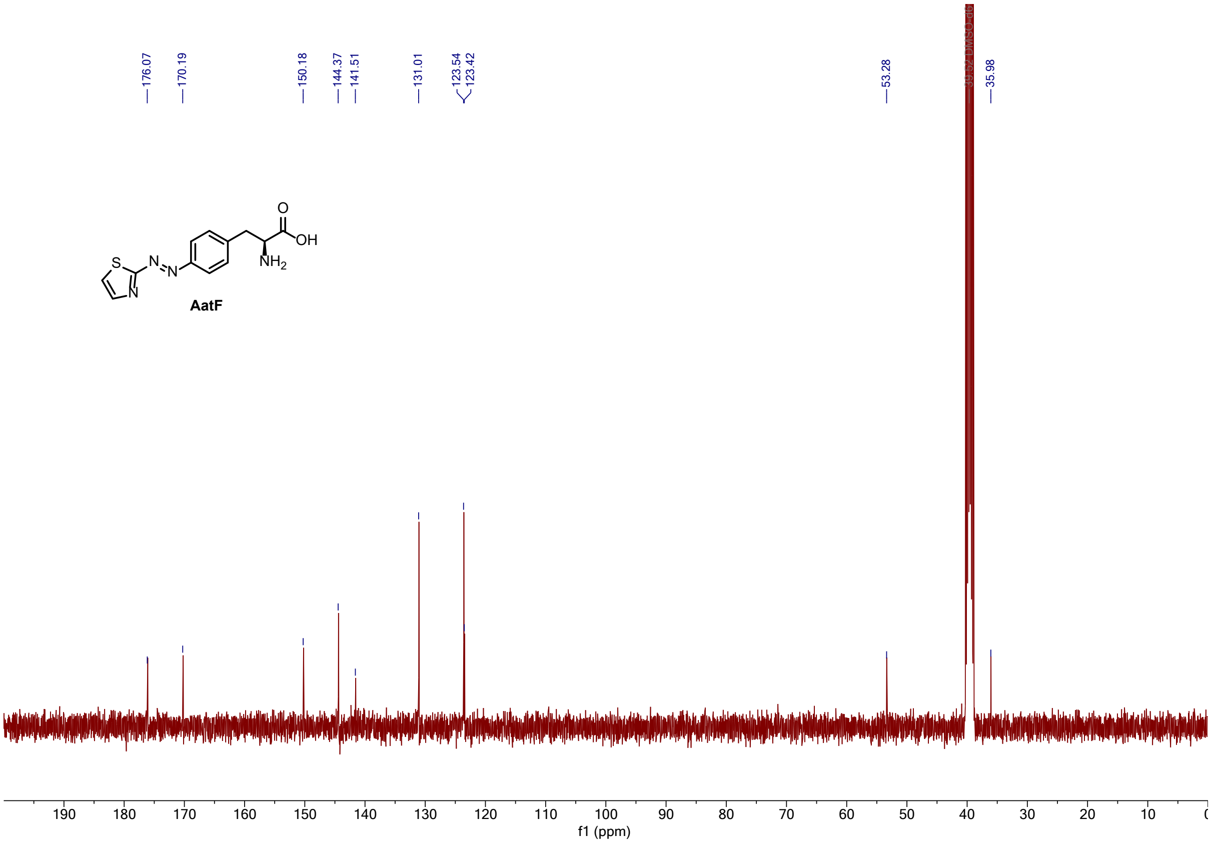
176.07
170.19
150.18
144.37
141.51
131.01
123.54
123.42
53.28
35.98
AatF
190 180 170 160 150 140 130 120 110 100 90 80 70 60 50 40 30 20 10
f1 (ppm)

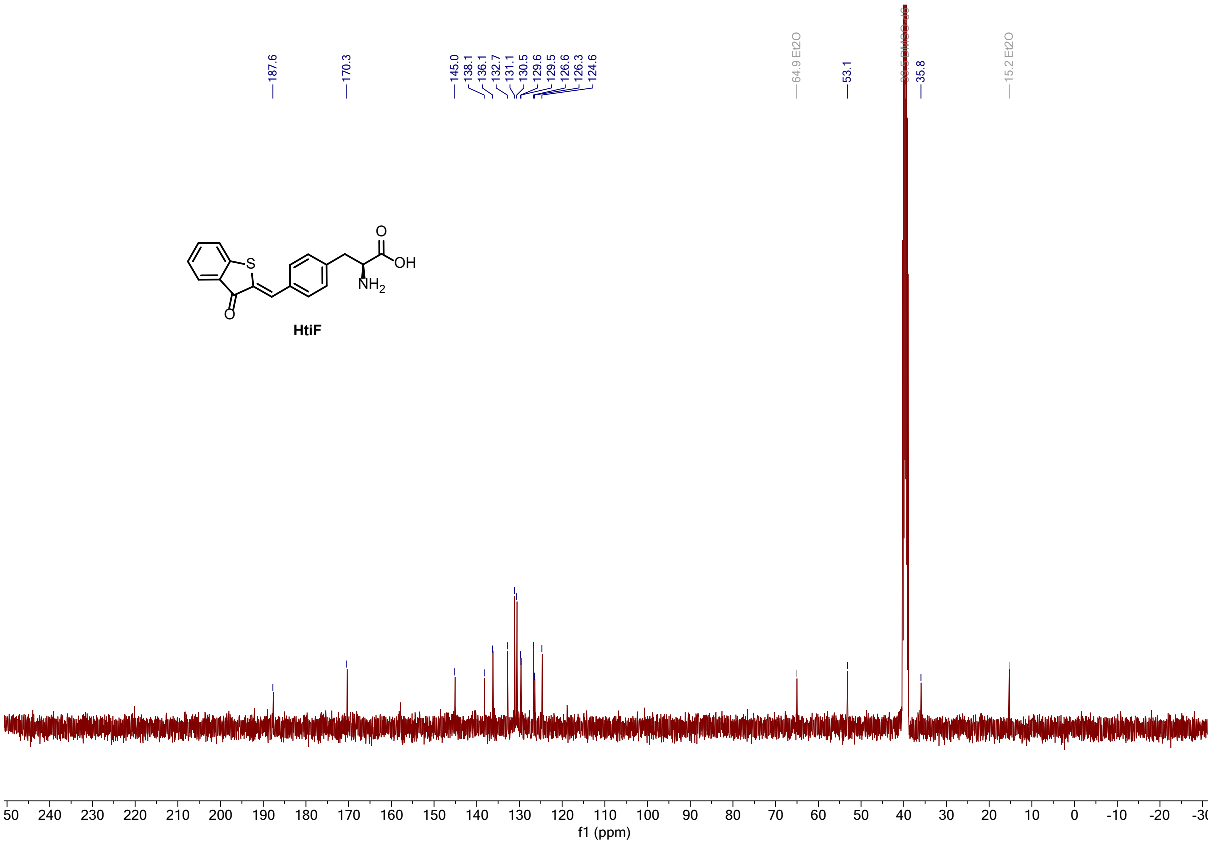
187.6
170.3
145.0
138.1
136.1
132.7
131.1
130.5
129.6
129.5
126.6
126.3
124.6
64.9 Et2O
53.1
35.8
15.2 Et2O
O
S
OH
NH2
O
HtiF
f1 (ppm)

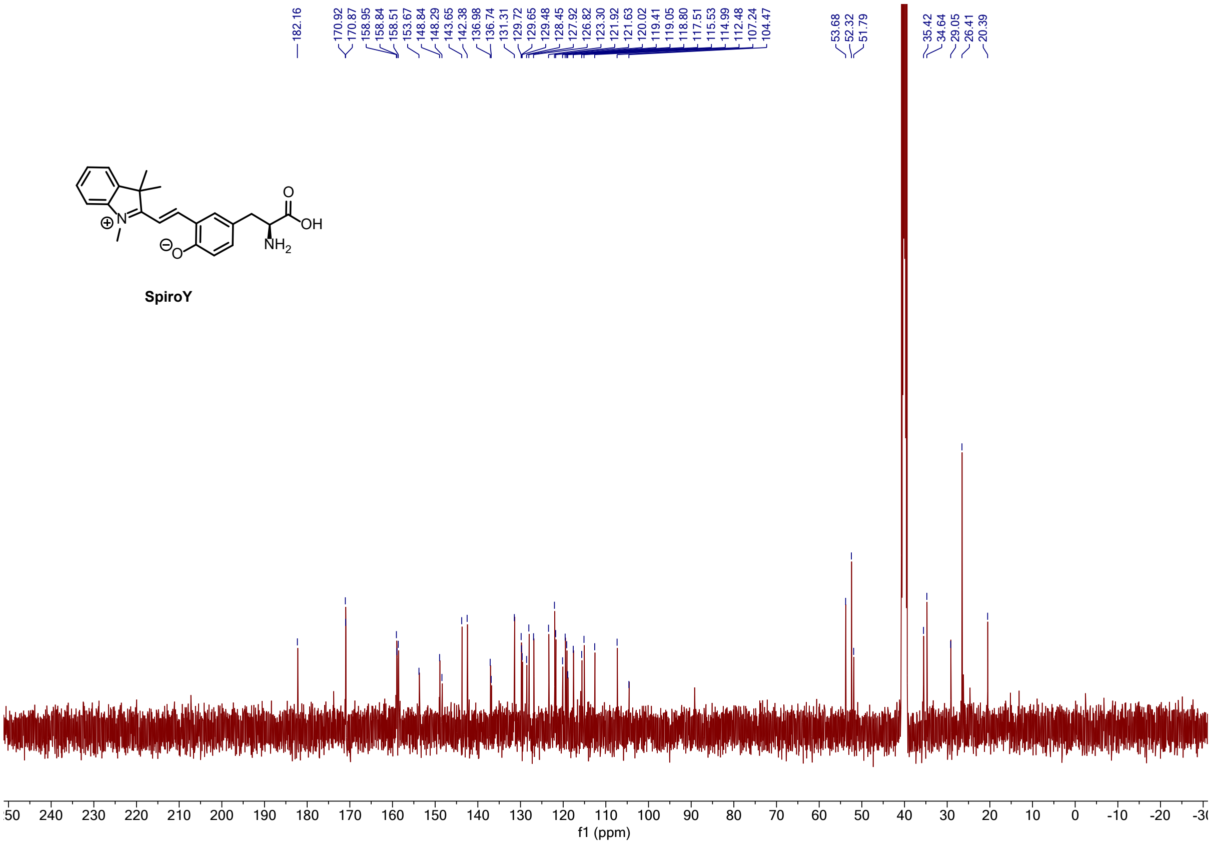
182.16
170.92
170.87
158.95
158.84
158.51
153.67
148.84
148.29
143.65
142.38
136.98
136.74
131.31
129.72
129.65
129.48
128.45
127.92
126.82
123.30
121.92
121.63
120.02
119.41
119.05
118.80
117.51
115.53
114.99
112.48
107.24
104.47
53.68
52.32
51.79
35.42
34.64
29.05
26.41
20.39
O
OH
NH2
N
O
SpiroY
f1 (ppm)

#### 2.2 Irradiation Devices

Irradiation of psUAAs and HisF-psUAA variants was conducted using a programmable power supply (Korad, KA3005P, 30 V, 5 A). The applied settings include a voltage of 3.9 V and a maximum current of 0.5 mA for each LED (**Table S14**), unless otherwise stated. To realize simultaneous irradiation and UV/Vis measurement, the LEDs were installed perpendicular to the measurement beam directly in front of the cuvette.

**Table S14.** Specifications of LEDs used for irradiation.

| LEDs | $\lambda_{max}$ [nm] | $I_{max}$ [mA][a] | $P_{max}$ [mW][b] |
|---|---|---|---|
| Seoul Viosys CUD4AF1B | 340 | 500 | 12 |
| Nichia NCSU033B | 365 | 700 | 240 |
| Nichia NCSU034B | 385 | 700 | 180 |
| Luxeon SZ-01-S2 LHUV-0395-A065 | 400 | 1000 | 155 |
| ILH-XC01-S410-SC211-WIR200 | 420 | 800 | 290 |
| Oslon SSL 80 LDCQ7P-1U3U | 455 | 1000 | 330 |
| Oslon SSL 80 LBCP7P-GYHY | 470 | 1000 | 260 |
| Winger WEPCN3-S1 | 505 | 700 | 155 |
| Oslon SSL 80 LTCP7P-KXKZ | 528 | 1000 | 120 |
| Luxeon Rebel ES Lime | 567 | 1000 | 130 |

[a]The maximum forward current was taken from the manufacturer specifications. [b]Intensity was determined with a Field $Max_{II}$-TO high sensitivity power sensor (Coherent, #1098579, Range 0–10 W) directly in front of the LED.

#### 2.3 UV/Vis absorbance spectra

UV/Vis absorbance spectra of the psUAAs and the HisF-psUAA variants were recorded in a Varian Cary 4000 Spectrometer in the range of 250–700 nm in a 1 cm quartz cuvette at 25 °C. Measurements of each isolated psUAA were performed in 50 mM Tris/HCl pH 7.5 (40 µM **AzoF**, **$NH_2$AzoF**, **MeAapF**, **HAapF**, and **AatF**, as well as 20 µM **HtiF**, and 40 µM **SpiroY**) or in DMSO (40 µM **AzoF**, **$NH_2$AzoF**, **MeAapF**, **HAapF**, and **AatF**, as well as 40 µM **HtiF**, and 40 µM **SpiroY**) and UV/Vis spectra of the HisF-psUAA variants were acquired in 50 mM Tris/HCl pH 7.5, 100 mM NaCl (28 µM K13AzoF, 13 µM K13$NH_2$AzoF, 9 µM K13MeAapF, 30 µM K13HAapF, 21 µM K13AatF, 9 µM K13HtiF). Samples were measured in the thermal equilibrium (TEQ), after establishment of a certain PSS ($PSS^{\lambda}$) or after back-isomerization to thermal equilibrium post irradiation ($TEQ^{post}$). Irradiation was maintained during the measurement to avoid back-isomerization of isomers with low thermodynamic stability. All spectra were baseline corrected at 700 nm.

#### 2.4 Wavelength screen and determination of isomerization rates

To identify the most effective wavelengths in terms of a *Z*-enriched or *E*-enriched PSS in 50 mM Tris/HCl pH 7.5 and DMSO as well as of the corresponding HisF-psUAA variants in 50 mM Tris/HCl pH 7.5, 100 mM NaCl, a wavelength screen was performed. At first, UV/Vis spectra of the thermally equilibrated state ($TEQ^{a.s.}$ or $TEQ^{a.i.}$) were recorded in the dark. Starting from this state, the absorbance at the maximum of either the π→π* or the n→π* transition was followed spectrophotometrically. At a certain timepoint, the sample was irradiated at a specific wavelength *λ* and the absorbance was followed until a plateau was reached, indicating the complete establishment of $PSS^{\lambda}$. The measurement was stopped, and a spectrum was recorded while irradiation was maintained to avoid thermal back-isomerization. Isomerization rates for *E*→*Z* isomerization were recorded starting from TEQ (isolated psUAAs) or $TEQ^{a.i.}$ (HisF-psUAA variants), while isomerization rates for *Z*→*E* isomerization were acquired starting from the $PSS^{\lambda}$ with the highest amount of *Z* isomer. Comparison of all spectra revealed the most effective wavelengths in terms of a high PSS.

For **HtiF**, the procedure to establish the best wavelengths is identical, however, the *Z* isomer is the thermodynamically favored one and consequently, the terminology *E*→*Z* and *Z*→*E* isomerization has to be reversed in the upper paragraph.

For **SpiroY**, the absorbance signal changed right after dilution of the stock solution in DMSO and thus, rates for *O*→*C* isomerization towards the newly established $TEQ^{post\ 1}$ were acquired in the dark instead of *O*→*C* photoisomerization.

Additionally, the rates of isomerization *k* were determined from the time-resolved absorbance values at the π→π* or the n→π* transition by fitting the data with a mono-exponential function (Equation 1).

$$\boldsymbol{y = y_0 + A\,(1 - e^{-x/k})}$$

**Equation 1**

The half-life of isomerization $t_{1/2}$ was then derived from the isomerization rate *k* using Equation 2.

$$\boldsymbol{t_{1/2} = ln(2) \times k}$$

**Equation 2**

For isomerization of **HtiF** with 420 nm, a double-exponential fit was required using Equation 3 and the half-life of isomerization $t_{1/2}$ was derived using Equation 4.

$$\boldsymbol{y = (y_0 D - P_D) \times e^{-k_D \times x} + P_D + y_0 A + (P_A - y_0 A) \times e^{-k_A \times x}}$$

**Equation 3**

$$\boldsymbol{t_{1/2} = \frac{ln(2)}{k}}$$

**Equation 4**

#### 2.5 Cycle performance

Cycle performance experiments of the isolated psUAAs and HisF-psUAA variants were performed to investigate their susceptibility to photobleaching. To this end, alternating isomerization between *Z*-enriched and *E*-enriched PSS$^{\lambda}$ (*O*-enriched and *C*-enriched for **SpiroY**) was induced either by irradiation or thermally for ten cycles, where switching to *E* (*O*) and subsequently to *Z* (*C*) or inversely represents one cycle. Isomerization was induced for a time period of at least ten times the $t_{1/2}$ of isomerization that was determined for each wavelength or thermally to ensure complete establishment of the respective PSS$^{\lambda}$. UV/Vis spectra of each PSS$^{\lambda}$ were measured and the absorbance at the π→π* or the n→π* transition was plotted against the number of cycles. For employed psUAA and HisF-psUAA concentrations see section 2.3.

#### 2.6 Determination of isomer ratios via high pressure liquid chromatography

To accurately determine the isomer composition of the isolated psUAAs in the established PSS, high pressure liquid chromatography (HPLC) was performed. Therefore, a reverse-phase Eclipse XDB-C18 column (Agilent, 80 Å, 5 µm, 4.6 x 150 mm) connected to an Agilent 1200 series HPLC system was employed. Mobile phase A consisted of 0.1% trifluoroacetic acid (TFA) in $H_2O$, mobile phase B of 0.1% TFA in acetonitrile. The isomers were separated at 15 °C (to prolong thermal relaxation for **$NH_2$AzoF** and **HtiF** when diluted in 50 mM Tris/HCl pH 7.5) or 25 °C (when diluted in DMSO) with a flow rate of 1 mL/min in the following manner: isocratic elution at 5% B from 0–3 minutes, linear gradient from 5–98% B from 4–23 minutes, and isocratic elution at 98% B from 24–26 minutes. To enable a quantitative comparison of the distinct isomers, their elution was followed at an isosbestic point (**Table S4**) describing a wavelength at which the absorption of both isomers is equal. Measurements of isolated psUAAs were performed in the thermal equilibrium as synthesized (TEQ), after irradiation with the selected wavelengths (PSS$^{\lambda}$) or after back-isomerization to thermal equilibrium post irradiation (TEQ$^{post}$). Isomerization was induced for a time period of at least ten times the $t_{1/2}$ of isomerization directly before the samples were subjected to HPLC. Isomer ratios of the psUAA were determined by integration of the respective isomer peaks. For HPLC measurements in both Tris buffer and DMSO, 400 µM of **AzoF**, **$NH_2$AzoF**, **MeAapF**, **HAapF**, **AatF**, as well as 400 µM **HtiF**, and 400 µM **SpiroY** were employed.

#### 2.7 Estimation of isomer ratios from UV/Vis spectra

The isomer ratios of the isolated psUAAs that are based on a diazo scaffold were also estimated as described in ref[14]. The following procedure describes the PSS estimation for a psUAA where the more thermodynamically stable isomer is the *E* isomer. The baseline-corrected spectra of the psUAA in the thermal equilibrium, as well as in the respective PSS were plotted in Origin 2022 (OriginLab). Initially, the fraction of *E* isomer present in the thermal equilibrium sample (TEQ) was assumed to be 100 %. Then, the spectrum of 100 % *Z* isomer was simulated based on the PSS with the highest concentration of *Z* isomer by estimating the residual fraction of *E* in this PSS using Equation 5.

$$A_Z = \frac{A_{PSS} - A_E \times f}{1 - f}$$

**Equation 5**

Where $A_Z$ is the absorbance of the 100 % *Z* isomer, $A_E$ is the absorbance of the 100% *E* isomer, $A_{PSS}$ is the absorbance of the photostationary state, and *f* is the fraction of *E* isomer present in the PSS. The fraction *f* was manually varied to reach a point for $A_Z$ at which the characteristic π-π* absorption signal approximates zero under the premise that the absorbance remained positive at all wavelengths (100 % *Z* spectrum). The other PSS ratios were assigned by interpolating between the *E* and *Z* isomer spectra using Equation 6.

$$f_{PSS} = \frac{A_{PSS} - A_Z}{A_E - A_Z}$$

**Equation 6**

Where $f_{PSS}$ is the fraction of *E* isomer present in the PSS and $A_Z$, $A_E$, and $A_{PSS}$ are the absorbances at the π-π* *E* isomer absorbance maximum.

Assessment of the isomer distribution based on UV/Vis spectra was not possible for **HtiF**. In the case of **SpiroY**, it was assumed that 100% of the closed form would not exhibit any characteristic absorbance at the determined $\lambda_{max}$ since the decisive conjugated π-system is disrupted in the closed form. Isomer ratios were then estimated using proportional calculations based on the “rule of three”. Estimation of the isomer ratios of the HisF-psUAA variants with a psUAA that is based on a diazo switching scaffold was equally feasible, because the protein signal at 280 nm does not interfere with the absorbance at the π→π* transition (>320 nm) of the respective psUAA.

#### 2.8 Thermal stability

Thermal stability of the psUAAs was measured in 50 mM Tris/HCl pH 7.5 at 25 °C. To this end, the PSS with the highest amount of thermodynamically less stable isomer, which is $PSS^{365}$ for **AzoF**, $PSS^{385}$ for **$NH_2$AzoF**, $PSS^{365}$ for **MeAapF**, $PSS^{340}$ for **HAapF**, $PSS^{400}$ for **AatF**, $PSS^{455}$ for **HtiF**, and $PSS^{455}$ for **SpiroY** was induced. In case of **HAapF**, a 340 nm LED was used to establish the $PSS^{340}$ which exhibits an even higher amount of *Z* isomer than the $PSS^{365}$ (**Figure S9**). For UAAs with a relatively fast thermal isomerization (**$NH_2$AzoF**, **AatF**, **HtiF**, and **SpiroY**), the regeneration of the absorbance at the $\pi \rightarrow \pi^*$ or the $n \rightarrow \pi^*$ transition after irradiation had stopped was followed over time using UV/Vis spectroscopy. For psUAAs that exhibit a longer thermal isomerization (**AzoF**, **MeAapF**, **HAapF**), HPLC measurements were conducted at certain timepoints after irradiation had stopped. The isomer ratios were deduced from the respective peak integrals and the fraction of *Z* isomer was plotted against time. Mono-exponential fitting obtained the rate constants of thermal back-isomerization *k* (Equation 1). The thermal half-life $t_{1/2}(\Delta T)$ was then derived using the isomerization rate *k* and Equation 2.

In the case of **SpiroY**, $t_{1/2}$ ($\Delta$T) was additionally measured starting from the TEQ after dilution from a 40 mM stock in DMSO to 40 µM in either DMSO or 50 mM Tris/HCl pH 7.5 ($TEQ^{post}$). The thermal half-life in buffer was sufficiently fast to be conducted via absorbance spectroscopy, while the measurement in DMSO was performed using HPLC.

#### 2.9 Computational Design of aaRS

The computational designs were created by Rosetta:MSF:NN[15] in combination with Rosetta FastDesign.[16] The design was limited to native rotamers at each position aside from positions flagged as mutable, where native amino acids were allowed. To find an aaRS suitable for **HAapF** incorporation, the following positions were allowed to mutate: 32, 65, 108, 109, 158, and 162. To improve the performance with **AzoF**, the following positions were allowed to mutate: 67, 155, 164, 167, 176, 180, and 188. In this case, the native amino acids on positions 164, 176, and 180 scored best and therefore were not mutated to another amino acid. An additional parameter file, as well as exemplary flags- and resfiles were provided for **HAapF**.

#### 2.10 Subcloning and site-directed mutagenesis

The plasmid pET28a_tmHisF_K13TAG was taken from previous work.[17] The plasmid pEVOL_AzoF-RS was provided by Peter Schultz (Scripps Research Institute, La Jolla, USA). The genes of sfGFP_wt, sfGFP_Y151TAG, AzoF-$RS_{copt}$, ONBY-RS,[10] Azo-RS 1, and AapF-RS, were codon-optimized for expression in *E. coli* and synthesized by ThermoFisher Scientific (GeneArt Strings and DNA fragments).

To introduce site-directed substitutions, deletions and insertions, or to prepare plasmids and gene fragments for Golden Gate Cloning[18] by equipping them with suitable restriction sites, PCR-based site-directed mutagenesis was employed based on a protocol from Finnzymes (ThermoFisher Scientific). Thereby, all employed primers (**Table S13**) were designed to initiate elongation in opposite directions while their 5' ends are directly adjacent to each other, with one or both primers containing the desired mutations or insertions. To introduce deletions, the primers were designed not to be adjacent to each other but flanking the region that should be deleted. The resulting linear amplicon was then re-circularized in a coupled phosphorylation-ligation reaction using T4 Polynucleotide Kinase and T4 DNA Ligase (ThermoFisher Scientific) according to the manufacturer's instructions. In all cases, the presence of mutations, insertions or deletions as well as the integrity of the plasmid was confirmed by DNA sequencing (MicroSynth AG).

Synthetic genes or amplified sequences were subcloned into plasmids using Golden Gate Cloning.[18] Golden Gate Cloning is a very efficient molecular cloning technique which allows for the simultaneous assembly of multiple DNA fragments using type IIS restriction enzymes such as BsaI and BbsI. Type IIS restriction enzymes recognize a certain palindromic site and cut the DNA outside of this recognitions site, thereby creating a four nucleotide overhang independent of the cutting site sequence. Hence, once the product is ligated, the recognition sites are no longer part of it and consequently, it cannot be cleaved again. Fragments or plasmids that contain suitable recognition sites were subjected to a one-pot restriction-ligation reaction as described in Semmelmann *et al.*.[19] To verify the integrity of all plasmids generated by Golden Gate Cloning, DNA sequencing was performed (MicroSynth AG).

In principle, the pEVOL plasmid contains two functional gene expression cassettes, one under the control of an arabinose-inducible promoter (araBAD) and one that is constitutively expressed (glnS). Using suitable primers, the two plasmids were amplified in a way that they contain either the arabinose-inducible (pBAD) or the constitutive expressing (pGLNS) cassette while simultaneously lacking the respective other one. After amplification from the pEVOL_AzoF-RS plasmid,[1] pBAD and pGLNS were equipped with suitable BsaI or BbsI recognition sites for subsequent subcloning via PCR.

All ordered synthetic genes of aaRS variants were directly subcloned into the pGLNS plasmid using Golden Gate cloning. Sequences of aaRS that have already been available in the lab (AzoF-RS$_{copt}$, pEVOL_ONBY-RS$_{i}$[20] and pEVOL_AzF-RS[21] previously obtained from Scripps Research Institute, La Jolla, USA) were equipped with suitable restriction sites by amplification from their respective plasmids and subcloned into pGLNS. Further point-mutations were introduced in the aaRS sequences using site-directed mutagenesis. For the incorporation of **H-AapF**, pEVOL_Azo-

RS 4 was generated by exchanging the two AzoF-RS genes on pEVOL_AzoF-RS[1] with Azo-RS 4 sequences in two steps of BsaI and BbsI subcloning.

The synthetic gene of sfGFP_Y151TAG was subcloned into pBAD under the control of the arabinose inducible expression cassette. Since pBAD and pGLNS carry both a chloramphenicol resistance, the whole expression module including promoter and terminator was subcloned onto a pET24a plasmid harboring a kanamycin resistance. To this end, the T7 expression cassette was removed from the pET24a plasmid and suitable restriction sites were introduced. The entire araBAD_sfGFP_Y151TAG_rrnB cassette was amplified from the pBAD plasmid with suitable restriction sites and subcloned into the prepared pET24a plasmid resulting in pETBAD_sfGFP_Y151TAG. Finally, the stop codon was reverted to the wild type tyrosine via site-directed mutagenesis.

#### 2.11 Synthetase screening

Chemically competent BL21 gold DE3 cells that already contained the pETBAD_sfGFP_Y151TAG plasmid were transformed with the respective pGLNS_aaRS plasmids. As negative and positive control, pGLNS empty vector was co-transformed with pETBAD_sfGFP_Y151TAG (–) and pETBAD_sfGFP_wt (+), respectively. The cell suspension was plated on $LB_{Cm+Kana}$ agar plates containing 30 µg/ml chloramphenicol and 75 µg/ml kanamycin and incubated at 37 °C overnight. To obtain biological triplicates, three colonies of each sample were used to inoculate 1.8 mL $LB_{Cm+Kana}$ (DWP) and cells were grown to saturation at 37 °C (>20 h). The next day, 100 µL of the pre-culture were used to inoculate 1.7 mL $2xYT_{Cm+Kana}$ (DWP). Cells were grown at 37 °C while shaking at 600 rpm and as soon as an $OD_{600}$ of >0.7 was reached, the cell suspension was transferred to a 96-well microtiter plate (MTP) with conical bottom. Gene expression was induced by addition of 0.1% L-arabinose and 0.4 mM psUAA and cultures were further incubated at 30 °C overnight. The cells were harvested by centrifugation (4000 rpm, 45 min, 20 °C) and subsequently resuspended in 150 µL 50 mM Tris/HCl pH 7.5 to reduce background fluorescence caused by the medium before 100 µL of the cell suspension were transferred to a black MTP (**Figure S14**). The sfGFP fluorescence was measured with a TECAN reader ($\lambda_{ex}$ = 488 nm, $\lambda_{em}$ = 530 nm, Z-position = 19912 µM, number of flashes = 16). To allow for a comparison of the expression levels between the different MTPs, the amplification was set to the same value for every sfGFP measurement.

To investigate whether the effect of a second aaRS copy on incorporation efficiency, chemically competent BL21 gold DE3 cells that already contained the pETBAD_sfGFP_Y151TAG plasmid were transformed with the pEVOL_aaRS-g plasmid and the experiment was conducted as described above.

#### 2.12 Computational methods

**System preparation.** The preparation of enzyme tyrosyl-tRNA synthetase variants from *Methanocaldococcus jannaschii* (MjTyr-RS) was performed using AlphaFold2 (v2.3.2)[22] (AF2) in monomer mode. The AF2 result of model 1 was selected for aminoacyl-tRNA synthetase (aaRS)-p and aaRS-q, whereas model 2 was used for aaRS-g to avoid the blockage of R163 from the active site (**Figure S16**). These predicted structures were then used to create the cluster model with the photoswitchable unnatural amino acids (psUAAs) incorporated into the active site by superposing the psUAA onto the tyrosine residue of the 1J1U PDB x-ray structure. The final cluster model for each psUAA and aaRS pair incorporates all residues and water molecules located within 6 Å of the psUAA (**Table S15**).

**Cluster model optimization.** The cluster models were optimized using the Atomic Simulation Environment (ASE)[23] Python library with the LBFGS optimizer and the AIMNET2 machine learning interatomic potential.[24] A soft harmonic constraint was applied to the peripheral anchoring carbon atoms, specifically at those where the C–C bond was cleaved to define the cluster region.[25]

**Table S15.** Cluster model composition for the selected psUAA and *Mj*Tyr-RS pairs.

| psUAAs | *Mj*Tyr-RS | cluster model composition |
|---|---|---|
| **AzoF** | aaRS-g | S30, A31, G32, I33, G34, F35, E36, P37, Q48, M52, I62, I63, I64, V65, L66, A67, D68, L69, H70, A71, Y72, N74, V103, Y104, G105, F108, Q109, M134, L136, I137, A138, R139, Y151, M154, Q155, V156, N157, G158, Y159, H160, Y161, R162, G163, V164, D165, V166, A167, V168, G169, G170, E172, Q173, I176, H177, V188, **AzoF**, WAT0, WAT1, WAT2, WAT3, WAT4, WAT5, WAT6, WAT7, WAT8, WAT9, WAT10, WAT11, WAT12, WAT13, WAT14, WAT15, WAT16, WAT17, WAT18, WAT19, WAT20, WAT21, WAT22, WAT23, WAT24, WAT25, WAT26, WAT27, WAT28, WAT29, WAT30, WAT31, WAT32, WAT33, WAT34, WAT35 |
| **$NH_2$AzoF** | aaRS-g | S30, A31, G32, I33, G34, F35, E36, P37, Q48, M52, I62, I63, I64, V65, L66, A67, D68, L69, H70, A71, Y72, N74, V103, Y104, G105, F108, Q109, Y114, M134, L136, I137, A138, R139, Y151, M154, Q155, V156, N157, G158, Y159, H160, Y161, R162, G163, V164, D165, V166, A167, V168, G169, G170, E172, Q173, I176, H177, V188, **$NH_2$AzoF**, WAT0, WAT1, WAT2, WAT3, WAT4, WAT5, WAT6, WAT7, WAT8, WAT9, WAT10, WAT11, WAT12, WAT13, WAT14, WAT15, WAT16, WAT17, WAT18, WAT19, WAT20, WAT21, WAT22, WAT23, WAT24, WAT25, WAT26, WAT27, WAT28, WAT29, WAT30, WAT31, WAT32, WAT33, WAT34, WAT35, WAT36, WAT37, WAT38 |
| **MeAapF** | aaRS-p | S30, A31, G32, I33, G34, F35, E36, P37, Q48, I62, I63, I64, E65, L66, A67, D68, L69, H70, A71, Y72, N74, V103, Y104, G105, M109, Y114, M134, L136, I137, A138, R139, Y151, M154, Q155, V156, N157, G158, I159, H160, Y161, H162, G163, V164, D165, V166, A167, V168, G169, G170, E172, Q173, I176, H177, V188, **MeAapF**, WAT0, WAT1, WAT2, WAT3, WAT4, WAT5, WAT6, WAT7, WAT8, WAT9, WAT10, WAT11, WAT12, WAT13, WAT14, WAT15, WAT16, WAT17, WAT18, WAT19, WAT20, WAT21, WAT22, WAT23, WAT24, WAT25, WAT26, WAT27, WAT28, WAT29, WAT30, WAT31, WAT32, WAT33, WAT34, WAT35 |

| | | |
|---|---|---|
| **HAapF** | aaRS-g | S30, A31, G32, I33, G34, F35, E36, P37, Q48, I63, I64, V65, L66, A67, D68, L69, H70, A71, Y72, N74, V103, Y104, G105, F108, Q109, M134, L136, I137, A138, R139, Y151, M154, Q155, V156, N157, G158, Y159, H160, Y161, R162, G163, V164, V166, A167, V168, G169, G170, E172, Q173, I176, H177, V188, **HAapF**, WAT0, WAT1, WAT2, WAT3, WAT4, WAT5, WAT6, WAT7, WAT8, WAT9, WAT10, WAT11, WAT12, WAT13, WAT14, WAT15, WAT16, WAT17, WAT18, WAT19, WAT20, WAT21, WAT22, WAT23, WAT24, WAT25, WAT26, WAT27, WAT28, WAT29, WAT30, WAT31, WAT32, WAT33, WAT34, WAT35 |
| **AatF** | aaRS-q | S30, A31, G32, I33, G34, F35, E36, P37, Q48, I63, I64, E65, L66, A67, D68, L69, H70, A71, Y72, N74, V103, Y104, G105, E109, M134, L136, I137, A138, R139, Y151, M154, Q155, V156, N157, A158, I159, H160, Y161, H162, G163, V164, V166, A167, V168, G169, G170, E172, Q173, I176, H177, V188, K204, **AatF**, WAT0, WAT1, WAT2, WAT3, WAT4, WAT5, WAT6, WAT7, WAT8, WAT9, WAT10, WAT11, WAT12, WAT13, WAT14, WAT15, WAT16, WAT17, WAT18, WAT19, WAT20, WAT21, WAT22, WAT23, WAT24, WAT25, WAT26, WAT27, WAT28, WAT29, WAT30, WAT31, WAT32, WAT33, WAT34, WAT35, WAT36, WAT37, WAT38 |
| **HtiF** | aaRS-g | S30, A31, G32, I33, G34, F35, E36, P37, H45, Q48, M52, I62, I63, I64, V65, L66, A67, D68, L69, H70, A71, Y72, N74, V103, Y104, G105, F108, Q109, M134, L136, I137, A138, R139, Y151, M154, Q155, V156, N157, G158, Y159, H160, Y161, R162, G163, V164, D165, V166, A167, V168, G169, G170, E172, Q173, I176, H177, V188, V189, C190, I191, **HtiF**, WAT0, WAT1, WAT2, WAT3, WAT4, WAT5, WAT6, WAT7, WAT8, WAT9, WAT10, WAT11, WAT12, WAT13, WAT14, WAT15, WAT16, WAT17, WAT18, WAT19, WAT20, WAT21, WAT22, WAT23, WAT24, WAT25, WAT26, WAT27, WAT28, WAT29, WAT30, WAT31, WAT32, WAT33, WAT34, WAT35 |

#### 2.13 Mass spectrometry

The HisF-psUAA variants were applied to an SDS-PAGE (1 µg) and stained with Coomassie G250 (SimplyBlue SafeStain, Lifetech). Protein bands were cut out, washed with 50 mM $NH_4HCO_3$, 50 mM $NH_4HCO_3$/acetonitrile (3/1), 50 mM $NH_4HCO_3$/acetonitrile (1/1) and subsequently lyophilized. After reduction/alkylation treatment and additional washing steps, proteins were *in gel* digested with trypsin (Trypsin Gold, mass spectrometry grade, Promega) at 37°C overnight. The resulting peptides were sequentially extracted with 50 mM $NH_4HCO_3$ and 50 mM $NH_4HCO_3$ in 50% acetonitrile. After lyophilization, peptides were reconstituted in 20 µL 1% TFA and separated by reversed-phase chromatography. An UltiMate 3000 RSLCnano System (Thermo Fisher Scientific, Dreieich) equipped with a C18 Acclaim Pepmap100 preconcentration column (100 µm i.d. x 20mm, Thermo Fisher Scientific) and an Acclaim Pepmap100 C18 nano column (75 µm i.d. x 250 mm, Thermo Fisher Scientific) was operated at flow rate of 300 nL/min and a 60 min linear gradient of 4% to 40% acetonitrile in 0.1% formic acid. The LC was online-coupled to a maXis plus UHR-QTOF System (Bruker Daltonics) via a CaptiveSpray nanoflow electrospray source. Acquisition of MS/MS spectra after CID fragmentation was performed in data-dependent mode at a resolution of 60000. The precursor scan rate was 2 Hz processing a mass range between m/z 175 and m/z 2000. A dynamic method with a fixed cycle time of 3 s was applied via the Compass 1.7 acquisition and processing software (Bruker Daltonics). Prior to database searching with Protein Scape 3.1.3 (Bruker Daltonics) connected to Mascot 2.5.1 (Matrix Science), raw data were processed in Data Analysis 4.2 (Bruker Daltonics). A customized database comprising the *T. maritima* entries from UniProt as well as manually added sequences of the mutated HisF proteins and common contaminants was used for database search with the following parameters: enzyme specificity trypsin with 2 missed cleavages allowed, precursor tolerance 10 ppm, MS/MS tolerance 0.04 Da. As general variable modifications, deamidation (0.984 Da) of asparagine and glutamine, oxidation of methionine 15.995 Da, carbamidomethylation of cysteine 57.021 Da, or propionamide modification of cysteine 71.037 Da were set. Specific variable modifications for identification of unnatural amino acids were as follows: **AzoF**: 104.037 Da **$NH_2$AzoF**: 119.048 Da, **MeAapF**: 108.044 Da, **HAapF**: 94.028 Da, **AthzF**: 110.989 Da, or **HtiF**: 159.998 Da. Spectra of peptides containing unnatural amino acids were inspected manually.

#### 2.14 Expression of HisF-psUAA variants in midi-scale

For expression of HisF-psUAA proteins, cells containing pET28a_HisF_K13TAG were transformed with the plasmids containing the appropriate aaRS variants for incorporation of **AzoF**, **$NH_2$AzoF**, **MeAapF**, **HAapF**, **AatF**, and **HtiF** (**Table S9**). A 50 mL $LB_{Cm+Kana}$ culture was inoculated and grown at 37 °C overnight. The next day, 50 mL $TB_{Cm+Kana}$ were inoculated to an $OD_{600}$ of 0.1 and incubated at 37 °C until an $OD_{600}$ >1.2 was reached. Gene expression was induced by addition of 0.4 mM psUAA and 0.5 mM IPTG (+0.1 % L-arabinose if the aaRS was encoded on the pEVOL plasmid) before the culture was incubated at 30 °C overnight. The cells were harvested by centrifugation (4000 rpm, 45 min, 20 °C) and subsequently resuspended in 5 mL 50 mM Tris/HCl pH 7.5, 100 mM NaCl, 10 mM imidazole. Cells were lysed by sonification (2 min, 30%, 2 s pulse, 2 pause, followed by a heat step at 65 °C and another centrifugation step to remove *E. coli* proteins. The lysate was subjected to immobilized metal ion affinity chromatography using HisSpinTrap™ columns (GE Healthcare) following to protocol provided by the manufacturer and elution was performed twice with 250 µL 50 mM Tris/HCl pH 7.5, 100 mM NaCl, 750 mM imidazole. The buffer of the eluates was exchanged using NAP-5 columns (GE Healthcare) according to the manufacturer's instructions. Concentration of the HisF-psUAA variants was determined with the Bradford Assay before their photochemical characterization was performed as described in sections above. For long-term storage, the proteins were dropped into liquid nitrogen and frozen at –80 °C.

#### 2.15 Heterologous expression of HisA

HisA from *T. maritima* was produced by heterologous gene expression in *E. coli* BL21 Gold (DE3) cells (Agilent Technologies) using pET21_HisA.[26] The crude extract was subjected to a heat step (15 min at 73 °C) and the recombinant protein was purified from the soluble fraction by nickel-affinity chromatography (HisTrap FF crude 5 ml; GE Healthcare). The column was equilibrated with 50 mM potassium phosphate, 300 mM NaCl, and 1 mM imidazole, pH 7.5 and bound protein as eluted by applying a linear gradient to 300 mM imidazole. Fractions containing pure protein were pooled and dialyzed against 50 mM potassium phosphate, pH 7.5. Based on SDS-PAGE analysis, the purity of all samples was at least 90%. The proteins were dripped into liquid nitrogen and stored at –80 °C.

#### 2.16 Biochemical synthesis of ProFAR

ProFAR was synthesized as described elsewhere,[27] starting from 5-phospho-D-ribosyl α-1-pyrophosphate and adenosine triphosphate in 50 mM $NH_4COOCH_3$ pH 7.8. The reaction progress was tracked spectrophotometrically and products were purified with ion-exchange chromatography using a POROS column (HQ 20, 10 mL, Applied Biosystems) and a linear gradient of $NH_4COOCH_3$ (50 mM / 1 M). ProFAR concentration was determined at 300 nm ($\varepsilon_{300}$ = 6069 $M^{-1}cm^{-1}$)[28] and ProFAR purity was assessed from the absorbance ratio $A_{290}/A_{260}$ for each fraction. Thereby, highly concentrated and > 90% pure ($A_{290}/A_{260}$ = 1.1–1.2 accounts for >95% purity)[29] fractions were lyophilized and stored at –80 °C. ProFAR identity was confirmed using the HisF assay described below.

#### 2.17 Activity measurements

The HisF activity was measured continuously following PrFAR turnover at 300 nm [$\varepsilon_{300}$(PrFAR-AICAR) = 5637 $M^{-1}cm^{-1}$].[30] Reaction conditions included 50 mM Tris acetate pH 8.5, 100 mM ammonium acetate, 1 µM HisA (to convert ProFAR to PrFAR) and 7 µM ProFAR in a total volume of 250 µL unless otherwise stated. ProFAR and HisA were synthesized as previously reported.[17]

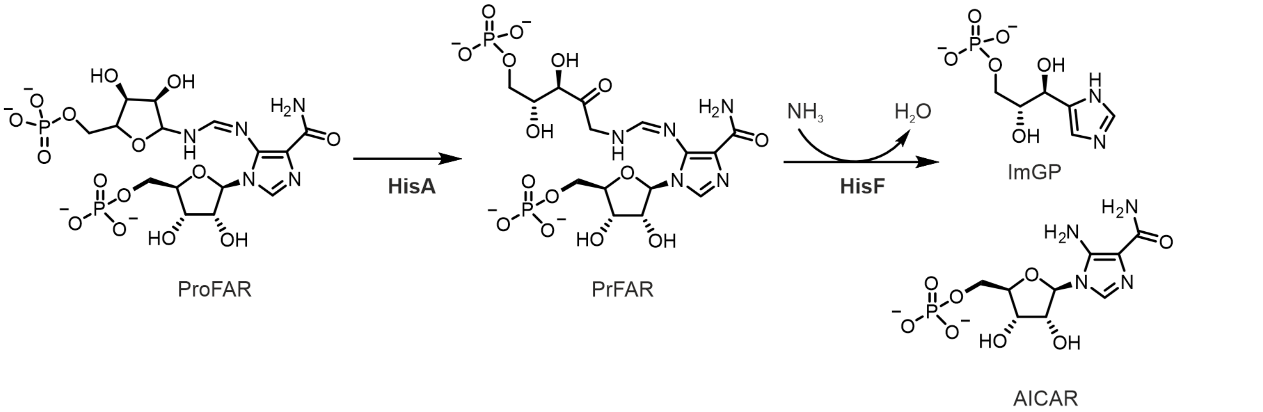

**Scheme S7.** Reaction assay for the determination of HisF activity. The metastable substrate PrFAR is in situ generated from the substrate analogue ProFAR using HisA. The turnover of PrFAR to ImGP and AICAR catalyzed by HisF can be monitored spectrophotometrically.

Measurements were performed in a 1 cm quartz cuvette at 25 C° The reaction was started with 50 nM HisF-wt, 33 nM K13AzoF, 50 nM K13$NH_2$AzoF, 100 nM K13MeAapF, 70 nM K13HAapF, 50 nM K13AatF, or 70 nM K13HtiF. The proteins were either used in the thermal equilibrium as isolated ($TEQ^{a.i.}$), after a certain $PSS^{\lambda}$ has been established or after back-isomerization to thermal equilibrium post irradiation ($TEQ^{post}$). To this end, irradiation with the appropriate wavelengths or incubation at 25 °C was conducted for a time period of at least ten times the $t_{1/2}$ of isomerization that was determined for each wavelength or thermally (**Table S11**) and was maintained over the whole measurement. To verify that the irradiation conditions are principally not harmful for the enzyme, HisF-wt was subjected to the same irradiation conditions as the variants. Measurements were performed in technical triplicates. The initial slopes were determined in a global linear fit using a concatenated linear regression including a direct weighting to consider the S.E., which is indicated as error bar. The slopes *m'* ± SE were then used to calculate the initial velocities $v/E_0$ ± SE ($s^{-1}$) using the Lambert Beer Equation, followed by normalization with the enzyme concentration. The LRF reflects the ratio of activities between TEQ and $PSS^{\lambda 1}$ or $PSS^{\lambda 1}$ and $PSS^{\lambda 2}$. The respective LRF values were determined by selecting the data sets of both states and fitting them with a global fit analysis using a linear regression interaction model (Equation 7) and direct weighting,

$$\boldsymbol{v = m' \cdot D \cdot t}$$

**Equation 7**

with *A* = absorbance and *t* = time. The slope *m'* was shared in the global fit and the dummy constant *D* of the slower reaction was fixed to 1. The fitted *D* ± S.E. value of the faster reaction then corresponded to the LRF ± S.E. as defined by Equation 8.

$$\boldsymbol{LRF = \frac{v_1}{v_2}} \qquad v_1 > v_2$$

**Equation 8**

The p-value of the fit thereby indicates the significance of the LRF and the S.E..
